# Menaquinone depletion resensitises bedaquiline-resistant tuberculosis

**DOI:** 10.64898/2026.02.10.702511

**Authors:** Jennefer Wetzel, John Dallow, Ellie Davis, William H. Pearson, Stijn Daems, Matthias Govaerts, Joyce Hereijgers, Joke Sprangers, Barry Truebody, Valerie Maes, Vera van Hasselt, Annelies Leemans, Venugopal Pujari, Ann Vos, Carlos M. Martínez Viturro, José Enrique Gómez, Mirte Peeters, Michelle Gerber, Nandini Chhabra, Annelies Wouters, Melissa Everaerts, Hannah Painter, Raisha Fathima, Sam J. Willcocks, Cadi Davies, Valerie Raeymaekers, Taane G. Clark, Ruxandra Draghia-Akli, Helen Fletcher, Marnix Van Loock, Martin L. Hibberd, Serge Mostowy, Dean C. Crick, Alexander S. Pym, Kirandeep Samby, Paul Jackson, Andrés A. Trabanco, Gerald Larrouy-Maumus, Adrie J.C. Steyn, Bart Stoops, Neeraj Dhar, Clara Aguilar-Pérez, Dirk A. Lamprecht, Richard J. Wall, Anil Koul

**Affiliations:** Johnson & Johnson, Beerse, Belgium; Department of Infection Biology, Faculty of Infectious and Tropical Disease, London School of Hygiene and Tropical Medicine, London, WC1E 7HT, UK; Charles River Laboratories, Turnhoutseweg 30, 2340, Beerse, Belgium; Africa Health Research Institute, University of KwaZulu Natal, Durban, South Africa; VIB-KU Leuven Center for Microbiology, 3001, Heverlee, Belgium; Microbiology, Immunology, and Pathology Department, Colorado State University, Fort Collins, CO, 80523, USA; Global Discovery Chemistry, Janssen-Cilag SA, a Johnson & Johnson company, C. Río Jarama, 75A, 45007 Toledo, Spain; Vaccine and Infectious Disease Organization, University of Saskatchewan, Saskatoon, Saskatchewan, S7N 5E3, Canada; Johnson & Johnson, Spring House, Pennsylvania, USA; Johnson & Johnson, High Wycombe, UK; Johnson & Johnson Pvt. Ltd, Gurugram 122003, India; Johnson & Johnson, La Jolla, California, USA; Centre for Bacterial Resistance Biology, Department of Life Sciences, Faculty of Natural Sciences, Imperial College London, London, SW7 2AZ, UK; Department of Microbiology, University of Alabama at Birmingham, Birmingham, AL, USA; Centers for AIDS Research and Free Radical Biology, University of Alabama at Birmingham, Birmingham, AL, USA

## Abstract

Tuberculosis remains a leading cause of global mortality, and rising bedaquiline resistance threatens the effectiveness of current drug-resistant treatment regimens. Bedaquiline resistance typically arises through mutations in *Rv0678* that upregulate drug efflux and confer cross-resistance to multiple drug classes. Here, we identify and optimise a chemical series targeting MenG, a central enzyme in the menaquinone biosynthesis pathway, yielding potent bactericidal inhibitors with *in vivo* efficacy. Strikingly, MenG inhibition restored bedaquiline susceptibility in efflux-mediated resistant strains, an effect confirmed *in vivo* where combination therapy achieved a 99.8% reduction in bacterial burden compared with bedaquiline alone. Potentiation also extended to pretomanid and other key agents. Disruption of upstream menaquinone and shikimate pathway enzymes produced similar resensitisation, establishing these pathways as tractable targets for restoring drug susceptibility in *Mycobacterium tuberculosis*. These findings provide a novel strategy to overcome bedaquiline resistance and strengthen future regimens for efflux-mediated drug-resistant tuberculosis.

## Main text

Tuberculosis (TB) remains one of the leading causes of death from an infectious disease, responsible for an estimated 1.23 million deaths in 2024 (*1*), and global control efforts are increasingly threatened by the rise of drug-resistant TB (DR-TB). The introduction of bedaquiline marked a significant advance in TB therapy as the first drug with a novel mechanism of action in decades (*2*). Incorporating bedaquiline into DR-TB treatment regimens enabled significant treatment shortening, exemplified by the six-month all-oral BPaL regimen (bedaquiline, pretomanid and linezolid) (*3, 4*). These regimens have transformed DR-TB care, delivering the highest treatment success rates seen in decades, reaching 71% in recent reports (*5*). However, clinical resistance to bedaquiline is emerging (*6*), with rates exceeding 14% in some countries (*7*), threatening to erode the gains achieved in DR-TB management and highlighting the urgent need for strategies that preserve or restore its efficacy.

Bedaquiline inhibits the ATP synthase *c*-subunit (AtpE) of *Mycobacterium tuberculosis*, the causative agent of TB, impairing its energy metabolism and depleting intracellular ATP (*8-10*). Resistance arises predominantly through mutations disrupting *Rv0678* (MmpR5), which encodes the repressor of the MmpS5–MmpL5 efflux pump, resulting in increased drug efflux (*11*). Bedaquiline’s long terminal half-life generates prolonged periods of subtherapeutic exposure, potentially favouring selection of such efflux-mediated mutants. Importantly, clinical *Rv0678*-mediated resistance cannot be completely overcome by dose escalation (*12, 13*) and confers cross-resistance to other anti-TB agents, including clofazimine and macozinone (*14*). Although the full clinical impact remains to be defined, *Rv0678* variants are increasingly reported worldwide, including lineages predating the clinical introduction of bedaquiline (*15*).

Here, we sought to identify and optimise a compound series capable of preventing or overcoming bedaquiline resistance. This effort led to the discovery of two lead compounds, JNJ-6887 and JNJ-1866, the latter exhibiting promising efficacy in a mouse infection model. Mechanistic studies identified MenG, a key enzyme in the menaquinone biosynthesis pathway, as the molecular target. In *M. tuberculosis*, menaquinone-9 (MK-9) mediates electron transfer along the electron transport chain (ETC) and is essential for oxidative phosphorylation during both active and latent infection (*16-18*). Building on this discovery, we demonstrate that dual inhibition of MenG and ATP synthase enhances bactericidal activity and restores bedaquiline susceptibility in a clinically relevant efflux-mediated mutant. Importantly, this resensitisation was confirmed *in vivo* in a mouse model of infection. Potentiation also extended to other compounds, including pretomanid, suggesting a broader opportunity for combination strategies.

The upstream shikimate pathway provides the precursors required for menaquinone biosynthesis (*19*), functionally linking the two metabolic pathways. Inhibition of other enzymes in the menaquinone and shikimate pathways produced identical resensitisation effects in an efflux-based resistant strain. Similar to ethionamide boosters (*20, 21*), these findings reveal a novel strategy to restore drug susceptibility, but with the added advantage of intrinsic bactericidal activity and broad potentiation of other TB drugs. Together, these findings establish MenG and the wider shikimate-menaquinone superpathway as tractable resistance-breaking targets and demonstrate that metabolic co-targeting offers a powerful strategy to counter emerging efflux-mediated resistance and strengthen future DR-TB treatment regimens.

## Results

### Hit-to-lead optimisation of GSK517A

To identify new inhibitors with activity against DR-TB, we focused on GSK517A (hereafter JNJ-8833; **Fig. 1a**), a chemically tractable hit compound previously identified in a high-throughput screen (*22*) and selected for structure-activity relationship (SAR)-guided optimisation to improve its activity and chemical characteristics. Medicinal chemistry optimisation of JNJ-8833 produced a bactericidal analogue, JNJ-6887, with a 25-fold improvement in potency (**Fig. 1a-b**). Bactericidal activity was further confirmed by single-cell microscopy, which enables direct observation of individual bacterial fates and a clearer distinction between bacteriostatic and bactericidal effects (*23, 24*) (**Fig. 1c; Fig. S1; Supplementary Video 1**).

**Fig. 1.**
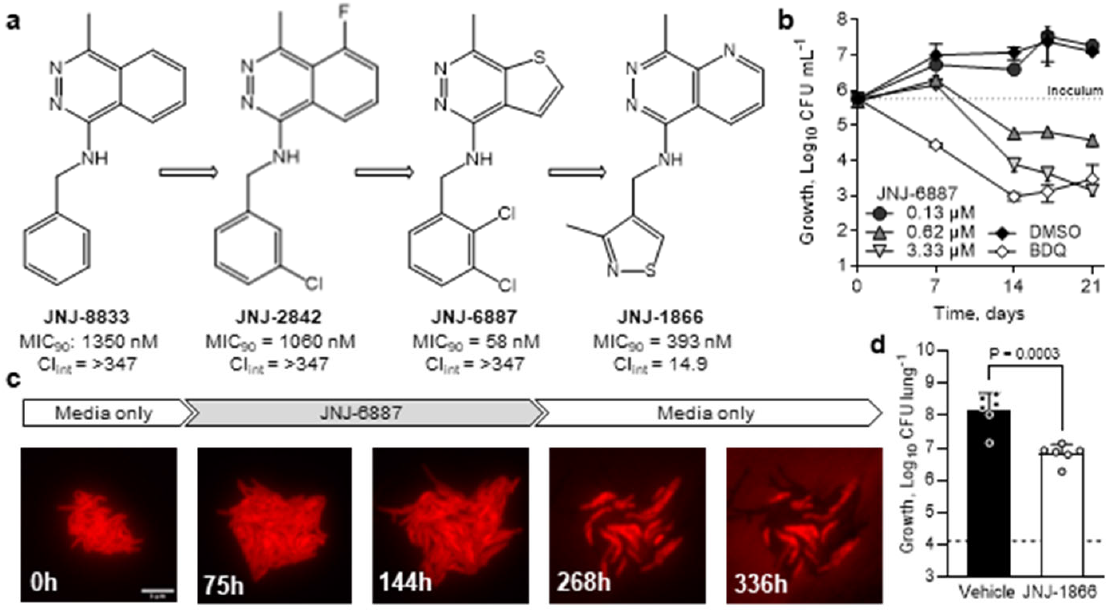
Hit-to-lead optimisation identifies potent, metabolically stable analogues. **a**, Compound optimisation from GSK517A (JNJ-8833) (*22*) leading to JNJ-6887 (∼25-fold increased potency) and JNJ-1866, which shows improved metabolic stability. MIC_90_ = minimum inhibitory concentration, 90% inhibition; Cl_int_ = mouse microsomal clearance (in µL min^-1^ mg^-1^; **Table S1**). **b**, Time-kill kinetics assay based on colony-forming units (CFU) with 0.13–3.33 µM (2–60x MIC_90_) JNJ-6887 compared with DMSO control and 0.5 µM (5x MIC_90_) bedaquiline (BDQ). n = 3 biological replicates. **c**, Single-cell imaging showing bactericidal activity of JNJ-6887. Representative image series of *M. tuberculosis* expressing *Td*Tomato grown in a microfluidic device and exposed to 1 µM JNJ-6887 between 76 and 267 h. Scale bar, 5 µm. See also **Fig. S1**; **Movie S1. d**, *In vivo* efficacy of JNJ-1866 in mice treated for 12 days (twice daily oral administration; 150 mg kg^-1^; white) compared with untreated (vehicle; black) at day 21 post-infection. Dotted line indicates CFU at treatment start. n = 6 mice. Significance was calculated with a two-sided unpaired *t*-test. For panels **b** and **d**, data shown are mean ± s.d.

However, JNJ-6887 showed poor metabolic stability in liver microsomes, predicting poor exposure and rapid clearance. Subsequent compound optimisation efforts therefore focused on improving metabolic stability while retaining bactericidal activity, leading to the synthesis of JNJ-1866, which exhibited an 8-fold reduction in potency compared with JNJ-6887 but showed markedly improved metabolic stability (**Fig. 1a; Table S1-2**). *In vitro* absorption, distribution, metabolism and excretion (ADME) and *in vitro* safety profiling, including mitochondrial toxicity, revealed an acceptable safety and physicochemical profile for JNJ-1866 **(Table S3**), supporting progression to *in vivo* pharmacokinetic (PK) evaluation. In mouse PK studies, JNJ-1866 demonstrated superior exposure and bioavailability compared with JNJ-6887 (**Fig. S2**; **Table S4-5**). On this basis, JNJ-1866 was advanced to efficacy testing in an acute mouse model of infection, where mice are infected for 7 days with *M. tuberculosis* H37Rv before treatment starts (**Fig. S3)**. Oral administration of 150 mg kg^-1^ JNJ-1866 for 12 days resulted in a significant 1.35-log reduction in lung bacterial burden relative to the end-of-study vehicle control, demonstrating *in vivo* proof-of-principle (**Fig. 1d**). These findings provided a strong foundation for further exploration of this chemical series.

### Inhibition of menaquinone biosynthesis

To identify the molecular target of this compound series, we selected *M. tuberculosis* mutants resistant to JNJ-6887 and JNJ-1866. The isolated independent colonies conferred 26-133-fold resistance and were cross-resistant to other series analogues (**Table S6**), while retaining susceptibility to first- and second-line TB drugs and late-stage clinical candidates, indicating a possible novel mode of action or resistance (**Table S6**). JNJ-6887 and JNJ-1866 both retained activity across drug-sensitive and -resistant clinical isolates (*25*), comparable to their activity against the laboratory-adapted H37Rv (**Table S7**).

Whole genome sequencing revealed single-nucleotide polymorphisms in *Rv0558*, which encodes the membrane-associated methyltransferase MenG (also annotated as UbiE/MenH; **Table S6**). MenG is an S-adenosylmethionine (SAM)-dependent methyltransferase that converts demethylmenaquinone-9 (DMK-9) to MK-9, a reaction essential for *M. tuberculosis* growth (**Fig. S4**) (*26-28*). Additional resistance-selection experiments with series analogues identified further mutations. Most mutations resided within or adjacent to the predicted UbiE/COQ5 (Coenzyme Q5) domains but did not impair growth (**Fig. S5**). The role of these mutations in resistance was validated using a recombinant strain expressing an additional *menG* copy harbouring a C146R substitution, which conferred resistance to MenG inhibitors while retaining susceptibility to a reference compound (**Fig. 2a**; **Fig. S6**). Analysis of a large clinical isolate diversity database (*29*) revealed a limited presence of MenG-based resistance-conferring mutations; only one isolate (<0.002% of total) carried the D25G variant (**Fig. S5**), suggesting that such mutations are rare but tolerated in clinical populations.

**Fig. 2.**
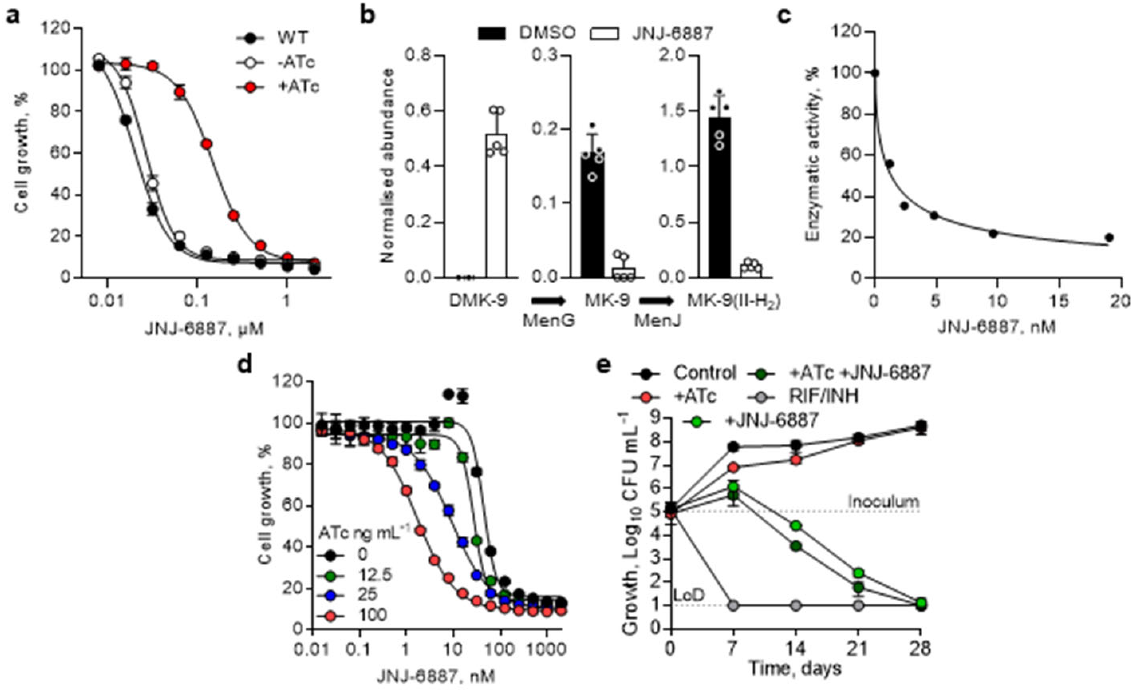
Target deconvolution identifies menaquinone biosynthesis enzyme, MenG. **a**, Dose-response curves for JNJ-6887 against WT *M. tuberculosis* (black), *M. tuberculosis* carrying a chromosomally integrated, inducible MenG-C146R expression plasmid (WT::C146R) with (red) and without (white) 500 ng mL^-1^ anhydrotetracycline (ATc) induction. n = 2 technical replicates. Representative of two independent experiments. **b**, Measurement of menaquinone levels in culture after 4 days in the presence (white) or absence (black) of 3.33 μM JNJ-6887, showing an increase in DMK-9 (MenG substrate) and a decrease in MK-9 (MenG product) and downstream MK-9(II-H_2_). n = 5 technical replicates. **c**, Direct inhibition of membrane-isolated MenG protein with JNJ-6887 in a dose-dependent manner. IC_50_ = 1.1 nM. Representative figure from three independent experiments. **d**, Dose-response curves of JNJ-6887 following CRISPRi-mediated “low” *menG* knockdown showing >20-fold increased inhibitor susceptibility upon induction. MenG knockdown was achieved in an ATc-dependent manner (0-100 ng mL^-1^). n = 3 technical replicates. Representative of two independent experiments shown. **e**, Bactericidal activity of a combination of CRISPRi-mediated “low” MenG knockdown with 100 ng mL^-1^ ATc and 3.33 µM JNJ-6887 over 28 days. Rifampicin (RIF; 14.58 µM) and isoniazid (INH; 5.8 µM) (grey) used as positive kill control. LoD: limit of detection. n = 3 biological replicates. For panels **a, b, d, e**, data shown are mean ± s.d.

In the absence of an *M. tuberculosis* MenG crystal structure, we generated an AlphaFold model and docked the SAM substrate based on the SAM-bound *S. cerevisiae* COQ5 structure (*30-32*). This revealed a putative ligand-binding pocket adjacent to resistance-conferring mutations (D25, N28, S32), with compounds likely obstructing DMK-9 binding (**Fig. S7**). Predicted interactions included hydrogen bonds between the hydrazine and R121, hydrophobic binding of the bicyclic ring with F17 and L185, and *pi-pi* stacking of the phenyl or isothiazole ring with F148 and W39. Reduced conformational flexibility of aromatic substituents (*33*) may explain the increased potency and activity shifts seen with JNJ-6887 and JNJ-1866 (**Fig. S7**; **Table S6**).

To evaluate target inhibition in a whole-cell context, we measured the abundance of menaquinone species after 4 days of JNJ-6887 exposure. Treatment resulted in an accumulation of the MenG substrate, DMK-9, and a corresponding decrease in MK-9 and β- dihydromenaquinone-9 (MK-9(II-H_2_)), the principal form of menaquinone in mycobacteria (*34*), consistent with MenG inhibition (**Fig. 2b; Fig. S4, 8**). Humans lack the menaquinone biosynthetic pathway, but they do obtain MK-4 (vitamin K_2_) from the diet for unrelated physiological functions. We therefore assessed whether exogenous MK-4 could restore bacterial growth. JNJ-6887 retained full bactericidal activity, even at MK-4 concentrations >50-fold above plasma levels (*35*), indicating limited potential for host rescue (**Fig. S9**).

Direct target engagement was confirmed by measuring radiolabelled methylation of DMK-8 to MK-8 in MenG-containing *M. tuberculosis* membrane preparations (*36*). We observed dose-dependent inhibition of enzymatic activity by JNJ-6887 (IC_50_ = 1.3 ± 0.4 nM) and JNJ-1866 (IC_50_ = 12 ± 2.8 nM), correlating with whole cell potency (**Fig. 2c**). Using a validated CRISPRi system (*26*), a *menG* “low” transcript knockdown sensitised *M. tuberculosis* >20-fold to JNJ-6887 in an ATc-dependent manner, further confirming on-target activity (**Fig. 2d; Fig. S10**). Finally, given mixed reports regarding the essentiality of the menaquinone pathway and its apparent low vulnerability in genome-wide essentiality screens (*26, 37*), we investigated whether sustained inhibition could achieve killing below the limit of detection. A time-course assay with 3.33 µM (60x MIC_90_) JNJ-6887 resulted in bactericidal activity below the limit of detection (**Fig. 2e**). Notably, combining treatment with CRISPRi-mediated *menG* “low” knockdown did not markedly accelerate killing.

### Resensitisation of bedaquiline resistance

Inhibitors of menaquinone biosynthesis have previously been shown to synergise with bedaquiline (*38, 39*). Checkerboard assays confirmed synergy between JNJ-6887 and bedaquiline (**Fig. S11**). At 1-2× MIC_90_ concentrations, monotherapies produced minimal reductions in CFU counts, whereas dual combination resulted in a ≥3-log reduction, demonstrating synergistic bactericidal activity (**Fig. S11**).

Given this synergy in a wild-type (WT) background, we hypothesised it might extend to bedaquiline-resistant strains. The most common clinical mechanism of bedaquiline resistance involves disruption of *Rv0678*, leading to upregulation of the MmpS5-MmpL5 efflux pump. To model this, we generated a bedaquiline-resistant strain (BDQR^Rv0678^) with complete disruption of Rv0678 and a concomitant increase in MmpS5-MmpL5 expression (**Fig. S12**). Kill kinetics were assessed using JNJ-6887 and bedaquiline, alone and in combination, in both WT and BDQR^Rv0678^ backgrounds. In the WT strain, monotherapy with BDQ (5x MIC_90_) and combination both reduced CFU by ∼2-log after 21 days compared with the starting inoculum (**Fig. 3a**; **Fig. S13**). In this background, bedaquiline appeared to drive most of the killing. In contrast, bedaquiline monotherapy had no effect in the BDQR^Rv0678^ background, consistent with the resistance phenotype, while JNJ-6887 retained equivalent activity in both strains, indicating that the resistance mechanism did not affect MenG inhibition. Remarkably, combination treatment in the resistant background resulted in a ≥3-log reduction in CFU counts after 21 days, indicating that MenG inhibition not only resensitised the strain to bedaquiline but also retained its synergistic capacity (**Fig. 3a**; **Fig. S13**). Equivalent results were also obtained with JNJ-1866 in the BDQR^R0678^ background, and JNJ-6887 produced the same effect in an alternative *Rv0678*-mediated resistance strain carrying a distinct resistance-conferring mutation (**Fig. S14-15**). In contrast, no resensitisation was observed with linezolid under the same experimental conditions (**Fig. S13**).

**Fig. 3.**
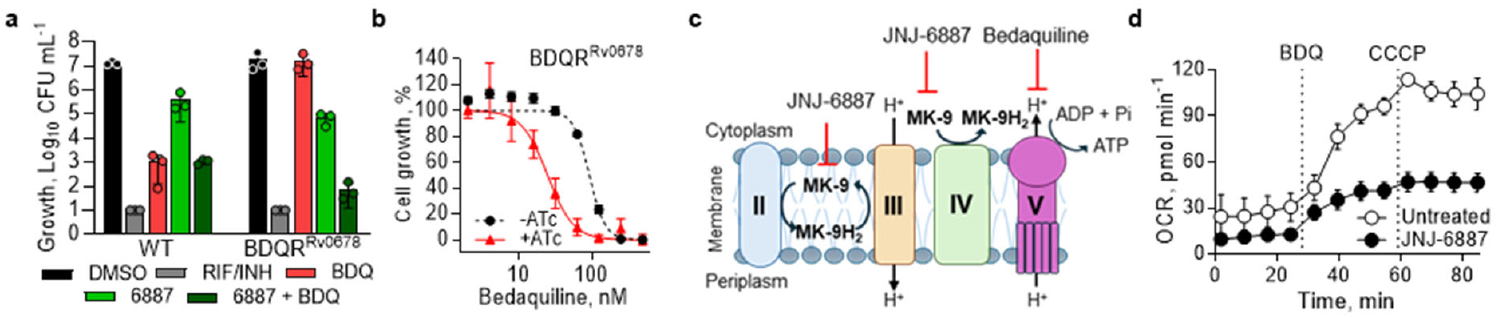
Resensitisation of bedaquiline resistance via MenG inhibition. **a**, Day-21 from a time-kill kinetics experiment of 0.5 µM BDQ (5x MIC_90_ activity in the WT background) and 0.2 µM JNJ-6887 (4x MIC_90_), alone and in combination, in WT and BDQR^Rv0678^ backgrounds. n = 3 technical replicates. Representative of two independent experiments shown. **b**, Dose-response curves of bedaquiline with the CRISPRi-mediated *menG* “low” knockdown strain (- ATc; MIC_50_ = 89.7 nM) compared with induction (+ATc; MIC_50_ = 23.6 nM). n = 3 technical replicates. Representative of three independent experiments shown. **c**, Brief outline of the ETC in *M. tuberculosis*. Complex II: succinate dehydrogenase; Complex III: Cytochrome *bc*_1_; Complex IV; Cytochrome *bd* oxidase; Complex V: ATP synthase. Expanded model of the ETC in **Fig. S17. d**, Oxygen consumption rate (OCR) assay in samples pre-treated with 660 nM JNJ-6887 for 3 days followed by the addition of 32 µM bedaquiline then 4 µM CCCP (carbonyl cyanide m-chlorophenyl hydrazone). n = 4 biological replicates. For panels **b-c, d**, data shown are mean ± s.d.

To confirm that resensitisation was not due to an off-target effect of the compounds, we used genetic knockdown of *menG* as an orthogonal approach. A CRISPRi-mediated *menG* “low” transcriptional knockdown, selected to avoid lethality during the assay, led to a 4-fold increase in bedaquiline susceptibility in the BDQR^Rv0678^ background, consistent with the effect observed using chemical inhibition (**Fig. 3b; Fig. S10**). Finally, to rule out the activation of known bedaquiline-resistance mechanisms, transcript profiling of the BDQR^Rv0678^ strain treated with JNJ-6887 revealed no differential expression of genes associated with resistance (**Fig. S16**). This supports the conclusion that MenG inhibition is specifically responsible for restoring bedaquiline activity in resistant strains.

MK-9 shuttles electrons along the ETC; therefore, MenG inhibition is expected to disrupt electron flow and reduce proton motive force (PMF), the electrochemical proton gradient that fuels ATP production (**Fig. 3c**; **Fig. S17**). Indeed, previous studies has demonstrated that MenG inhibition reduces oxygen consumption and impairs ATP synthesis (*38*). Efflux systems of the RND family, including MmpS5-MmpL5, are thought to be PMF-dependent (*40*). We therefore hypothesised that reduced PMF would impair efflux pump activity and, together with ETC disruption, account for synergy with ATP synthase inhibitors (complex V; bedaquiline) and the observed resensitisation phenotype. Although oxygen consumption rate (OCR) does not directly measure PMF, it does reflect changes in electron transport that influence proton pumping. Following 3-day exposure to 10x MIC_50_ JNJ-6887, basal OCR was reduced by ∼50% relative to untreated control (**Fig. 3d**). Addition of bedaquiline, previously shown to increase respiration rate, and CCCP, a classic uncoupler, highlighted the impairment of electron transfer through respiratory complexes caused by MenG inhibition. Short-term exposure to JNJ-6887 did not measurably alter membrane potential (**Fig. S18**), used here as a proxy for PMF-dependent efflux, suggesting that resensitisation is not explained by acute depolarisation. This is consistent with the delayed onset of activity observed with MenG inhibition (**Fig. 2e**). As a complementary approach, we tested whether combining bedaquiline with JNJ-6887 could restore the activity of JNJ-6887 in a MenG-resistant strain (**Fig. S19**). This combination produced a 5-log improvement in activity relative to either monotherapy, demonstrating that reciprocal resensitisation occurs for this drug combination. Finally, to investigate whether alternative respiratory pathways compensate for MenG inhibition, we assessed the activity of JNJ-6887 in a cytochrome *bd* (complex IV) knockout (*bd*KO) strain. Loss of cytochrome *bd* did not enhance sensitivity to JNJ-6887, suggesting that MenG inhibition is not compensated by this respiratory branch (**Fig. S20**). Taken together, these results demonstrate that MenG inhibitors resensitise *M. tuberculosis* BDQR^Rv0678^ to bedaquiline through combined disruption of bioenergetic pathways and impairment of the PMF.

### Enhancement of other TB therapies and in vivo bedaquiline resensitisation

We next assessed whether MenG inhibition could enhance the activity of other TB therapies and overcome resistance beyond bedaquiline. Macozinone and pretomanid both inhibit the formation of decaprenylphosphoryl-D-arabinose (DPA), an essential precursor for the mycobacterial cell wall, by disrupting DprE1 and DprE2, respectively (*41, 42*). DprE1 functions as a menaquinone-dependent dehydrogenase that facilitates FAD reoxidation (*43*), leading us to hypothesise that MenG inhibition would enhance the activity of both compounds (**Fig. 4a**). Indeed, time-kill kinetics showed that MenG inhibition enhanced the bactericidal activity of both drugs (**Fig. 4b**; **Fig. S21**). Despite its distinct mode of action, macozinone shares the same efflux-mediated resistance mechanism as bedaquiline (*14*); JNJ-6887 fully restored macozinone susceptibility in the BDQR^Rv0678^ strain.

**Fig. 4.**
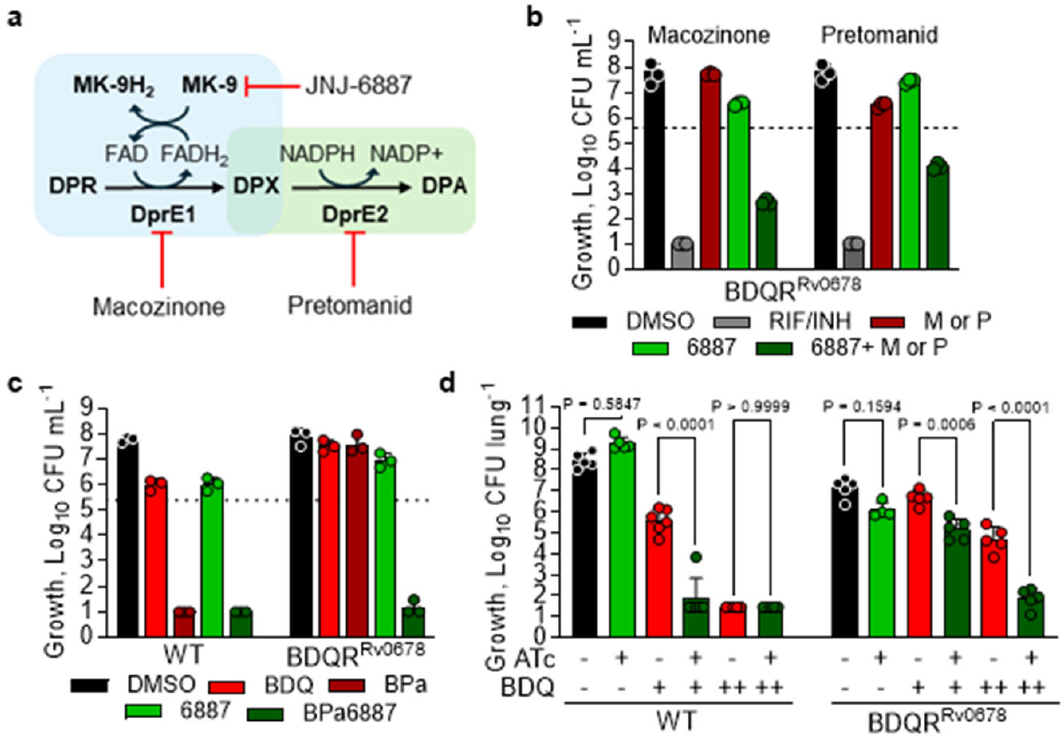
MenG inhibition enhances key TB drugs and restores resistance *in vivo*. **a**, The decaprenylphosphoryl-D-arabinose (DPA) pathway and its interactions with macozinone, pretomanid and JNJ-6887. Expanded explanation in **Fig. S21**. Decaprenylphosphoryl-D-ribose (DPR); decaprenylphosphoryl-2-keto-erythro-pentofuranose, DPX; flavin adenine dinucleotide, FAD; nicotinamide adenine dinucleotide phosphate, NADPH; menaquinol-9, MK-9H_2_. **b**, Day 21 from time-kill kinetics assays of 1.5 nM macozinone (M) or 500 nM pretomanid (P) with 0.2 µM JNJ-6887 in BDQR^Rv0678^. The dotted line indicates the starting inoculum. n = 3 technical replicates. Representative of two independent experiments shown. Full dataset shown in **Fig. S21. c**, Day 21 from a time-kill kinetics assay of combination treatment. Bedaquiline (0.5 µM; B), pretomanid (7 µM; Pa) and JNJ-6887 (0.2 µM; 6887). The dotted line indicates the starting inoculum. n = 3 technical replicates. Representative of two independent experiments shown. Full dataset shown in **Fig. S23. d**, *In vivo* efficacy of MenG “low” CRISPRi in WT (left) and BDQR^Rv0678^ (right) backgrounds, after 28-day treatment with induction with doxycycline included in the chow, with 6.25 mg kg^-1^ (+) or 25 mg kg^-1^ (++) bedaquiline (BDQ; once daily oral administration, 5/7 days). n = 5 mice. ATc/doxycycline positive conditions were pre-induced for 7 days prior to infection. Significance was calculated with one-way analysis of variance (ANOVA) with Šídák’s multiple comparisons. Full time-course is shown in **Fig. S24**. For panels **b-d**, data shown are mean ± s.d.

Given that JNJ-6887 enhanced the activity of bedaquiline and pretomanid, both backbone drugs for DR-TB treatment, we next investigated whether JNJ-6887 could restore bedaquiline activity in the BDQR^Rv0678^ background using human exposure-equivalent concentrations of bedaquiline and pretomanid. In contrast to WT, bedaquiline-pretomanid (BPa) was inactive in the BDQR^Rv0678^ strain, whereas the addition of only 4x MIC_90_ JNJ-6887, which had no activity as a monotherapy, fully restored BPa activity (**Fig. 4c**; **Fig. S22**). These findings support the inclusion of a MenG-targeting inhibitor in future DR-TB combination therapies without the need to determine efflux-based resistance, aligning with drug-susceptibility testing-independent treatment strategies envisioned by the PAN-TB framework (*44*).

To assess whether the resensitisation phenotype extended to more physiologically relevant conditions, we used a 3-day *ex vivo* THP-1 macrophage infection model with the BDQR^Rv0678^ strain, which similarly demonstrated robust resensitisation to bedaquiline in the presence of JNJ-6887 (**Fig. S23**). Encouraged by these findings, we next investigated whether this effect translated *in vivo*. Mice were infected with either WT or BDQR^Rv0678^ strains for 7 days and treated with a subcutaneous long-acting injectable (LAI) formulation of JNJ-1866, a strategy previously shown to enhance *in vivo* compound exposure levels (*45*) (**Fig. S2**; **Table S6**). In a WT background, 1,250 mg kg^-1^ JNJ-1866 LAI clearly enhanced bedaquiline activity; however, in the BDQR^Rv0678^ background, combination treatment showed only a non-significant trend towards resensitisation (**Fig. S3, 24**). As an alternative approach, we used the *menG* “low” knockdown strains in mice (**Fig. S3**). In these models, we confirmed that bedaquiline had reduced activity in the BDQR^Rv0678^ background compared with WT, while *menG* knockdown alone had minimal impact in either strain. Combining *menG* knockdown with 6.25 mg kg^-1^ bedaquiline in the WT background enhanced activity of the combination, matching the synergy observed *in vitro*. Importantly, combination with 25 mg kg^-1^ bedaquiline in the BDQR^Rv0678^ background resulted in a 3-log reduction in bacterial load (**Fig. 4d**; **Fig. S24**), restoring levels comparable to WT activity. These findings provide clear *in vivo* proof-of-concept that MenG inhibition can resensitise resistant strains to bedaquiline. Finally, to complement this work, we used *Mycobacterium marinum*, one of the closest genetic relatives of *M. tuberculosis* and a pathogen that enables *in vivo* efficacy testing in its natural host, the zebrafish. Treatment with JNJ-1866 demonstrated clear *in vivo* proof-of-concept after 2 days. We subsequently generated an efflux-mediated (*MMAR-1007*; *Rv0678*-equivalent) bedaquiline-resistant *M. marinum* strain and confirmed that it conferred *in vivo-*relevant resistance to bedaquiline. Importantly, inhibition of MenG resensitised this strain to bedaquiline, mirroring the phenotype observed in *M. tuberculosis* (**Fig. S25**).

### Pathway inhibition restores bedaquiline susceptibility

The shikimate biosynthetic pathway provides the chorismate precursor essential for the MK-9 biosynthesis, linking aromatic amino acid biosynthesis to respiratory energy generation (**Fig. 5a**). To determine whether resensitisation of bedaquiline activity was a general consequence of menaquinone biosynthesis disruption, we performed matched time-kill kinetics using CRISPRi-mediated “low” knockdown of *menG, menE, aroG* and *aroK*, targeting multiple steps across the menaquinone and upstream shikimate biosynthetic pathways. Experiments were carried out in both WT and BDQR^Rv0678^ backgrounds (**Fig. 5b-e**; **Fig. S10, 26**). Monotherapy with 0.5 µM bedaquiline had no impact on CFU in the BDQR^Rv0678^ background. Similarly, individual knockdown of each gene produced only a modest reduction in CFU, consistent with a “low” knockdown phenotype. However, combining bedaquiline exposure with “low” knockdown led to a marked and reproducible reduction in CFU (**Fig. 5b-e**), confirming a resensitisation phenotype comparable to that observed with JNJ-6887 and suggesting that disruption of any gene in the two pathways can resensitise bedaquiline resistance.

**Fig. 5.**
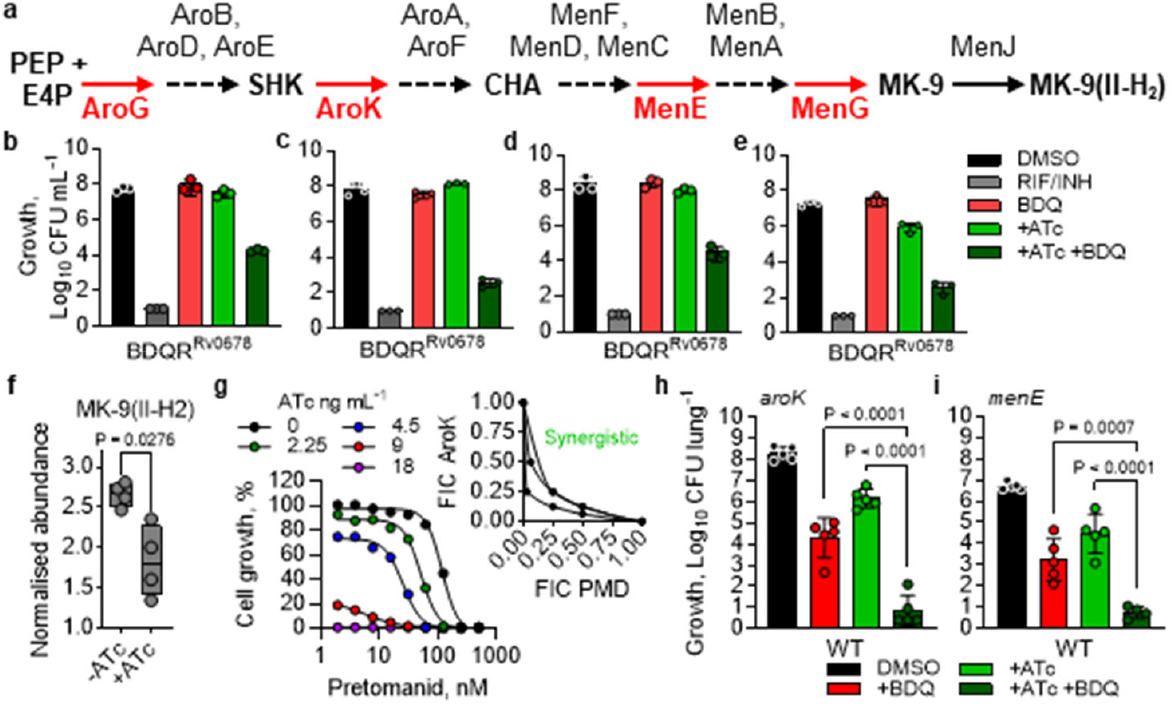
Drug resistance rescued by entire menaquinone biosynthesis pathway. **a**, Schematic of the shikimate and menaquinone biosynthetic pathways in *M. tuberculosis*. **b-e**, Day 14 CFU counts from time-kill assays using CRISPRi-mediated “low” knockdown of (**b**) *aroG*, (**c**) *aroK*, (**d**) *menE* or (**e**) *menG* with 0.5 µM BDQ (10× MIC_90_) in BDQR^Rv0678^ background. Strains were pre-induced with 100 ng mL^-1^ ATc for 5 days (*aroG* and *aroK*) or 7 days (*menE* and *menG*). DMSO (black); rifampicin (RIF; 14.58 µM) and isoniazid (INH; 5.8 µM) as positive kill control (dark grey). Representative of two independent experiments with three biological replicates. Time-kill kinetics data are shown in **Fig. S26. f**, LC-MS analysis showing MK-9(II-H_2_) reduction in the *aroK* “high” CRISPRi strain after induction with ATc for 5 days. n = 4 biological replicates. Representative of two independent experiments. Significance was calculated with two-sided (Bonferroni–Dunn) Student’s t-test with Welch correction. **g**, Dose-response curves of pretomanid following *aroK* “high” transcript reduction in an ATc-dependent manner, showing 13-fold increase in inhibitor potency after 14 days. Representative of three independent experiments. Inset: isobologram using fractional inhibitory concentrations (FIC) of pretomanid (PMD) and aroK induction (ATc concentration). n = 3 independent experiments. **h-i**, *In vivo* efficacy of CRISPRi-mediated aroK “high” (**h**) and *menE* “low” (**i**) knockdown in a WT background, induced with doxycycline in chow, and in combination with 6.25 mg kg^-1^ bedaquiline (BDQ; once daily oral administration, 5/7 days). ATc/doxycycline positive *menE* conditions were pre-induced for 7 days prior to infection. n = 5 mice. Significance was calculated with one-way analysis of variance (ANOVA) with Šídák’s multiple comparisons. Full time-course is shown in **Fig. S28**. For panels **b-f, h-i**, the data shown are mean ± s.d.

In the absence of validated inhibitors, CRISPRi-mediated “high” knockdowns of *aroG* and *aroK* were generated to facilitate combination studies and provide further genetic validation that these enzymes represent viable drug targets (**Fig. S10**). Silencing of *aroG* and *aroK* produced a robust bactericidal effect, reaching the limit of detection after 14 days, in agreement with published vulnerability scores for these enzymes (*26*) (**Fig. S27**). LC-MS analysis of the *aroK* “high” knockdown strain revealed increased shikimate, the substrate of AroK, and reduced MK-9(II-H_2_) levels, consistent with its role upstream of the menaquinone biosynthesis pathway and supporting the hypothesis that the resensitisation phenotype is specifically due to inhibition of menaquinone biosynthesis (**Fig. 5f; Fig. S27**). We next used our *aroK* “high” knockdown strain to investigate potential enhancement of pretomanid activity (**Fig. 5g**). This resulted in a robust, ATc concentration-dependent 13-fold increase in susceptibility to pretomanid; conversion of these data into an isobologram demonstrated that pretomanid and ATc-dependent *aroK* knockdown act synergistically (**Fig. 5g**). Finally, in an acute mouse model, combining *aroG* “high”, *aroK* “high” or *menE* “low” knockdown with 6.25 mg kg^-1^ bedaquiline in the WT background significantly enhanced bedaquiline activity compared with either monotherapy or knockdown alone after 21 days (**Fig. 5h-i; Fig. S28**), replicating the *in vivo* resensitisation phenotype seen with *menG* inhibition. These findings indicate that disruption of any component of menaquinone biosynthesis is sufficient to both enhance the activity of other compounds and overcome bedaquiline resistance, highlighting enzymes from both pathways as potential “resistance-breaking” drug targets that should be prioritised for future screening efforts.

## Discussion

The emergence of bedaquiline resistance represents a growing threat to DR-TB treatment. Current global prevalence of bedaquiline resistance remains uncertain due to the absence of robust prospective surveillance data, but it appears to be increasing in some regions, with estimates ranging from 0.2% in China in 2022 (*46*) to 14% in Mozambique in 2021 (*7*). The MmpS5-MmpL5 efflux pump-mediated resistance mechanism also confers cross-resistance to clofazimine, benzothiazinones (including macozinone), Q203 and next-generation ATP synthase inhibitors (TBAJ-587 and TBAJ-876) (*14*). Clinically, baseline *Rv0678* mutations in bedaquiline-naïve patients enrolled in early registrational trials did not affect initial culture conversion (*12, 47, 48*). However, lower conversion rates were observed among patients who acquired *Rv0678* mutations compared with those who did not (*12*). Patients with bedaquiline-resistant TB also have poorer outcomes overall, with only 57% achieving treatment success compared to 72% among those with susceptible infections (*49*). The risk of poor outcomes is further increased when *Rv0678* variants occur in patients previously exposed to bedaquiline or clofazimine. This risk is highest in pre–extensively drug-resistant TB (pre-XDR-TB) and extensively drug-resistant TB (XDR-TB) populations, particularly in the presence of fluoroquinolone resistance and when treatment regimens lack additional active drugs (*50, 51*). Currently, alternative regimens used when bedaquiline cannot be given are longer and not standardised, supported by weaker evidence of efficacy and associated with greater toxicity (*52*). As bedaquiline remains the cornerstone of DR-TB treatment, strategies that both limit the spread of resistance and overcome established resistance are urgently required.

Combination therapy remains fundamental to TB control. Drugs with distinct modes of action can interact synergistically or antagonistically, producing effects greater or less than expected from the addition of monotherapies. Synergistic combinations could be particularly valuable clinically, as they may accelerate bacterial clearance, shorten treatment duration, reduce the risk of resistance and lessen adverse effects. Here, we show that MenG inhibition enhances the activity of bedaquiline, pretomanid and macozinone. This is consistent with earlier reports (*38*), demonstrating that MenG inhibition also synergises with the first-line drugs rifampicin and isoniazid. Previous work has likewise shown that MenA inhibition enhances the activity of clofazimine (*39*).

Crucially, the synergy observed with MenG inhibition resensitised strains with clinically relevant efflux-mediated bedaquiline resistance. Previous work with non-clinical compounds has shown that combinations containing chlorpromazine, an ETC inhibitor, can restore the efficacy of spectinomycin, a protein synthesis inhibitor, against *M. tuberculosis* strains carrying spectinomycin-resistance mutations, highlighting the potential of potentiating combinations to bypass resistance (*53*). Furthermore, although efflux pump inhibitors such as verapamil can also restore bedaquiline activity *in vitro* (**Fig. S29**), they lack bactericidal activity as monotherapies and do not restore bedaquiline efficacy *in vivo* (*12, 54*). Inhibiting menaquinone biosynthesis therefore provides a more effective strategy to reverse resistance.

Ongoing work exploring non-bactericidal “booster” molecules that enhance ethionamide activity further highlights the potential of agents that strengthen existing regimens (*20, 21*). Importantly, MenG inhibition is intrinsically bactericidal, offering a key advantage over these booster compounds. As a result, menaquinone biosynthesis inhibitors could contribute directly to bacterial clearance while simultaneously enhancing the activity of partner drugs. This dual action raises the possibility of meaningful treatment shortening. Our current compound series lacks high-level efficacy in a mouse model of infection; however, this broader principle is supported by work on cytochrome *bc*_*1*_ inhibitors, which exhibit limited or static bactericidal activity but show strong combinatorial effects with multiple drug classes (*55*).

The representative molecule from the current series, JNJ-1866, has served as a valuable tool compound for characterising the mode of action and assessing the relevance of MenG. However, further optimisation is required to improve drug-like properties and achieve a druggable candidate. Addressing its metabolic instability will be essential for defining the PK/PD relationship in animal models and for improving the translatability of these efficacy models to human settings.

Both the menaquinone and shikimate biosynthetic pathways are essential in mycobacteria (*26, 56*) but absent in humans, making them attractive therapeutic targets. *M. tuberculosis* relies on MK-9 as the principal electron carrier linking primary dehydrogenases to terminal oxidoreductases in the ETC. We demonstrated that MenG inhibition depletes menaquinone levels, disrupts electron transport and, when combined with drugs exported via the MmpL-MmpS5 efflux system, restores susceptibility to near WT levels. As both ATP synthase and menaquinone-dependent respiration are essential, dual inhibition may also enhance killing of persistent subpopulations, with potential to shorten treatment duration and reduce relapse risk.

Given the rising burden of DR-TB, prolonged treatment duration and increasing resistance rates, new therapeutic strategies are urgently needed. Our study provides clear *in vivo* proof-of-principle that inhibiting menaquinone biosynthesis can restore the efficacy of bedaquiline against efflux-mediated resistance. Since depletion of MK-9 appears to underlie this resensitisation, targeting any essential enzyme within the shikimate or menaquinone pathways, such as MenA, which has already been validated as druggable (*57*), may yield similar outcomes. Together, these findings confirm these pathways as critical targets in *M. tuberculosis* and, for the first time, demonstrate that their inhibition can reverse efflux-mediated bedaquiline resistance, potentially extending the clinical lifespan of key TB drugs and informing the rational design of next-generation combination regimens aimed at shortening therapy and preventing the spread of resistance.

## Disclaimer

This article is a preprint reporting new research and has not been peer-reviewed; it should not be used to guide clinical practice or be reported in the press as conclusive.

## Acknowledgements

The authors would like to thank Peggy Janssens and Peggy Geluykens for technical support; Dr Lieve Lammens for scientific toxicology support; Dr Ingrid Eshun-Wilsonova for critical scientific review; Anne T. Henze for administrative support at Johnson & Johnson. Tesnim Alattar, Dr Suzanne Harris and Dr Shahida Rafique for technical assistance at the London School of Hygiene & Tropical Medicine. James Stratta for LC support at Waters. The authors would like to thank the TB Drug Accelerator (TBDA) MenG Working Group for their guidance and support during the progression of this programme. We thank GSK for providing access to compounds from their published TB-active set. The *M. tuberculosis* mc^2^ 6230 strain was kindly provided by Dr Bill Jacob.

## Author contributions

CA-P, DAL and RJW managed and coordinated the project. CMMV, JEGP, KS, PJ and AAT coordinated and guided the experimental medicinal chemistry or compound scale-up. JW, ED, VvH, JH, SD and MG planned, coordinated and performed the work related to the time kill kinetics with compounds. JD profiled the compounds in clinical isolates. BT and AJCS planned and performed the oxygen consumption experiment. VP and DC planned and performed the MenG enzymatic assay. JS, AL and VvH generated and profiled the MenG resistant strains. VM generated and profiled the bedaquiline resistant strain. AV performed the computational modelling and docking. WHP, ED, BT, AJCS and GL-M performed and analysed the metabolomics experiments. AW, ME and BS planned, coordinated and performed the DMPK and toxicology analysis. MG, NC and ND planned, performed and analysed the single-cell imaging data and membrane potential measurements. TGC provided access to the clinical isolates. HP, JD, RF and SM planned and performed experiments with *M. marinum* and zebrafish. JD planned and performed THP-1 macrophage experiments. RD-A, HF, ASP, MVL and MLH provided intellectual guidance. JD, ED, SJW and CD planned and performed the work related to the CRISPRi strains. CA-P, SD, MG, MP, VvH, JH and JW planned, coordinated and performed all of the *in vivo* studies. MG, SD, VvH and JH performed RNA extractions from *in vivo* samples. AK and RJW provided overall supervision of the study. RJW wrote the manuscript and coordinated its assembly, with contributions from CA-P, DAL and AK and comments from all other authors. All authors reviewed the manuscript for intellectual content and gave final approval to submit for publication.

## Data, code, and materials availability

All data generated or analysed during this study are included in this published article (and its supplementary information files).

## References and notes

1 World Health Organisation Global Tuberculosis Programme. Global tuberculosis report 2025, https://www.who.int/publications/i/item/9789240116924 (2025).

2 Andries, K. et al. Acquired Resistance of Mycobacterium tuberculosis to Bedaquiline. PLOS ONE 9, e102135, doi:10.1371/journal.pone.0102135 (2014).

3 Waller, N. J. E., Cheung, C.-Y., Cook, G. M. & McNeil, M. B. The evolution of antibiotic resistance is associated with collateral drug phenotypes in Mycobacterium tuberculosis. Nat Commun 14, 1517, doi:10.1038/s41467-023-37184-7 (2023).

4 Andries, K. et al. A diarylquinoline drug active on the ATP synthase of Mycobacterium tuberculosis. Science 307, 223–227, doi:10.1126/science.1106753 (2005).

5 Conradie, F. et al. Bedaquiline-Pretomanid-Linezolid Regimens for Drug-Resistant Tuberculosis. N Engl J Med 387, 810–823, doi:10.1056/NEJMoa2119430 (2022).

6 Conradie, F. et al. Treatment of Highly Drug-Resistant Pulmonary Tuberculosis. N Engl J Med 382, 893–902, doi:10.1056/NEJMoa1901814 (2020).

7 Rukasha, I., Kaapu, K. G., Fortune, S. M. & Lekalakala-Mokaba, M. R. Treatment Outcomes for Drug-Resistant Tuberculosis Patients on Bedaquiline-Based Regimens in a Mostly Rural South Africa. Infect Drug Resist 18, 1819–1829, doi:10.2147/idr.S502302 (2025).

8 Omar, S. V., Ismail, F., Ndjeka, N., Kaniga, K. & Ismail, N. A. Bedaquiline-Resistant Tuberculosis Associated with Rv0678 Mutations. N Engl J Med 386, 93–94, doi:10.1056/NEJMc2103049 (2022).

9 Barilar, I. et al. Emergence of bedaquiline-resistant tuberculosis and of multidrug-resistant and extensively drug-resistant Mycobacterium tuberculosis strains with rpoB Ile491Phe mutation not detected by Xpert MTB/RIF in Mozambique: a retrospective observational study. The Lancet Infectious Diseases 24, 297–307, doi:10.1016/S1473-3099(23)00498-X (2024).

10 Koul, A. et al. Diarylquinolines target subunit c of mycobacterial ATP synthase. Nat Chem Biol 3, 323–324, doi:10.1038/nchembio884 (2007).

11 Mahajan, R. Bedaquiline: First FDA-approved tuberculosis drug in 40 years. Int J Appl Basic Med Res 3, 1–2, doi:10.4103/2229-516x.112228 (2013).

12 Matteelli, A., Carvalho, A. C., Dooley, K. E. & Kritski, A. TMC207: the first compound of a new class of potent anti-tuberculosis drugs. Future Microbiol. 5, 849–858, doi:10.2217/fmb.10.50 (2010).

13 Kaniga, K., Lounis, N., Zhuo, S., Bakare, N. & Andries, K. Impact of Rv0678 mutations on patients with drug-resistant TB treated with bedaquiline. Int J Tuberc Lung Dis 26, 571–573, doi:10.5588/ijtld.21.0670 (2022).

14 Liu, Y. et al. Reduced Susceptibility of Mycobacterium tuberculosis to Bedaquiline During Antituberculosis Treatment and Its Correlation With Clinical Outcomes in China. Clinical Infectious Diseases 73, e3391–e3397, doi:10.1093/cid/ciaa1002 (2020).

15 Nimmo, C. et al. Detection of a historic reservoir of bedaquiline/clofazimine resistance-associated variants in Mycobacterium tuberculosis. Genome Med 16, 34, doi:10.1186/s13073-024-01289-5 (2024).

16 Cook, G. M. et al. Oxidative Phosphorylation as a Target Space for Tuberculosis: Success, Caution, and Future Directions. Microbiol Spectr 5, doi:10.1128/microbiolspec.TBTB2-0014-2016 (2017).

17 Gengenbacher, M., Rao, S. P. S., Pethe, K. & Dick, T. Nutrient-starved, non-replicating Mycobacterium tuberculosis requires respiration, ATP synthase and isocitrate lyase for maintenance of ATP homeostasis and viability. Microbiology (Reading) 156, 81–87, doi:10.1099/mic.0.033084-0 (2010).

18 Koul, A. et al. Diarylquinolines Are Bactericidal for Dormant Mycobacteria as a Result of Disturbed ATP Homeostasis*. J. Biol. Chem. 283, 25273–25280, doi: 10.1074/jbc.M803899200 (2008).

19 Nunes, J. E. S. et al. Mycobacterium tuberculosis Shikimate Pathway Enzymes as Targets for the Rational Design of Anti-Tuberculosis Drugs. Molecules 25, 1259 (2020).

20 Flipo, M. et al. The small-molecule SMARt751 reverses Mycobacterium tuberculosis resistance to ethionamide in acute and chronic mouse models of tuberculosis. Sci Transl Med 14, eaaz6280, doi:10.1126/scitranslmed.aaz6280 (2022).

21 Gries, R. et al. Discovery of dual-active ethionamide boosters inhibiting the Mycobacterium tuberculosis ESX-1 secretion system. Cell Chemical Biology 31, 699-711.e696, doi:10.1016/j.chembiol.2023.12.007 (2024).

22 Ballell, L. et al. Fueling open-source drug discovery: 177 small-molecule leads against tuberculosis. ChemMedChem 8, 313–321, doi:10.1002/cmdc.201200428 (2013).

23 Wakamoto, Y. et al. Dynamic Persistence of Antibiotic-Stressed Mycobacteria. Science 339, 91–95, doi:10.1126/science.1229858 (2013).

24 Dhar, N. et al. Rapid cytolysis of Mycobacterium tuberculosis by faropenem, an orally bioavailable β-lactam antibiotic. Antimicrob Agents Chemother 59, 1308–1319, doi:10.1128/aac.03461-14 (2015).

25 Crampin, A. C., Glynn, J. R. & Fine, P. E. What has Karonga taught us? Tuberculosis studied over three decades. Int J Tuberc Lung Dis 13, 153–164 (2009).

26 Bosch, B. et al. Genome-wide gene expression tuning reveals diverse vulnerabilities of M. tuberculosis. Cell 184, 4579-4592.e4524, doi:10.1016/j.cell.2021.06.033 (2021).

27 DeJesus, M. A. et al. Comprehensive Essentiality Analysis of the Mycobacterium tuberculosis Genome via Saturating Transposon Mutagenesis. mBio 8, 10.1128/mbio.02133-02116, doi:10.1128/mbio.02133-16 (2017).

28 Dhiman, R. K. et al. Menaquinone synthesis is critical for maintaining mycobacterial viability during exponential growth and recovery from non-replicating persistence. Mol Microbiol 72, 85–97, doi:10.1111/j.1365-2958.2009.06625.x (2009).

29 Phelan, J. et al. An open-access dashboard to interrogate the genetic diversity of Mycobacterium tuberculosis clinical isolates. Scientific Reports 14, 24792, doi:10.1038/s41598-024-75818-y (2024).

30 Dai, Y. N. et al. Crystal structures and catalytic mechanism of the C-methyltransferase Coq5 provide insights into a key step of the yeast coenzyme Q synthesis pathway. Acta Crystallogr D Biol Crystallogr 70, 2085–2092, doi:10.1107/s1399004714011559 (2014).

31 Jumper, J. et al. Highly accurate protein structure prediction with AlphaFold. Nature 596, 583–589, doi:10.1038/s41586-021-03819-2 (2021).

32 Varadi, M. et al. AlphaFold Protein Structure Database: massively expanding the structural coverage of protein-sequence space with high-accuracy models. Nucleic Acids Research 50, D439–D444, doi:10.1093/nar/gkab1061 (2021).

33 Das, K., Lewi, P. J., Hughes, S. H. & Arnold, E. Crystallography and the design of anti-AIDS drugs: conformational flexibility and positional adaptability are important in the design of non-nucleoside HIV-1 reverse transcriptase inhibitors. Prog Biophys Mol Biol 88, 209–231, doi:10.1016/j.pbiomolbio.2004.07.001 (2005).

34 Upadhyay, A. et al. Mycobacterial MenJ: An Oxidoreductase Involved in Menaquinone Biosynthesis. ACS Chemical Biology 13, 2498–2507, doi:10.1021/acschembio.8b00402 (2018).

35 Dunovska, K., Klapkova, E., Sopko, B., Cepova, J. & Prusa, R. LC-MS/MS quantitative analysis of phylloquinone, menaquinone-4 and menaquinone-7 in the human serum of a healthy population. PeerJ 7, e7695, doi:10.7717/peerj.7695 (2019).

36 Pujari, V., Rozman, K., Dhiman, R. K., Aldrich, C. C. & Crick, D. C. Mycobacterial MenG: Partial Purification, Characterization, and Inhibition. ACS Infectious Diseases 8, 2430–2440, doi:10.1021/acsinfecdis.2c00190 (2022).

37 McNeil, M. B., Keighley, L. M., Cook, J. R., Cheung, C. Y. & Cook, G. M. CRISPR interference identifies vulnerable cellular pathways with bactericidal phenotypes in Mycobacterium tuberculosis. Mol Microbiol 116, 1033–1043, doi:10.1111/mmi.14790 (2021).

38 Sukheja, P. et al. A Novel Small-Molecule Inhibitor of the Mycobacterium tuberculosis Demethylmenaquinone Methyltransferase MenG Is Bactericidal to Both Growing and Nutritionally Deprived Persister Cells. mBio 8, 10.1128/mbio.02022-02016, doi:10.1128/mbio.02022-16 (2017).

39 Berube, B. J. et al. Novel MenA Inhibitors Are Bactericidal against Mycobacterium tuberculosis and Synergize with Electron Transport Chain Inhibitors. Antimicrob Agents Chemother 63, doi:10.1128/aac.02661-18 (2019).

40 Chim, N. et al. The Structure and Interactions of Periplasmic Domains of Crucial MmpL Membrane Proteins from Mycobacterium tuberculosis. Chemistry & Biology 22, 1098–1107, doi:10.1016/j.chembiol.2015.07.013 (2015).

41 Makarov, V. et al. Benzothiazinones Kill Mycobacterium tuberculosis by Blocking Arabinan Synthesis. Science 324, 801–804, doi:10.1126/science.1171583 (2009).

42 Abrahams, K. A. et al. DprE2 is a molecular target of the anti-tubercular nitroimidazole compounds pretomanid and delamanid. Nature Communications 14, 3828, doi:10.1038/s41467-023-39300-z (2023).

43 Neres, J. et al. Structural basis for benzothiazinone-mediated killing of Mycobacterium tuberculosis. Sci Transl Med 4, 150ra121, doi:10.1126/scitranslmed.3004395 (2012).

44 Pan-TB. Our Approach, https://www.pan-tb.org/what-is-pan-tb/our-approach/ (2025).

45 Lamprecht, D. et al. Targeting de novo purine biosynthesis for tuberculosis treatment. Nature 644 (8075), 214–220, doi:10.1038/s41586-025-09177-7 (2025).

46 Li, S. et al. The emerging threat of fluroquinolone-, bedaquiline-, and linezolid-resistant Mycobacterium tuberculosis in China: Observations on surveillance data. J Infect Public Health 17, 137–142, doi:10.1016/j.jiph.2023.11.018 (2024).

47 Diacon, A. H. et al. Multidrug-resistant tuberculosis and culture conversion with bedaquiline. N Engl J Med 371, 723–732, doi:10.1056/NEJMoa1313865 (2014).

48 Pym, A. S. et al. Bedaquiline in the treatment of multidrug- and extensively drug-resistant tuberculosis. Eur Respir J 47, 564–574, doi:10.1183/13993003.00724-2015 (2016).

49 Mdlenyani, L. et al. Treatment outcomes of bedaquiline-resistant tuberculosis: a retrospective and matched cohort study. The Lancet Infectious Diseases 25, 1149–1158, doi:10.1016/S1473-3099(25)00218-X (2025).

50 Ismail, N. A. et al. Assessment of epidemiological and genetic characteristics and clinical outcomes of resistance to bedaquiline in patients treated for rifampicin-resistant tuberculosis: a cross-sectional and longitudinal study. Lancet Infect Dis 22, 496–506, doi:10.1016/s1473-3099(21)00470-9 (2022).

51 U.S. Food and Drug Administration. Prescribing information for bedaquiline, https://www.accessdata.fda.gov/drugsatfda_docs/label/2012/204384s000lbl.pdf (2012).

52 World Health Organisation. Treatment of drug-resistant TB using longer regimens. https://tbksp.who.int/en/node/2978

53 Omollo, C. et al. Developing synergistic drug combinations to restore antibiotic sensitivity in drug-resistant Mycobacterium tuberculosis. Antimicrob Agents Chemother 65, doi:10.1128/aac.02554-20 (2023).

54 Gupta, S. et al. Efflux inhibition with verapamil potentiates bedaquiline in Mycobacterium tuberculosis. Antimicrob Agents Chemother 58, 574–576, doi:10.1128/aac.01462-13 (2014).

55 Aguilar-Pérez, C. et al. The role of cytochrome bc1 inhibitors in future tuberculosis treatment regimens. Nature Communications, 16, 9344, doi:10.1038/s41467-025-64427-6 (2025).

56 Puffal, J., Mayfield, J. A., Moody, D. B. & Morita, Y. S. Demethylmenaquinone Methyl Transferase Is a Membrane Domain-Associated Protein Essential for Menaquinone Homeostasis in Mycobacterium smegmatis. Frontiers in Microbiology 9, doi:10.3389/fmicb.2018.03145 (2018).

57 Debnath, J. et al. Discovery of selective menaquinone biosynthesis inhibitors against Mycobacterium tuberculosis. J Med Chem 55, 3739–3755, doi:10.1021/jm201608g (2012).

58 Cole, S. T. et al. Deciphering the biology of Mycobacterium tuberculosis from the complete genome sequence. Nature 393, 537–544, doi:10.1038/31159 (1998).

59 Manina, G. & Dhar, N. Single-Cell Analysis of Mycobacteria Using Microfluidics and Time-Lapse Microscopy. Methods Mol Biol 2314, 205–229, doi:10.1007/978-1-0716-1460-0_8 (2021).

60 Schindelin, J. et al. Fiji: an open-source platform for biological-image analysis. Nat Methods 9, 676–682, doi:10.1038/nmeth.2019 (2012).

61 Schmittgen, T. D. & Livak, K. J. Analyzing real-time PCR data by the comparative C(T) method. Nat Protoc 3, 1101–1108, doi:10.1038/nprot.2008.73 (2008).

62 Berg, K. et al. SAR study of piperidine derivatives as inhibitors of 1,4-dihydroxy-2-naphthoate isoprenyltransferase (MenA) from Mycobacterium tuberculosis. Eur J Med Chem 249, 115125, doi:10.1016/j.ejmech.2023.115125 (2023).

63 Lamprecht, D. A. et al. Turning the respiratory flexibility of Mycobacterium tuberculosis against itself. Nature Communications 7, 12393, doi:10.1038/ncomms12393 (2016).

64 Kingdon, A. D. H., Meosa-John, A.-R., Batt, S. M. & Besra, G. S. Vanoxerine kills mycobacteria through membrane depolarization and efflux inhibition. Frontiers in Microbiology Volume 14 -2023, doi:10.3389/fmicb.2023.1112491 (2023).

65 Matty, M. A., Oehlers, S. H. & Tobin, D. M. Live Imaging of Host-Pathogen Interactions in Zebrafish Larvae. Methods Mol Biol 1451, 207–223, doi:10.1007/978-1-4939-3771-4_14 (2016).

66 Takaki, K., Davis, J. M., Winglee, K. & Ramakrishnan, L. Evaluation of the pathogenesis and treatment of Mycobacterium marinum infection in zebrafish. Nature Protocols 8, 1114–1124, doi:10.1038/nprot.2013.068 (2013).

67 Lu, H. R. et al. Identifying Acute Cardiac Hazard in Early Drug Discovery Using a Calcium Transient High-Throughput Assay in Human-Induced Pluripotent Stem Cell-Derived Cardiomyocytes. Frontiers in Physiology 13, doi:10.3389/fphys.2022.838435 (2022).

68 Sharma, P. et al. Evolution of Small Molecule Inhibitors of Mycobacterium tuberculosis Menaquinone Biosynthesis. Journal of Medicinal Chemistry 68, 5774–5803, doi:10.1021/acs.jmedchem.4c03156 (2025).

69 Huitric, E. et al. Rates and mechanisms of resistance development in Mycobacterium tuberculosis to a novel diarylquinoline ATP synthase inhibitor. Antimicrob Agents Chemother 54, 1022–1028, doi:10.1128/aac.01611-09 (2010).

70 Yang, J. et al. Molecular characteristics and in vitro susceptibility to bedaquiline of Mycobacterium tuberculosis isolates circulating in Shaanxi, China. International Journal of Infectious Diseases 99, 163–170, doi:10.1016/j.ijid.2020.07.044 (2020).

