## Supplementary material for "Menaquinone depletion resensitises bedaquiline-resistant tuberculosis"

**The PDF file includes:**

Materials and Methods

Supplementary Text

Figs. S1 to S30

Tables S1 to S8

References mentioned in Supplementary

**Other Supplementary Materials for this manuscript include the following:**

Movies S1

Materials and Methods

Animal Ethics statement

All *in vivo* studies in mice were performed at Johnson & Johnson Innovative Medicines in Beerse within a certified BSL-3 facility, in accordance with European Directive 2010/63/EU and national regulations governing the use of animals for scientific purposes. All procedures were approved by the Johnson & Johnson Innovative Medicines Ethics Committee, which has been accredited by the Association for Assessment and Accreditation of Laboratory Animal Care International (AAALAC) since 2004 (unit number 001131; https://www.aaalac.org/). Zebrafish experiments were performed in accordance with the Animals (Scientific Procedures) Act 1986 and were approved by the UK Home Office (PPL PP5900632).

Bacterial strains and growth conditions

*Mycobacterium tuberculosis* H37Rv was cultured in Middlebrook 7H9 broth (BD) supplemented with 10% oleic acid-albumin-dextrose-catalase (OADC; BD), 0.2-0.5% glycerol and 0.05% Tween-80 (Sigma-Aldrich) at 37 °C, or on Middlebrook 7H10 solid agar (BD) supplemented with 10% OADC (Difco), 0.5% glycerol and, where required, 0.4% (w/v) activated charcoal. MenG inhibitor- and bedaquiline-resistant strains were maintained under the same conditions as WT. The *M. tuberculosis* H37Rv strain used in this study was kindly provided by Roland Brosch, Institut Pasteur, France and originates from a 1999 stock closely related to the strain sequenced in 1998 (*58*).

Clinical and drug-resistant isolates

The Karonga study and follow-up work were approved by the Health Sciences Research Committee in Malawi (#424) and by the London School of Hygiene & Tropical Medicine ethics committee (#5067). Consent was obtained for three sputum samples collected from each patient. Additional resistant isolates including MDR, pre-XDR and *in vitro* strains harbouring resistance-conferring mutations were purchased from the Belgian Coordinated Collections of Microorganisms at the Institute of Tropical Medicine, Antwerp, Belgium.

Determination of MIC_90_ and MBC_99_

MIC_90_ values were determined in Middlebrook 7H9 broth. *M. tuberculosis* cultures were adjusted to an inoculum of approximately 5 x 10^5^ CFU mL^-1^ and added to a drug dilution series prepared in 96-well or 384-well plates, with compounds diluted from 100x DMSO stocks. Plates were incubated for 7 days at 35-37^o^C, after which OD_600_ was measured using an automated optical plate reader to identify the concentration exhibiting 10% bacterial growth.

MBC assays were set up in the same manner as MIC experiments, except plates were incubated with the drug dilutions for 21 days at 37°C. After incubation, serial dilutions from each well were plated on 7H10 agar supplemented as above with 0.4% activated charcoal. Following colony counting, the MBC_99_ was defined as the lowest compound concentration that produced a 99% reduction in CFU relative to the starting inoculum.

Generation of JNJ-6887, JNJ-1866 and bedaquiline resistant strains

*M. tuberculosis* H37Rv was plated at ~1.20 x 10^8^ CFU onto 7H10 agar plates (10% OADC and 1% glycerol) containing 50x MIC_50_ concentration of either JNJ-1866 or JNJ-6887. Plates were incubated for 3-4 weeks at 37 °C until colonies emerged. Single colonies were restreaked onto fresh 7H10 plates containing 50x MIC_50_ of the same compound and incubated for a further 3 weeks. Individual colonies were then inoculated in 7H9 broth (10% OADC, 0.2% glycerol, 0.05% tween-80) and grown for 3-7 days until visible growth was observed. Resistance was confirmed by MIC assays performed alongside the parental WT strain. A panel of reference antimicrobials was included to assess potential cross-resistance.

Bedaquiline-resistant strains were generated using the same procedure, with selection on 7H10 agar plates containing ~25x MIC_90_ bedaquiline (2.7 µM). After two rounds of selection, single colonies were expanded in 7H9 broth (10% OADC, 0.05% tween-80 and 0.2% glycerol) to an OD_600_ of ~1. Key bedaquiline-resistance genes: *rv0678*, *atpE* and *pepQ* were analysed via Sanger sequencing. Strains harbouring mutations only in *rv0678* were selected for further characterisation. Genomic DNA was extracted for whole genome sequencing and MIC assays were performed to confirm resistance and evaluate cross-resistance relative to the WT parental strain.

Whole genome sequencing and Sanger sequencing

Genomic DNA was isolated from 5 mL cultures of *M. tuberculosis* at OD_600_ 0.9 using a Quick-DNA Fungal/Bacterial Miniprep kit. Whole-genome sequencing was performed by Eurofins. Sequencing reads were aligned to the *M. tuberculosis* H37Rv genome (release 4, 2021-03-23; mycobrowser.epfl.ch) using Bowtie2 and Samtools. BCFtools were used to identify SNPs and insertions/deletions (indels). Delly was used to identify larger genetic deletions and rearrangements and Artemis was used to visualise the data and manually confirm the results

For Sanger sequencing, genes of interest were PCR-amplified using gene-specific primers (**Table S8**) and sequenced at Eurofins using the same primers set.

Timelapse microscopy experiments

*M. tuberculosis* constitutively expressing *Td*Tomato was cultured to mid-exponential phase and seeded into a microfluidic device for imaging, as described previously (*45, 59*). Bacteria were imaged on an inverted fluorescent microscope (Thunder Imaging System, Leica Microsystems) using a x100/1.32 NA oil immersion objective (Leica Microsystems) and a K8 sCMOS camera (Leica Microsystems). Images were acquired in the phase and red channels (555 nm excitation, 590 nm emission) at 1 h intervals for 14-18 days. 7H9 growth medium was perfused through the device at a flow rate of 10-15 µL min^-1^ and exposed to 1 µM JNJ-6887 at the indicated time points. At least 20-30 independent xy-positions were imaged per experiment, and each experiment was performed at least twice. Images were acquired and assembled using the LAS-X software (Leica Microsystems) and analysed using FIJI (*60*).

Generation of genetically modified strains

For the CRISPRi-mediated knockdown strains, small guide RNAs (sgRNAs) targeting *aroG*, *aroK*, *menE* and *menG* were cloned into a plRL2 plasmid using Golden Gate Assembly as described previously (*26*). Briefly, complementary single-stranded oligonucleotides were annealed and ligated into BsmBI-digested plRL2 (Addgene; **Table S8**). Plasmids were sequence-verified and electroporated into *M. tuberculosis* WT and BDQR^Rv0678^ strains. Colonies were selected from 7H10 agar plates containing 50 µg mL^-1^ kanamycin. Knockdown efficiency was assessed by CFU counts over a time-course and by qRT-PCR. Bacterial stocks pre-induced with 100 ng mL^-1^ anhydrotetracycline (ATc) for 5 days (*aroG* and *aroK*) or 7 days (*menE* and *menG*), were diluted to an OD_600_ of 0.0025 in 7H9 broth (with 50 µg mL^-1^ kanamycin; 7H9K) in the presence and absence of 100 ng mL^-1^ ATc. Cultures were incubated at 37°C and CFU counts were obtained on days 3, 7, 10 and 14 by plating serial dilutions onto 7H10 agar containing 0.4% activated charcoal.

Overexpression of *menG*-C146R was achieved using the ATc-inducible pDE43-MCK integrative vector (phage L5 attachment site). Site-directed mutagenesis of the WT *menG* allele was performed using the QuikChange Lightning Kit (Agilent; **Table S8**). Electroporation into WT and colony selection were performed as described above. Cultures were diluted to OD_600_ 0.05 and grown for 5 days with or without 500 ng mL^-1^ ATc before storage. MIC assays using JNJ-6887 were performed to determine ATc-dependent shifts in susceptibility, validating overexpression of the mutant allele.

qRT-PCR

Knockdown strains were grown to mid-log phase and diluted to an OD_600_ of 0.05 in 7H9K, with and without 100 ng mL^-1^ ATc. The empty vector strain was grown in the presence of ATc. After 2-4 days incubation (37°C, 150 rpm), RNA was extracted using the RNeasy kit (Qiagen) and genomic DNA was depleted using Turbo DNAse (Invitrogen). cDNA was generated using Superscript III reverse transcriptase (Invitrogen). Relative transcript knockdown was determined using SYBR green and gene-specific primers (**Table S8**). qRT-PCR was performed using the Applied Biosystems™ 7500 Fast Real-Time PCR System with the following conditions: 50°C for 20 s, 95°C for 10 min, then 40 cycles of 95°C for 10 s, 60°C for 1 min, followed by 95°C for 15 s, 60°C for 1 min, 95°C for 30 s and 60°C for 15 s. Analysis was done by the comparative C(T) method (*61*).

Menaquinone liquid chromatography-mass spectrometry (LC-MS) analyses

For measurement of menaquinone levels following inhibitor treatment (**Fig. 2b**), *M. tuberculosis* cultures (initial OD_600_ = 0.05) were treated with JNJ-6887 (3.33 μM final concentration) or DMSO. After 4 days of incubation (37 ^o^C, rolling), 5 mL of each culture was centrifuged (3,000 g, 10 min), the supernatant removed, and pellets resuspended in 2 mL ultrapure H_2_O. Methanol (6 mL, dry ice-cooled) containing 0.5% formic acid and 1 µM MK-4 internal standard (Sigma-Aldrich) was added. Samples were vortexed and extracted with 6 mL cold petroleum ether (40-60^o^C fraction, Fisher Scientific). Following incubation on dry ice with frequent agitation (45 min), samples were centrifuged (2,000 g, 5 min, 4^o^C) and the upper organic phase transferred to borosilicate glass tubes. The aqueous phase was re-extracted with a further 6 mL of petroleum ether and, following centrifugation as before, the organic phases were combined and evaporated under nitrogen. Samples were stored at -70^o^C. For LC-MS analysis, dried extracts were resuspended in 2 mL of isopropanol:acetonitrile:water (6:3:1). Equal aliquots from each sample were pooled to create a dilution series and identical run-control samples distributed throughout the LC-MS sequence.

LC-MS was performed on an Agilent 1290 Infinity II LC system connected to an Agilent 6545 QTOF mass spectrometer. Chromatography used an Acquity Premier CSH C18 column (Waters; 2.1 x 100 mm, 1.7 μm) at 55 ^o^C. A binary gradient was used with 10 mM ammonium formate in acetonitrile: water (60:40) plus 0.1 % formic acid for mobile phase A and 10 mM ammonium formate in isopropanol: acetonitrile: water (90:9:1) plus 0.1% formic acid for mobile phase B. Flow rate was 400 μL min^-1^ with the percent proportion of mobile phase B changing linearly between times/holds as follows: 0-2 min, 40%; 3 min, 50%; 4 min, 55%; 5-8 min, 70 %; 13-16 min, 95%. All solvents and additives used in metabolite extraction and liquid chromatography were of LC-MS grade quality.

Mass spectrometry was performed using electrospray ionisation (ESI) in positive mode with a nebulizer pressure of 35 psig and a nitrogen drying gas flow rate of 11 L min^-1^ at 320 °C. The capillary, nozzle and fragmentor were set to 3500 V, 1000 V and 175 V, respectively.

Metabolites were identified by exact mass (within 10 ppm of theoretical masses: DMK-9 = 770.6002, MK-9 = 784.6158, MK-9(II-H_2_) = 786.6315) and by MS/MS fragmentation patterns and retention times to MK-4 and MK-9 standards (LKT Laboratories). MK-9(II-H_2_) was distinguished from menaquinol (MKH_2_-9; same exact mass) by the presence of a 187.07 m/z fragment, indicative of the non-reduced naphthoquinone head group present in both MK-4 and MK-9.

Chromatogram alignment, targeted feature extraction and quantification were performed using the Agilent MassHunter software suite. Peak areas were normalised by dividing by the peak area of the MK-4 internal standard and by the OD_600_ for each sample. Metabolites included in the analysis had an r-squared value of > 0.95 in the pooled-sample dilution series and a < 10% relative standard deviation in run-controls.

Menaquinone analysis was independently reproduced at a separate site (**Fig. S8).** *M. tuberculosis* batch cultures treated with JNJ-6887 for 6 days (final concentration 660 nM) were quenched in 6 mL ice-cold 0.2 M HClO_4_ in methanol followed by petroleum ether extraction as above. Extracts were resuspended in 100  μL ethanol and separated on a Thermo Ultimate 3000 UPLC using a C8 column (Supelco Ascentis 100 x 2.1 mm) with a ternary gradient consisting of water, acetonitrile and isopropanol, all containing 0.1 % formic acid (flow rate 200 µL min^-1^). Eluent was analysed on a Thermo Q Exactive mass spectrometer using ESI in positive mode and parallel reaction monitoring (PRM) targeting the MK-9 molecular ion (785.6231, [M+H]+) and the DMK-9 molecular ion (771.6075, [M+H]+). MS/MS fragment spectra were scanned using Skyline set up to analyse the RAW sample files for the presence of the target ions and the resultant fragments characteristic of the naphthoquinone head region for MK-9 (187.0754, 225.0910) or DMK-9 (173.0597, 211.0754) and isoprenoid tail (149.1325). MK-9 and MK-4 standards were run with each batch to confirm system functionality. Organic solvents (Merck) were MS grade. Total ion counts (TIC) for DMK-9 and MK-9 were normalised to CFU values determined by plating 100 µL of JNJ-6887 treated culture on 7H10 agar.

LC-MS analysis of *aroK* CRISPRi strain

A primary culture of *aroK* ‘high’ knockdown strain was grown mid-log phase and subcultured to OD_600_ of ~0.1 in 7H10 broth supplemented with 50 µg mL^-1^ kanamycin, with or without 5 µg mL^-1^ ATc. After 5 days of incubation (37°C at 150 rpm), samples were collected for metabolite extraction alongside OD_600_ and CFU measurements. MK-9 extraction and analysis were performed as described above. For shikimate measurement, the 8 mL aqueous fraction remaining after the petroleum ether extraction was neutralised with 350 μL of 3 M ammonia solution and mixed with 3.4 mL acetonitrile before storage at -70 ^o^C. Samples were thawed and incubated at -20°C for 2 h, vortexed and centrifuged (2,000 g, 10 min, 4 ^o^C). Supernatants (750 μL) were filtered through prewashed Spin-X 0.22 μm centrifuge filters (Corning) by centrifuging for 15,000 g for 10 min at 4°C. Filtrates were diluted 1:1 with acetonitrile:methanol:water (4:4:2), then further diluted 1:1 with 0.2% acetic acid in acetonitrile. Samples were incubated on ice for 10 min, centrifuged (15,000 g, 10 min) and supernatants transferred to autosampler vials.

Shikimate was analysed on an Agilent 1290 Infinity II LC system coupled to an Agilent Accurate Mass 6545 QTOF mass spectrometer. Chromatography was performed using a Cogent Diamond Hydride Type C silica HPLC column (MicroSolv) at 25 ^o^C. A binary gradient was used with 0.2% acetic acid in water for mobile phase A and 0.2% acetic acid in acetonitrile for mobile phase B. The flow rate was 400 μL min^−1^ with the solvent gradient changing from 85% mobile phase B to 20 % mobile phase B over 25 min, before returning to 85% mobile phase B for 5 min re-equilibration. For MS, ionization was performed using ESI in negative mode with nebulizer pressure of 50 psig and a nitrogen drying gas flow rate of 5 L min^−1^ at 300 °C. The capillary, nozzle and fragmentor voltages were set to 1,500 V, 2,000 V and 100 V, respectively. Shikimate peak heights were normalised to sample OD_600_.

Enzymatic assay

*Membrane preparation from M. tuberculosis mc^2^ 6230: M. tuberculosis* mc^2^ 6230 was grown in 7H9 broth (supplemented with 0.5% (v/v) oleic acid, 0.5% (w/v) albumin, 0.2% (w/v) dextrose, 24 μg mL^-1^ D-pantothenate and 0.2% casamino acids). Washed cells were resuspended in Buffer A (50 mM MOPS pH 7.9, 5 mM MgCl_2_, 5 mM DTT, 10% glycerol [v/v]), at 2 mL g^-1^ of cells, and disrupted by probe sonication on ice with a Sanyo Soniprep 150 (10 cycles of 60 s on and 90 s off). The whole cell lysates were centrifuged (27,000 g, 20 min, 4 °C). The supernatant was further centrifuged (100,000 g, 30 min, 4 ^o^C) in an Optima TLX Ultracentrifuge (Beckman). The membrane-enriched pellets were washed (resuspended) in Buffer A followed by ultracentrifugation at 100,000 g. The washed pellets were resuspended in Buffer A, divided into aliquots and frozen at -80 °C. The protein concentration of the membrane-enriched fraction was estimated using a BCA protein assay kit (Pierce).

*Partial purification and quantification of DMK-8 substrate from E. coli ΔubiE mutant: E. coli* Δ*ubiE* mutant cells were grown in Lysogeny Broth (LB) in the presence of kanamycin (50 µg mL^-1^) at 37°C. Cells were harvested by centrifugation and washed with water. The cell pellets were extracted twice with 20 volumes of chloroform/methanol (2:1, v/v), and the combined extracts were washed with 1/5 vol of water. The upper aqueous phase was discarded, and the lower organic phase was completely evaporated under a gentle nitrogen stream. The lipids were re-dissolved in chloroform and applied to a 100−200 mesh silicic acid (Sigma) column pre-equilibrated in the same solvent. Neutral lipids containing partially purified DMK were eluted with chloroform, the solvent was evaporated under nitrogen stream, and the sample was re-dissolved in ethanol (100%) for subsequent LC-MS analysis. LC-MS was performed using an atmospheric pressure photo-ionization source in positive mode on an Agilent 6220 time-of-flight mass spectrometer. Chromatographic separation of the lipids was accomplished using a gradient of 100% solvent A (methanol) to 50% solvent B (isopropyl alcohol) on a Waters X-Bridge (C18, 2.1 mm×150 mm, 3.5 μm particle size) column heated to 40°C at a flow rate of 0.3 mL min^-1^. The drying gas temperature was 350°C, the vaporizer temperature was 300°C, and the fragmentor voltage was set to 120 V. Mass spectra were acquired from m/z 110 to 1800 with a frequency of one scan/sec. LC-MS analysis was also performed on vitamin K_1_, K_2_ (MK-4) and MK-9 standards; concentration curves were constructed and used to quantitate DMK-8, which was ultimately used as a substrate in enzyme assays.

*Radiochemical M. tuberculosis MenG inhibition assay:* Assay mixtures (100 μL) contained 100 mM Tris-HCl pH 7.4, 1 mM DTT, 5 mM MgCl_2_, 0.1% CHAPS, 500 ng of DMK-8, 40 μM radiolabelled SAM, and inhibitors at the indicated concentrations. Specific activity of the [^14^C]SAM was 52.6 mCi mmol^-1^. Compounds with MIC_50_ values determined to be >12 µM were screened at 10 µM, those with MIC_50_ values <12 µM and >5 µM inhibitor were screened at 5 µM and those with MIC_50_ values < 5 µM inhibitor were screened at 0.02 and 0.0012 µM. In all experiments, we included ‘no inhibitor’ and positive controls. Reactions were initiated by the addition of 50–100 μg of *M. tuberculosis* membrane protein and incubated at 37 °C for 1 h. Reactions were stopped by the addition of 0.1 M acetic acid in methanol (0.5 mL), and radiolabelled products were extracted with hexane (2 × 3 mL). Pooled extracts were washed with 1 mL of water, evaporated to dryness under a N_2_ stream, and dissolved in CHCl_3_/CH_3_OH (2:1, v/v). An aliquot was subjected to liquid scintillation counting (LS 6500, Beckman Coulter); a second aliquot and authentic standards (DMK-8 and MK-9) were subjected to reverse-phase thin-layer chromatography (TLC) developed in acetone/water (97:3). Standards were visualised under UV light, and distribution of radioactivity was detected by phosphorimaging (Azure Bioanalytical system) and quantified with ImageQuant TL v2005 software (Amersham Biosciences). The use of radioisotopes in screening assays greatly increases assay time and costs, therefore an efficient method of compound testing was developed as detailed previously (*62*).

*In vivo* animal work

Six-to-eight week-old female Balb/cBy mice were purchased from Charles River Laboratories (France) or Janvier (France). Mice were housed in individually ventilated cages in HEPA-filtered racks (IVC cages: 1291H – IVS 800 cm^2^ or ER1050 – IVC 1050 cm^2^, Techniplast), in a 12-hour light-dark cycle and with access to water and food *ad libitum*. An ambient temperature of 22±2˚C, a relative humidity of 55±10% and a negative pressure of -20Pa were maintained. All mice were allowed to acclimatise for at least 5 days.

Short acute mouse model

Mice were infected intranasally with either 1,000 CFU (**Fig. 1d**) or 200 CFU (**Fig. S24**) per mouse with *M. tuberculosis* H37Rv WT or BDQR^Rv0678^ strains. To verify infection levels, a subgroup of six mice was euthanised one day after infection. Mice were allowed to establish infection for 7 days before treatment was initiated, and were treated daily for 12 consecutive days. Mice were euthanised 3 days after the last PO dose to prevent compound carryover. Additional control groups were euthanised at day 7 (treatment start) and day 21 (end of treatment). Compounds were administered by oral gavage (100 µL, drencher with rounded end straight, 0.9mm x 25mm, Socorex Swiss) except for the long-acting formulation, which was administered subcutaneously in the upper back (100 µL) using a 25G x 16mm needle (BD Microlance™).

CRISPRi-mediated knockdown mouse model

Three to five days before infection, mice assigned to knockdown-induction groups were provided doxycycline-containing food (A04 2g kg^-1^ Doxycycline Hyclate +BLUE, E8220P01R00569, Tecnilab-BMI). Control groups received the standard diet. Mice were infected intranasally with 500 CFU of CRISPRi knockdown strain in either *M. tuberculosis* H37Rv WT or BDQR^Rv0678^ backgrounds. Bacterial strains targeting *menE* and *menG* were pre-induced for 7 days with 100 ng mL^-1^ ATc, whereas *aroG*, *aroK* and all control arms were not pre-induced. To verify infection levels, a subgroup of five mice per strain was euthanised one day after infection. One day after infection, subgroups of mice were treated with 1.5, 6.5, or 25 mg kg^-1^ of bedaquiline by oral gavage (100 µL, Instech flexible tip 18GA x 30 mm). Subgroups of five mice were euthanized at 7, 14, 21 and 26 (*aroG* and *aroK*) or 28 (*menE* and *menG*) days after infection. During euthanasia on days 7 and 26 or 28, the two upper right lobes were collected, snap-frozen in liquid nitrogen and stored at -80˚C for RNA extraction.

Enumeration of CFU from lung

At sacrifice, whole lungs were aseptically collected in gentleMACS™ tubes (M tubes with strainer, Miltenyi biotec) containing 2.5 mL of phosphate-buffered saline (PBS) and homogenised using the “RNA_01_01” settings of gentleMACS™ Octo Dissociator (Miltenyi Biotec). Lung homogenate was diluted in PBS and plated in 7H10 charcoal agar plates containing antibiotics (amphotericin: 100 µg mL^-1^ Polymyxin B: 25 µg mL^-1^, Carbenecillin: 50 µg mL^-1^, Trimethoprim: 20 µg mL^-1^). Plates were incubated at 37°C for 3- 5 weeks before CFU counts were recorded. Data are presented as the mean log_10_ CFU lung^-1^ for each group. Statistical analyses were performed using one-way ANOVA with Sidak’s test for multiple comparisons (GraphPad Prism).

Microsampling bleeding for pharmacokinetic (PK) analysis

The mice were warmed to dilate the blood vessel in a warm chamber at 37°C for 10 min prior to the blood collection. Mice should be properly restrained, leaving the tail free. The microsampling was performed in the lateral tail vein. A lancet was used to bleed the mice. Samples of 50 µL were taken with a microcapillary coated with Ethylene diamine tetraacetic acid (EDTA; Vitrex) attached to the injury to allow blood to fill the capillary. After collection, the capillary was sealed with wax and placed on ice. Samples were centrifuged (3000 g, 10 min), then 10 µL of plasma was collected using an end-to-end capillary (Vitrex). To decontaminate the sample, 200 µL of acetonitrile was used for further PK analysis.

THP-1 derived macrophage model

THP-1 monocyte cells (ATCC TIB-202) cultured in RPMI with 10% heat-inactivated foetal bovine serum (hiFBS; Gibco) for 3-4 weeks until cells were free of clumps and ready for infection. Cells (2.5 x 10^5^) were added to each well of a 24-well plate in media containing 100 nM phorbol 12-myristate 13-acetate (PMA; Merck). Cells were allowed to adhere at 37°C 5% CO_2_ for 48 h before washing with PBS and replacement with RPMI and 10% hiFBS. After 24 h incubation at 37°C 5% CO_2_ cells were infected with BDQR^Rv0678^ at an MOI of 0.4. After 4 h incubation with the bacteria, media was removed and replaced with media containing gentamicin 50 µg mL^-1^ for 1 h at 37°C. Media was removed and cells washed three times with PBS before addition of drug treatments in RPMI with 10% hiFBS. Day 0 counts were retained for lysis with remaining conditions incubated for 72 h at 37°C, 5% CO_2_. At Day 0 or Day 3, cells were incubated for 10 min with 0.2% triton-X, serially diluted and plated onto 7H10 agar with 0.4% activated charcoal. CFU enumeration was performed after 3 weeks incubation at 37°C.

Oxygen consumption rate (OCR) assay

*M. tuberculosis* was grown in 7H9 media (Difco) supplemented with 10% OADC (Difco) and 0.01% Tyloxapol at 37 °C. After reaching OD_600_ of 0.6 the culture was passaged to an OD_600_ of 0.05. JNJ-6887 was added to the culture to final concentration of 10 x MIC_50_ (0.661 µM). After 3 days of inhibitor exposure, aliquots of *M. tuberculosis* culture were taken for CFU enumeration and optical density measurements. Remaining cultures were prepared for extracellular flux assay as previously described (*63*).

The OCR of *M. tuberculosis* bacilli adhered to the well bottom of a Cell-Tak (Corning) coated XF cell culture microplate (Seahorse Biosciences), at 4x10^6^ bacilli per well, was measured using a XFe96 Extracellular Flux Analyser (Seahorse Biosciences). Cell-Tak has a negligible effect on *M. tuberculosis* basal respiration (*63*). Assays were carried out in unbuffered 7H9 broth (pH 7.35) without carbon source. Basal OCR was measured for ~ 25 min before the sequential, automatic addition of bedaquiline (final concentration of 32.4 µM) and carbonyl cyanide m-chlorophenyl hydrazone (CCCP; final concentration of 4 µM) respectively through the drug ports of the sensor cartridge. The addition of CCCP causes an uncoupling of the proton gradient reducing the functional capacity of ATP synthase. After the addition of CCCP, to stimulate maximum respiration, OCR was measured for an additional 14 min. Vertical dotted lines in OCR profile indicate addition of bedaquiline and CCCP. OCR measurements are representative of a minimum of five replicate wells with assay time point data representative of the average OCR of a 4-min measurement. OCR data point averages were calculated using Wave Desktop software (V2.6.0.31, Seahorse Biosciences). P-values were determined by a multiple paired t-test using GraphPad Prism 10.2.0. Two independent experiments were conducted with each OCR assay including a minimum of two biological replicates for the JNJ-6887 treated *M. tuberculosis*. Bedaquiline and JNJ-6887 were obtained from Janssen Pharmaceuticals (Beerse, Belgium). All other compounds were obtained from Sigma-Aldrich unless otherwise stipulated.

Measurement of membrane potential

Membrane potential was measured in mycobacteria using a method previously published (*64*). Briefly, *M. tuberculosis* cultured to mid-exponential phase (OD_600_ 0.3-0.6) were harvested and resuspended in PBS containing 0.05% Tween-80 and 30 µM DiOC_2_(3) (Invitrogen). Bacteria were incubated at 37°C, in an orbital shaker (100 rpm) for 90 min. The bacterial cultures were then washed with PBS to remove excess DiOC_2_(3) and resuspended in 500 µL of PBS containing 0.05% Tween-80. Samples (198 µL) were dispensed into individual wells of a 96-well plate with 2 μL of either 100% ethanol, CCCP (Sigma Aldrich) or JNJ-6887 (final concentrations: 1%, 25 μM and 1 μM, respectively). Fluorescence was measured using a SpectraMax i3x plate reader (excitation: 485 nm, emission: 520 nm, 620 nm) at 37 °C after 1 h of exposure. Experiments were carried out three times each with at least two biological replicates.

Vitamin K_2_ rescue assay

A concentrated stock solution of vitamin K_2_ (0.1 mg mL^-1^; Sigma, V9378) was prepared. *M. tuberculosis* H37Rv (final concentration 5 x 10^5^ CFU mL^-1^) was treated with 0.15 or 3.33 µM JNJ-6887 and a dilution series of vitamin K_2_ (40 ng mL^-1^ to 0.15 ng mL^-1^) in 7H9 supplemented with 10% OADC and 0.05% Tween-80. Plates were incubated at 37 ºC for 17 days. For each condition, 5 µL was stamped onto a 7H10 charcoal agar plate. Plates were incubated at 37 ºC for 3-4 weeks to allow CFU enumeration.

Checkerboard assay

To assess the combined effects of two compounds on *M. tuberculosis* growth, 500 nL of each two-fold serially diluted compound was spotted into 96-well plates in a prespecified checkerboard layout. Bacterial inoculum was prepared at 5.0 x 10^5^ CFU mL^-1^ in 7H9 media supplemented with 10% OADC and 0.05% tween, and 200 µL was added to each well. Plates were incubated for 10 days at 37^o^C. On Days 7 and 10, OD_620_ was determined using a Perkin Elmer Envision Reader. On Days 0, 7 and 10, 5 µL from each well was stamped onto 7H10 agar plates (96-well format), allowed to air dry and incubated inverted at 37°C for 14 days. Plates were imaged and bacterial growth was scored manually.

Time-kill kinetics

For chemical inhibition of MenG, mid-log phase *M. tuberculosis* cultures were adjusted to ~5 x 10^5^ CFU mL^-1^ in 7H9 media and added to 96-well plates containing DMSO or compounds pre-diluted to 100x stock concentrations. At 7, 14 and 21 days, wells were sampled and CFU enumerated by serial dilution and plating onto 7H10 agar supplemented with 0.4% activated charcoal.

Time-kill kinetics using the CRISPRi-mediated knockdown strains were performed similarly except that bacterial stocks were diluted to an initial OD_600_ of ~0.0025 in 7H9 media with or without 100 ng mL^-1^ ATc. For *menE* and *menG* CRISPRi strains, ATc was replenished every 3-4 days to maintain knockdown (equivalent DMSO was added to all ATc-negative wells). At 7, 10, 14 and 21 days, samples were collected and CFU enumerated as described above.

Computational modelling

Since no X-ray crystal structure of TB MenG is available, an AlphaFold model was used to get some structural insight into the possible binding mode of the inhibitors. In the generated AlphaFold model (*31*), a SAM molecule was modelled based on a SAM-bound crystal structure of Yeast Coq5, a methyltransferase with 35% sequence similarity with *Mt*MenG (PDB: 4OBW) (*30*). In the obtained model a putative ligand binding pocket is present close to the resistance mutations D25G or D25H, N28D, and S32P. A model of JNJ-6887 in this binding pocket was obtained by IFD-MD (Schrödinger Release 2021-3: IFD-MD, Schrödinger, 2021) and other ligands (JNJ-8833 and JNJ-1866) were docked using Glide (Schrödinger Release 2021-3: Glide, Schrödinger, 2021).

Generation of a bedaquiline-resistant *M. marinum* strain

*M. marinum* was cultured as previously described (*65*). To generate a bedaquiline-resistant *M. marinum* strain, *M. marinum* ‘M’ strain (1 x 10^9^ CFU mL^-1^; pMSP12::Wasabi (*66*)) in 7H9 supplemented with 10% OADC, 0.2% glycerol, 0.05% Tween-80 and 50 µg mL^-1^ hygromycin, was spread onto 7H10 agar plates containing 0.3 µM (3x MIC_90_) bedaquiline at 33^o^C. Retention of the plasmid containing the wasabi fluorescent protein in resistant colonies was confirmed with a UV transilluminator. Individual colonies were grown in 7H9 broth containing 0.3 µM bedaquiline to confirm resistance and frozen in 7H9 broth supplemented with 10% OADC and 5% glycerol. Genomic DNA was extracted using the Quick-DNA Fungal/Bacterial Miniprep Kit based on the manufacturer’s protocol (Zymo Research) and the MMAR_1007 locus (501 bp) was sequenced (Full Circle Labs; **Table S8**). The bedaquiline-resistant strain was found to have a point mutation (C325T) which introduced a stop codon within the open reading frame.

Zebrafish husbandry, injections, drug treatment, imaging and burden analyses

Adult wild-type AB zebrafish were housed in the Biological Services Facility at LSHTM. Embryos were obtained from naturally spawning zebrafish, and larvae were maintained at 28.5^o^C in embryo medium (1x E3 medium). From 1 day post fertilisation (dpf), embryos were maintained in E3 media with 1-phenyl 2-thiourea (PTU; 0.036 g L^-1^) to inhibit melanogenesis and improve optical transparency for imaging.

Briefly, 50–100 mL *M. marinum* was grown at 33°C in 7H9 broth supplemented with 10% OADC, 0.2% glycerol, 0.05% Tween-80 and 50 μg mL^-1^ hygromycin. The supernatant was passed through a 27G needle 10 times and centrifuged (100 g, 1 min) then resuspended as single-cell bacterial suspensions in 1 mL freezing media (7H9 supplemented with 0.2% glycerol only). This was passed through a 5 µm filter and stored at −80°C. CFU was quantified via plating on 7H10 agar grown at 33°C.

At 48-hours post fertilisation (hpf), wild-type AB zebrafish embryos were decorionated and anesthetised with tricaine (200 μg mL^-1^; Sigma-Aldrich) in E3. Embryos were infected using thawed single-cell suspensions diluted in a solution of 4% polyvinyl-pyrrolidone (Sigma-Aldrich) in PBS and 0.5% phenol red (Sigma-Aldrich) to obtain an infection dose of 200 CFU nl^-1^. For injection, 1-2 nL bacterial suspension was microinjected via the caudal vein (specific infection dose indicated in figure legend). Embryos were recovered in E3 media + PTU and maintained at 28°C. Embryos were randomly assigned to groups and treated with various concentrations of JNJ-1866 or bedaquiline prepared in DMSO. Control embryos were treated with DMSO only. The duration of treatment is indicated in the relevant figure legend. Live imaging was performed on anaesthetised embryos suspended in 3% methylcellulose. Images were acquired using a Leica M205FA Fluorescent Stereo Microscope equipped with a Leica DFC365FX monochrome digital camera (Leica Microsystems). Images were analysed using ImageJ software to quantify the fluorescent pixel count, defined as fluorescent signal above a consistent set background determined empirically for each experimental dataset (*65*). Data are presented as total fluorescent area (pixels) above the background level. All statistical analyses were carried out using GraphPad Prism Software version 9.4.1 (GraphPad, USA). Statistical significance was tested using the Kruskal-Wallis test with Dunn’s multiple comparisons test.

### Supplementary Text

Chemical synthesis

**1: NMR instruments:**

NMR experiments were performed using a Bruker BBFO ASCEND^®^ 400 AVANCE III HD 400 MHz and Bruker BBFO ULTRASHIELD^®^ 300 AVANCE III HD 300 MHz. Internal standard: Tetramethylsilane (TMS).

**2: General Analytical LCMS Methods**

Instrument: analyses were conducted with a Shimadzu LCMS-2020 with electrospray ionization in positive ion detection mode with a 40 ADXR (20AD & 20ADXR) pump, SIL-40 ACXR (SIL-20ACXR) autosampler, CTO-40AC (CTO-20AC) column oven, M40A (M20A) PDA Detector and LCMS 2020 LCMS detector.

The LCMS detector was configured with electrospray ionization as ionizable source; Acquisition mode: Scan; Nebulizing Gas Flow: 1.5 L/min; Drying Gas Flow: 15 L/min; Detector Voltage: 0.95 - 1.25 kv; DL Temperature: 250 °C; Heat Block Temperature: 250 °C; Scan Range: 90.00 - 900.00 m/z.

**Synthesis of JNJ-8833**

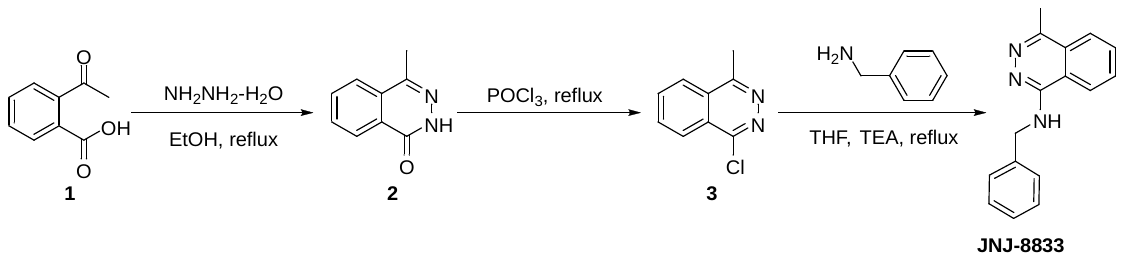

**JNJ-8833: *N*-benzyl-4-methylphthalazin-1-amine**

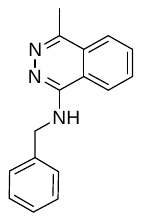

Step 1. 4-Methylphthalazin-1(2*H*)-one (**2**). 2-Acetylbenzoic acid (**1**, 5.00 g, 30.5 mmol), hydrazine monohydrate (1.83 g, 36.6 mmol), a stir bar, and EtOH (100.0 mL) were added to an oven-dried 250 mL round-bottomed flask. The resulting mixture was stirred at 90 ^o^C overnight. After cooled to rt, the solid was collected by filtration, washed with EtOH and dried in air to give 4-methylphthalazin-1(2*H*)-one (**2**, 3.70 g, 74% yield) as a white solid. LCMS (ESI): mass calcd. for C_9_H_8_N_2_O, 160.1; m/z found, 161.1 [M+H]^+^.

Step 2: 1-Chloro-4-methylphthalazine (**3**). 4-Methylphthalazin-1(2*H*)-one (**2**, 3.70 g, 23.1 mmol), a stir bar, and POCl_3_ (80.0 mL) were added to an oven-dried 250 mL round-bottomed flask. The resulting mixture was stirred at 110 ^o^C for 2 h. After cooled to rt, the volatiles were removed in vacuo and the residue was re-dissolved in DCM, then adjust the pH of mixture with sat. K_2_CO_3_ solution at 0 ^o^C. The mixture was extracted with DCM, dried over Na_2_SO_4_, and concentrated i*n vacuo* to give 1-chloro-4-methylphthalazine (**3**, 3.50 g, 84% yield) as a yellow solid. LCMS (ESI): mass calcd. for C_9_H_7_ClN_2_, 178.0/180.0; m/z found, 179.3/181.3 [M+H]^+^.

Step 3: *N*-Benzyl-4-methylphthalazin-1-amine (**JNJ-8833**). 1-Chloro-4-methylphthalazine (100.0 mg, 0.5600 mmol), phenylmethanamine (120.0 mg, 1.120 mmol), TEA (170.0 mg, 1.680 mmol), a stir bar, and THF (2.00 mL) were added to an 8 mL vial. The mixture was stirred at 80 ^o^C overnight. After cooled to rt, the solvent was removed *in vacuo* and the residue was subject to *prep*-HPLC (Column: XBridge Prep OBD C18 Column, 30 × 150 mm, 5 μm; Mobile Phase A: water (10 mmol/L NH_4_HCO_3_ + 0.1% NH_3_), Mobile Phase B: ACN; Flow rate: 60 mL/min; Gradient: 18 to 50% B in 7 min; 254/220 nm; Rt_1_ (min): 6.6 min) to give *N*-benzyl-4-methylphthalazin-1-amine as a white solid (**JNJ-8833**, 32.8 mg, 24% yield

HRMS (ESI+) *m/z* [M + H]^+^ calculated for C_16_H_16_N_3_: 250.1266; found: 250.1348. ^1^H NMR (400 MHz, DMSO-*d*_6_) δ ppm 8.30 - 8.41 (m, 1 H) 7.95 - 8.04 (m, 1 H) 7.81 - 7.93 (m, 3 H) 7.38 (d, *J*=7.2 Hz, 2 H) 7.29 (t, *J*=7.4 Hz, 2 H) 7.15 - 7.25 (m, 1 H) 4.77 (d, *J*=6.0 Hz, 2 H) 2.66 (s, 3 H). ^13^C NMR (101 MHz, DMSO-*d*_6_) δ ppm 152.89 (s, 1 C) 147.89 (s, 1 C) 140.46 (s, 1 C) 131.47 (s, 1 CH) 130.96 (s, 1 CH) 128.07 (s, 2 CH) 127.18 (s, 2 CH) 126.38 (s, 1 CH) 126.29 (s, 1 C) 124.65 (s, 1 CH) 122.19 (s, 1 CH) 117.78 (s, 1 C) 44.00 (s, 1 CH2) 19.01 (s, 1 CH3)

**Synthesis of JNJ-2842**

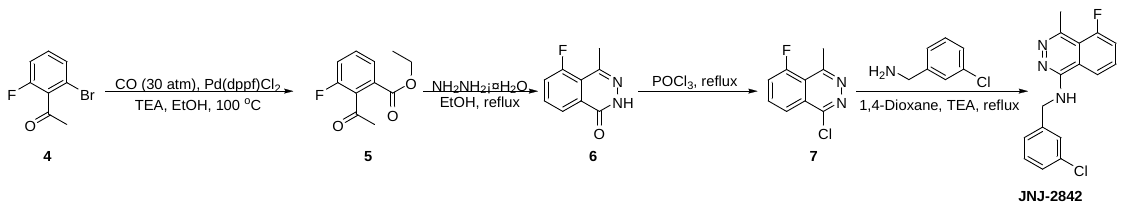
**JNJ-2842: *N*-(3-chlorobenzyl)-5-fluoro-4-methylphthalazin-1-amine**

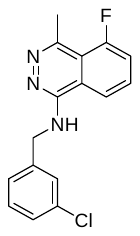

Step 1: Ethyl 2-acetyl-3-fluorobenzoate (**5**). 1-(2-Bromo-6-fluorophenyl)ethanone (**4**, 2.00 g, 9.22 mmol), EtOH (20.0 mL), TEA (2.80 g, 27.6 mmol), a stir bar, and Pd(dppf)Cl_2_ (674.0 mg, 0.9220 mmol) were added to a 30 mL sealable metal container equipped with a gas inlet pressure gauge. The vessel was pressurized with carbon monoxide at 30 atm and stirred at 100 °C overnight. After cooled to rt, the solvent was removed *in vacuo* and the residue was subjected to silica gel column (0-50% ethyl acetate/petroleum ether) to give ethyl 2-acetyl-3-fluorobenzoate as a yellow oil (**5**, 1.30 g, 66% yield). LCMS (ESI): mass calcd. for C_11_H_11_FO_3_, 210.0; m/z found, 211.2 [M+H]^+^.

Step 2: 5-Fluoro-4-methylphthalazin-1(2*H*)-one (**6**). Ethyl 2-acetyl-3-fluorobenzoate (**5**, 1.30 g, 6.19 mmol), hydrazine hydrate (310.0 mg, 6.185 mmol) and EtOH (20.0 mL) were added to a 50 mL flask. The reaction mixture was stirred at 80 ^o^C for 3 h. After cooled to rt, the solid was collected by filtration and dried in air to give 5-fluoro-4-methylphthalazin-1(2*H*)-one as a white solid (**6**, 740.0 mg, 67% yield). LCMS (ESI): mass calcd. for C_9_H_7_FN_2_O, 178.1; m/z found, 179.2 [M+H]^+^.

Step 3: 1-Chloro-5-fluoro-4-methylphthalazine (**7**). 5-Fluoro-4-methylphthalazin-1(2*H*)-one (**6**, 740.0 mg, 4.153 mmol), a stir bar, and POCl_3_ (20.0 mL) were added to a 40 mL sealed tube. The reaction mixture was stirred at 110 ^o^C for 1 h. After cooled to rt, the mixture was concentrated *in vacuo*, the residue was dissolved in DCM (50 mL) and adjusted pH to 8 with K_2_CO_3_. The resulting mixture was separate with dichloromethane (3 x 50 mL) and water (50 mL). Then the organic layers were combined and concentrated to give 1-chloro-5-fluoro-4-methylphthalazine as a red solid (**7**, 700.0 mg, 86% yield). LCMS (ESI): mass calcd. for C_9_H_6_ClFN_2_, 196.0; m/z found, 197.2 [M+H]^+^.

Step 4: *N*-(3-chlorobenzyl)-5-fluoro-4-methylphthalazin-1-amine. 1-Chloro-5-fluoro-4-methylphthalazine (**7**, 300.0 mg, 1.526 mmol), 1,4-dioxane (1.00 mL), (3-chlorophenyl)methanamine (1.08 g, 7.63 mmol), a stir bar, and TEA (772.0 mg, 7.629 mmol) were added to an 8 mL sealed tube. The reaction mixture was stirred at 110 ^o^C for 2 hours. After cooled to rt, the volatiles were removed *in vacuo* and the residue was subjected to *prep*-HPLC (Column: XBridge Prep OBD C18 Column, 30 × 150 mm, 5 μm; Mobile Phase A: water (10 mmol/L NH_4_HCO_3_ + 0.1% NH_3_), Mobile Phase B: ACN; Flow rate: 60 mL/min; Gradient: 20 to 75% B in 7 min; 254/220 nm; Rt_1_ (min): 6.65) to give *N*-(3-chlorobenzyl)-5-fluoro-4-methylphthalazin-1-amine as a light yellow solid (**JNJ-2842**, 74.6 mg, 16% yield

HRMS (ESI+) *m/z* [M + H]^+^ calculated for C_16_H_14_ClFN_3_: 302.0782; found: 302.08658. ^1^H NMR (400 MHz, DMSO-*d*_6_) δ ppm 8.15 (dd, *J*=8.3, 0.7 Hz, 1 H) 8.03 (t, *J*=5.9 Hz, 1 H) 7.91 (td, *J*=8.1, 5.3 Hz, 1 H) 7.70 (ddd, *J*=12.0, 8.1, 0.7 Hz, 1 H) 7.40 - 7.47 (m, 1 H) 7.22 - 7.38 (m, 3 H) 4.75 (d, *J*=5.8 Hz, 2 H) 2.75 (d, *J*=7.2 Hz, 3 H). ^13^C NMR (101 MHz, DMSO-*d*_6_) δ ppm 159.19 (s, 1 C) 156.67 (s, 1 C) 151.70 (d, *J*=4.2 Hz, 1 C) 144.35 (d, *J*=5.7 Hz, 1 C) 142.95 (s, 1 CH) 132.81 (s, 1 C) 132.32 (s, 1 CH) 132.23 (s, 1 CH) 130.01 (s, 1 CH) 125.64 - 127.54 (m, 1 C) 119.69 (d, *J*=4.2 Hz, 1 C) 118.54 (d, *J*=3.5 Hz, 1 CH) 117.66 (s, 1 CH) 117.44 (s, 1 CH) 43.67 (s, 1 CH2) 23.10 (d, *J*=9.2 Hz, 1 CH3). ^19^F NMR (376 MHz, DMSO-*d*_6_) δ ppm -110.21 (s, 1 F).

**Synthesis of JNJ-6887**

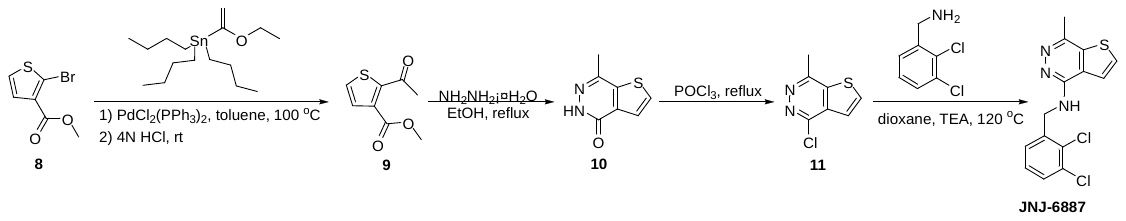

**JNJ 6887: *N*-(2,3-dichlorobenzyl)-7-methylthieno[3,2-*d*]pyridazin-4-amine**

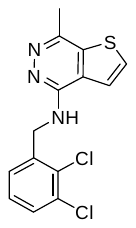

Step 1: Methyl 2-acetylthiophene-3-carboxylate (**9**). Methyl 2-bromothiophene-3-carboxylate (**8**, 9.00 g, 40.7 mmol), tributyl(1-ethoxyvinyl)stannane (29.41 g, 81.42 mmol), Pd(PPh_3_)_2_Cl_2_ (2.86 g, 4.07 mmol), a stir bar, and toluene (135 mL) were added to a 250 mL round bottomed flask. The resulting mixture was maintained under nitrogen and stirred at 100 °C for 16 hours. After cooling down to rt, the mixture was filtered and concentrated *in vacuo*. The residue was dissolved in 4N HCl (50 mL). The reaction was stirred at rt for 1 hour, extracted with ethyl acetate (3 x 100 mL), the organic layers were combined, washed with brine (200 mL), dried over anhydrous sodium sulfate, filtered and concentrated. The residue obtained was subjected to silica gel chromatography (15-20% ethyl acetate/petroleum ether) to afford methyl 2-acetylthiophene-3-carboxylate as a yellow oil (**9**, 7.00 g, 93% yield). LCMS (ESI): mass calcd. for C_8_H_8_O_3_S, 184.0; m/z found, 185.2 [M+H]^+^.

Step 2. 7-Methylthieno[3,2-*d*]pyridazin-4(5*H*)-one (**10**). Methyl 2-acetylthiophene-3-carboxylate (**9**, 6.00 g, 32.6 mmol), hydrazine hydrate (3.84 g, 65.1 mmol), a stir bar, and ethanol (60.0 mL) were added to a 250 mL round bottomed flask. The resulting mixture was stirred at 80 °C for 16 hours. After cooling down to rt, the product was collected by filtration and dried in air to give 7-methylthieno[3,2-*d*]pyridazin-4(5*H*)-one as a white solid (**10**, 5.20 g, 96% yield). LCMS (ESI): mass calcd. for C_7_H_6_N_2_OS: 166.0; m/z found, 167.0 [M+H]^+^.

Step 3: 4-Chloro-7-methylthieno[3,2-*d*]pyridazine (**11**). 7-Methylthieno[2,3-*d*]pyridazin-4(5*H*)-one (**10**, 5.20 g, 31.3 mmol), a stir bar, POCl_3_ (50.0 mL) were added to a 100 mL round bottomed flask. The resulting mixture was stirred at 100 °C for 3 hours. After cooled to rt, the mixture was concentrated *in vacuo*, the residue was dissolved in DCM (200 mL) and the pH was adjusted to 8 with K_2_CO_3_. The resulting mixture was separate with DCM (3 x 200 mL) and water (200 mL). Then the organic layers were combined and concentrated to give 4-chloro-7-methylthieno[3,2-*d*]pyridazine as a light pink solid (**11**, 5.70 g, 99% yield). LCMS (ESI): mass calcd. for C_7_H_5_ClN_2_S: 184.0; m/z found, 185.1 [M+H]^+^.

Step 4: *N*-(2,3-dichlorobenzyl)-7-methylthieno[3,2-*d*]pyridazin-4-amine (**JNJ-6887**). 4-Chloro-7-methylthieno[2,3-*d*]pyridazine (**11**, 200.0 mg, 1.083 mmol), (2,3-dichlorophenyl)methanamine (572.0 mg, 3.249 mmol), TEA (548.0 mg, 5.416 mmol), a stir bar, and 1,4-dioxane (1.00 mL) were added to an 8 mL vial. The resulting mixture was stirred at 120 °C for 2 d. After cooling down to rt, the reaction was quenched with water (10 mL). The resulting mixture was extracted with ethyl acetate (3 x 10 mL), the organic layers were combined, washed with brine (1 x 20 mL), dried over anhydrous sodium sulfate, filtered and concentrated to afford a yellow oil. The yellow oil was subjected to *prep*-HPLC (Column: YMC-Actus Triart C18, 30 mm x 150 mm, 5 μm; Mobile Phase A: water (10 mmol/L NH_4_HCO_3_ + 0.1% NH_3_), Mobile Phase B: ACN; Flow rate: 60 mL/min; Gradient: 45 to 65% B in 7 min; 254 nm; Rt_1_(min):7.73) to afford *N*-(2,3-dichlorobenzyl)-7-methylthieno[2,3-*d*]pyridazin-4-amine as a white solid (**JNJ-6887**, 48.3 mg, 13% yield).

HRMS (ESI+) *m/z* [M + H]^+^ calculated for C14H11Cl2N3S: 324.0051; found: 324.0133. ^1^H NMR (400 MHz, DMSO-*d*_6_) δ ppm 8.10 (d, *J*=5.3 Hz, 1 H) 7.89 (d, *J*=5.3 Hz, 1 H) 7.85 (t, *J*=5.8 Hz, 1 H) 7.52 (dd, *J*=7.6, 1.8 Hz, 1 H) 7.31 - 7.37 (m, 1 H) 7.24 - 7.30 (m, 1 H) 4.82 (d, *J*=5.8 Hz, 2 H) 2.59 (s, 3 H)

^13^C NMR (101 MHz, DMSO-*d*_6_) δ ppm 152.19 (s, 1 C) 145.21 (s, 1 C) 139.88 (s, 1 C) 138.83 (s, 1 C) 131.54 (s, 1 C) 131.10 (s, 1 CH) 129.95 (s, 1 C) 128.62 (s, 1 CH) 127.91 (s, 1 CH) 126.96 (s, 1 CH) 125.15 (s, 1 C) 121.57 (s, 1 CH) 42.67 (s, 1 CH2) 20.02 (s, 1 CH3)

**Synthesis of JNJ-1866**

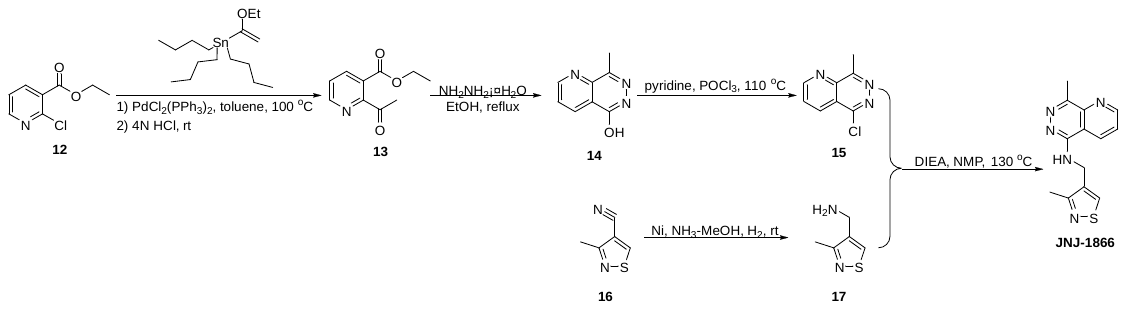

**JNJ-1866: 8-Methyl-*N*-((3-methylisothiazol-4-yl)methyl)pyrido[3,2-*d*]pyridazin-5-amine**

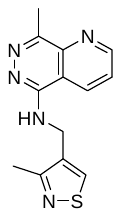

Step 1. Ethyl 2-acetylnicotinate (**13**). Ethyl 2-chloronicotinate (**12**, 20.00 g, 107.8 mmol), tributyl(1-ethoxyvinyl) stannane (77.83 g, 215.5 mmol), a stir bar, and toluene (240 mL) were added to a 1 L three-necked round-bottomed flask, and the reaction vessel evacuated/backfilled with N_2_ three times. Pd(PPh_3_)_2_Cl_2_ (7.56 g, 10.8 mmol) was then added in one portion under N_2_ atmosphere and the reaction mixture was stirred at 100 °C overnight. After cooled to rt with a water bath, 4N HCl (200 mL) was added and the mixture was stirred for further 1 h. The resulting mixture was then extracted with ethyl acetate (3 x 200 mL). The combined organic extracts were washed with brine, dried over anhydrous Na_2_SO_4_, filtered, and concentrated to dryness *in vacuo* to give a brown oil. The brown oil was subjected to silica gel chromatography (eluting with 50% to 70% ethyl acetate/petroleum ether) to afford ethyl 2-acetylnicotinate (**13**, 12.20 g, 59%) as a yellow oil. LCMS (ESI): mass calcd. for C_10_H_11_NO_3_, 193.1; m/z found, 194.2 [M+H]^+^.

Step 2: 8-Methylpyrido[3,2-*d*]pyridazin-5(6*H*)-one (**14**). Ethyl 2-acetylnicotinate (**13**, 9.20 g, 47.6 mmol), hydrazine hydrate (2.62 g, 52.4 mmol), a stir bar, and EtOH (100.0 mL) were added to a 250 mL round-bottomed flask. The resulting mixture was stirred at 50 °C for 2 h. The reaction mixture was then poured into ice-water (300 mL) and the resulting solid was collected by vacuum filtration. The solid was washed with water then air dried (on the filter paper) to afford 8-methylpyrido[3,2-*d*]pyridazin-5(6*H*)-one as an off-white solid (**14**, 5.50 g, 72% yield). LCMS (ESI): mass calcd. for C_8_H_7_N_3_O, 161.1; m/z found, 160.0 [M-H]^-^.

Step 3. 5-Chloro-8-methylpyrido[3,2-*d*]pyridazine (**15**). 8-Methylpyrido [3,2-*d*] pyridazin-5(6*H*)-one (**14**, 200.0 mg, 1.241 mmol), a stir bar, and phosphorus oxychloride (10.0 mL) were added to a 50 mL round-bottomed flask at rt. Pyridine (98.0 mg, 1.24 mmol) was then added dropwise via syringe. The reaction vessel was stirred at 110 °C for 3 h. After cooling, the reaction was poured into 50 mL ice-cold water and extracted with chloroform (3 x 50 mL). The combined organic layer was washed with water (50 mL), then with 1N sodium hydroxide (50 mL), dried over anhydrous Na_2_SO_4_, filtered, and concentrated to dryness *in vacuo* to give a brown oil. The brown oil was subjected to reversed phase column (eluting with 0 to 80% acetonitrile/water) to afford 5-chloro-8-methylpyrido [3,2-*d*] pyridazine (**15**, 116.0 mg, 52%) as a yellow solid. LCMS (ESI): mass calcd. for C_8_H_6_ClN_3_, 179.0/181.0; m/z found, 180.0/182.0 [M+H]^+^.

Step 4: (3-Methylisothiazol-4-yl)methanamine (**17**). 3-Methylisothiazole-4-carbonitrile **(16**, 700.0 mg, 5.638 mmol), ammonia in MeOH (70.0 mL, 7 M), Nickel (168.0 mg), and a stir bar were added to an oven-dried and nitrogen-purged 250 mL three-necked round-bottomed flask, which was subsequently evacuated and refilled with H_2_ (~3 atm) three times and stirred at rt for 16 h. The reaction mixture was then filtered, and the filtrate concentrated to dryness *in vacuo* to give (3-methylisothiazol-4-yl)methanamine as a yellow solid (**17**, 570.0 mg, 79%). LCMS (ESI): mass calcd. for C_5_H_8_N_2_S, 128.0; m/z found, 129.1 [M+H]^+^.

Step 5: 8-Methyl-*N*-((3-methylisothiazol-4-yl)methyl)pyrido[3,2-*d*]pyridazin-5-amine (**JNJ-1866**). (3-Methylisothiazol-4-yl)methanamine (**17**, 200.0 mg, 0.4210 mmol), a stir bar, DIEA (163.8 mg, 1.267 mmol), NMP (2.00 mL), and 5-chloro-8-methylpyrido[2,3-*d*]pyridazine (**15**, 75.7 mg, 0.421 mmol) were added to an 8 mL vial, and the resulting mixture was stirred at 130 °C for 16 h. The reaction vessel was removed from the heating mantle and allowed to gradually cool to rt, and the reaction mixture was then treated with water (50 mL), followed by extracting with ethyl acetate (3 x 50 mL), and the combined extracts were washed with brine, dried over anhydrous Na_2_SO_4_, filtered, and concentrated to dryness *in vacuo* to give a brown oil. The brown oil was subjected to *prep*-HPLC (Column: XBridge Prep OBD C18 Column, 19 x 150 mm, 5 μm; Mobile Phase A: water (10 mmol/L NH_4_HCO_3_ + 0.1% NH_3_), Mobile Phase B: ACN; Flow rate: 60 mL/min; Gradient: 5 to 20% B in 9 min; Wave Length: 254 nm; Rt_1_ (min): 8.38) to give 8-methyl-*N*-((3-methylisothiazol-4-yl)methyl)pyrido[2,3-*d*]pyridazin-5-amine as an off-white solid (**JNJ-1866**, 8.8 mg, 8%).

HRMS (ESI+) *m/z* [M + H]^+^ calculated for C_13_H_14_N_5_S: 272.0892; found: 272.0976. ^1^H NMR (400 MHz, DMSO-*d*_6_) δ ppm 9.15 (dd, *J*=4.4, 1.6 Hz, 1 H) 8.69 - 8.82 (m, 2 H) 7.94 (t, *J*=5.3 Hz, 1 H) 7.89 (dd, *J*=8.3, 4.4 Hz, 1 H) 4.73 (d, *J*=5.1 Hz, 2 H) 2.72 (s, 3 H) 2.48 (s, 3 H). ^13^C NMR (101 MHz, DMSO-*d*_6_) δ ppm 165.51 (s, 1 C) 154.45 (s, 1 CH) 153.20 (s, 1 C) 150.62 (s, 1 C) 146.58 (s, 1 CH) 141.91 (s, 1 C) 135.52 (s, 1 C) 131.12 (s, 1 CH) 125.97 (s, 1 CH) 113.54 (s, 1 C) 38.01 (s, 1 CH2) 17.82 (s, 1 CH3) 17.39 (s, 1 CH3).

The 1^H^, 19^F^, 13^C^ NMR spectra are shown in **Fig. S30**.

Mitochondrial toxicity

To assess the mitochondrial toxicity potential of compounds, the cytotoxicity potential (determined by ATP measurement) of the compounds was compared in glucose versus galactose culture conditions. Briefly, HepG2 cells, either grown in glucose or galactose-containing DMEM media, were seeded at 3000 cells/well in 384-well plates (Corning Inc., New York) and allowed to adhere for 24 h. The HepG2 cell line (human hepatocellular carcinoma cells) was originally obtained from ATCC (HB-8065). The cells were grown, expanded and frozen (mother bank) according to the provider’s instructions. Compounds were prepared as 100× concentrated stock solutions in dimethyl sulfoxide (DMSO), and the final DMSO concentration was 1%. Cells were exposed to the compounds (concentration range: 0.2–100 μM, dilution factor 2) for 24 h before cellular ATP content was measured. Cellular ATP concentrations were assessed by using the CellTiter-Glo 2.0 Assay (Promega, Madison, WI), and the readout was performed by detecting luminescence on an EnVision 2105 Multilabel Reader (PerkinElmer, Waltham, MA). Results were imported in Genedata Screener software (Genedata, Basel, Switzerland) to create dose–response curves and calculate the IC_50_ values of each compound. Subsequently, the ratio between Glu IC_50_ and Gal IC_50_ for each compound was calculated. Compounds with a significantly higher cytotoxicity potential in galactose-grown cells (Glu/Gal ratio ≥ 5) are determined as substances that induce mitochondrial dysfunction as the primary mode of action.

Cytotoxicity

To assess the cytotoxicity potential of compounds, cellular adenosine triphosphate (ATP) content was determined, as an indicator of metabolically active cells. Briefly, HepG2 cells, grown in EMEM media, were seeded at 1000 cells/well in 384-well plates (Corning Inc., New York) and allowed to adhere for 24 h. The HepG2 cell line (human hepatocellular carcinoma cells) was originally obtained from ATCC (HB-8065). The cells were grown, expanded and frozen (mother bank) according to the provider’s instructions. Compounds were prepared as 100× concentrated stock solutions in dimethyl sulfoxide (DMSO), and the final DMSO concentration was 1%. Cells were exposed to the compounds (concentration range: 0.2–100 μM, dilution factor 2) for 72 h before cellular ATP content was measured. Cellular ATP concentrations were assessed by using the CellTiter-Glo 2.0 Assay (Promega, Madison, WI), and the readout was performed by detecting luminescence on an EnVision 2105 Multilabel Reader (PerkinElmer, Waltham, MA). Results were imported in Genedata Screener software (Genedata, Basel, Switzerland) to create dose–response curves and calculate the IC_20_ values of each compound. Test items are strongly cytotoxic with IC_20_ value <10 μM and moderately cytotoxic with IC_20_ values between 10 and 30 μM.

Ames II Mutagenicity Assay

Test compounds and/or their metabolites were evaluated for their mutagenic ability using histidine-requiring *Salmonella typhimurium* strains in the absence or presence of an exogenous mammalian metabolic activation system (S9 homogenate). Histidine-negative bacteria were exposed to 8 concentrations (2-fold serial dilution for 500 µg mL^-1^ final concentration) of the test compound for 90 min at 37°C in 0.5 ml exposure medium with histidine to support two cell divisions (in a 48-deepwell block). Afterwards, 3.5 mL pH indicator medium (lacking histidine) was added to each well after which 40 μL was transferred to each 8 wells of a 384 well plate in twelve-fold and incubated in the dark at 37°C for 48 hours.

Growth of histidine-reversed colonies was assessed colorimetric (based on pH shift) in high throughput using the AmesII^TM^ Mutagenicity Assay (Xenometrix AG, Allschwil, Switzerland) with a total of 96 wells counted per concentration and condition. Test evaluation criteria used to define a mutagenic effect in at least one treatment condition consisted of (1) >3-fold increase in the number of revertant colonies upon exposure at more than one concentration compared to the concurrent solvent control (2) concentration-related increase in the number of revertants (3) number of revertants outside of historical solvent control range and (4) revertant colonies observed in the wells after microscopical evaluation. Positive controls must show at least a 3-fold increase in the number of revertant wells above solvent control value and above the historical solvent control range.

DMSO was used as solvent control, 2-aminoanthracene (10 µg mL^-1^) as positive control in the presence of metabolic activation and 2-nitrofluorene (10 µg mL^-1^) and 4-nitroquinoline-N-oxide (1 µg mL^-1^) as positive controls in the absence of metabolic activation. The mammalian liver post-mitochondrial fraction (S9 homogenate) was used for metabolic activation and was prepared from male Sprague Dawley rats induced with 5.6-benzoflavone/phenobarbital and purchased from Molecular Toxicology Inc. (Boone, USA). The cofactors used consisted of a NADPH regenerating system which was purchased from Molecular Toxicology Inc.

hERG

The effect of the test compounds on the hERG current was studied in a human embryonic kidney cell line (HEK293) with stable transfection of hERG (human *ether-à-go-go*-related gene; purchased from Dr. Z. Zhou, University of Wisconsin, Madison, USA) using an automated planar patch clamp system. Final dilutions of the test compounds were prepared with recording solution using automated liquid handling (final DMSO concentration: 0.3 %). Nominal concentrations of 1 μM, 3 μM, 10 μM and 30 μM (rounded values) were tested. Patch-clamp experiments were performed in the voltage-clamp mode and whole-cell currents were recorded with an automated patch-clamp assay utilising the SyncroPatch® 384PE system (Nanion Technologies, Munich, Germany). After establishing whole-cell configuration, test pulses were given to stabilise the ion channel currents in control conditions. While continuing the pulse protocols, either vehicle control, test article, or positive control was added and effects were measured after at least 4-5 min of drug application. The leak-corrected hERG current (K+-selective outward current) was determined as the maximal tail current at -30 mV after a 2-sec depolarization to +70 mV. The median current from three sequential voltage pulses was taken at the end of the control period and after adding test article and vehicle to calculate the percent inhibition. Concentration/response relation (IC_50_) was calculated by non-linear least-squares fits using individual values.

Ca^2+^ transient measurement assay on human induced pluripotent stem cell-derived cardiomyocytes (CTCM)

Human iPSC-derived cardiomyocytes (iCell-Cardiomyocytes2) (donor: 01434, Female, age <18,) were purchased from CDI (Cellular Dynamics Inc. - Fujifilm, USA), as frozen vials. Cells were seeded in fibronectin-coated 96-well plates at a density suited to form a monolayer (i.e. 50,000 cells/well) and maintained in culture in a stage incubator (37^o^C, 5% CO_2_), according to the instructions of the cell provider. The experiments with test compounds were carried out 4 to 6 days after plating, when the cells formed a living, beating monolayer. On the day of the experiment, the hiPSC-CMs were incubated with a physiological saline solution (Tyrode, Sigma), supplemented with KCl, HEPES and glucose. After 1h, cells were loaded with the Ca2+-dye ‘Cal-520’ (Cat 21130; AAT Bioquest) (50 µg, MW: 1103/mol). Spontaneous electrical activity was recorded using the Functional Drug Screen System (FDSS/µCell; Hamamatsu, Japan) and the recordings were subsequently analysed using the internal data management and analysis software SPEc II (BPT House Belgium). The calcium-transient properties were evaluated at baseline, and at 5, 15 and 30 min after compounds addition. Several parameters were used to score the compounds from ‘No’ to ‘Very High’ hazard score, including the duration of the calcium transient at 90% of the decay (CTD90), the amplitude of the transient (Amp), and beating rate (BR), at 30 min after compounds addition. Additionally, the presence of arrhythmia-like events (such as early-after depolarization) and quiescence (i.e., a complete cessation of beating of the preparation), were also accounted in the score generation. Further details on how the pro-arrhythmic hazard score is generated have been published previously (*67*).

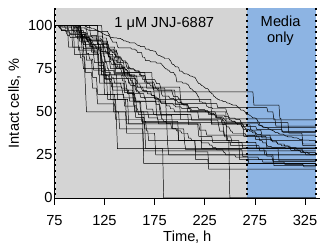

Fig. S1: Graph depicting time-lapse microscopy data – Fraction of intact cells upon exposure to 1 µM JNJ-6887 treatment (grey shading) and after washout (blue), from the single-cell imaging experiments. The lines represent independent xy positions imaged. n = 29 positions, containing ~1,290 cells. Data shown are representative of two independent experiments.

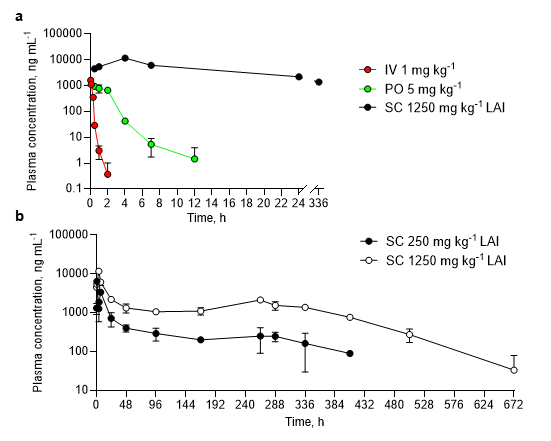

Fig. S2: Pharmacokinetic (PK) studies with JNJ-1866 – **a**, PK profiles of JNJ-1866 administered as IV (1 mg kg^-1^), PO (5 mg kg-1) or SC (1,250 mg kg^-1^; long-acting injectable [LAI]). n = 3 mice (IV and PO) or n = 3 mice (from a total of 6) for LAI PK studies, alternating sampling in 3 mice per time point. **b**, PK profiles following 250 mg kg^-1^ or 1,250 mg kg^-1^ JNJ-1866 over 28 days. n = 3 mice (from a total of 6) alternating sampling in 3 mice per time point. Data shown are mean ± s.d.

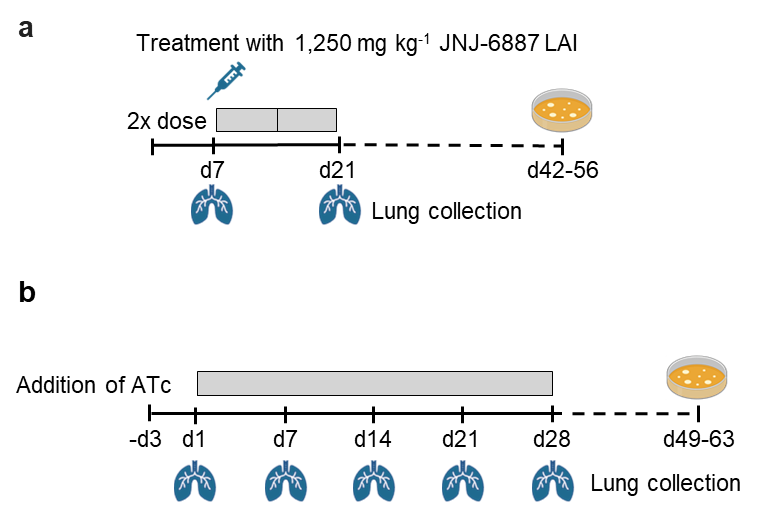

Fig. S3: Schematic of animal studies – Models for short acute (**a**) and CRISPRi-mediated knockdown-based (**b**) animal models of infection.

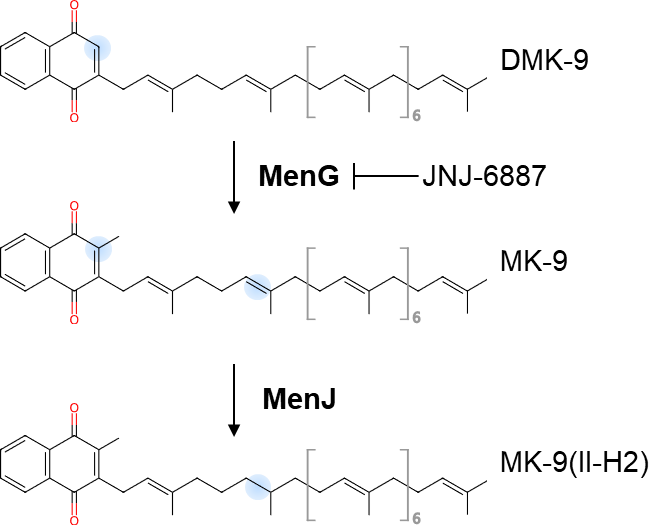

Fig. S4: Schematic of enzyme activity – DMK-9: demethylmenaquinone; MK-9: menaquinone-9; MK-9(II-H_2_): β-dihydromenaquinone-9.

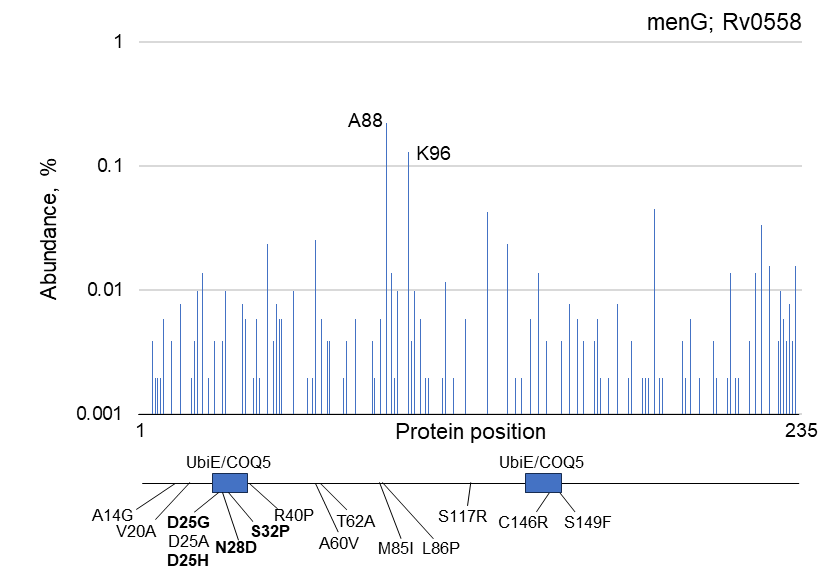

Fig. S5: Genetic diversity of MenG in clinical isolates of *M. tuberculosis* and genetic map of resistance-conferring mutations - Genetic variance of MenG in 51,183 clinical TB isolates (*29*). Protein domains predicted using InterPro domain search (predicted COQ5 domains: 24-39 aa and 135-149 aa). Mutations identified from resistance generation experiments with other MenG-targeting inhibitors shown using other series compounds or reported in the literature (*68*), with resistance-conferring mutations specifically selected against JNJ-6887 and JNJ-1866 shown in bold.

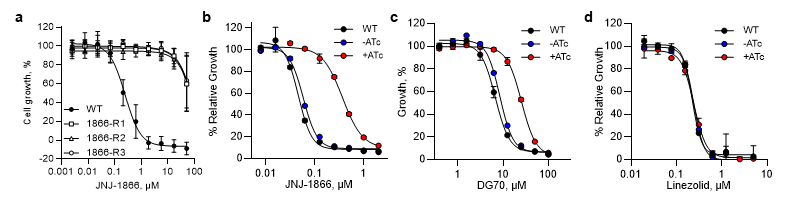

Fig. S6: Confirmation of resistance in MenG resistance and validation strains – **a**, Representative *M. tuberculosis* dose-response curves of JNJ-1866 resistant mutants compared with a drug-sensitive WT strain. MIC_90_ values shown in **Table S6**. n = 4 biological replicates. **b-d**, Potency determination for JNJ-1866 (**b**; MIC_50_: WT = 0.051 µM, -ATc = 0.063 µM, +ATc = 0.416 µM), DG70 (*38*) (**c**; MIC_50_: WT = 7.26 µM, -ATc = 9.21 µM, +ATc = 26.0 µM) and linezolid (**d**; MIC_50_: WT = 0.230 µM, -ATc = 0.240 µM, +ATc = 0.240 µM; negative control) against WT *M. tuberculosis* (black), *M. tuberculosis* maintaining a chromosomally integrated MenG-C146R expression plasmid without (blue) and with (red) 7-day ATc induction. n = 2 biological replicates. Data shown are mean ± s.d.

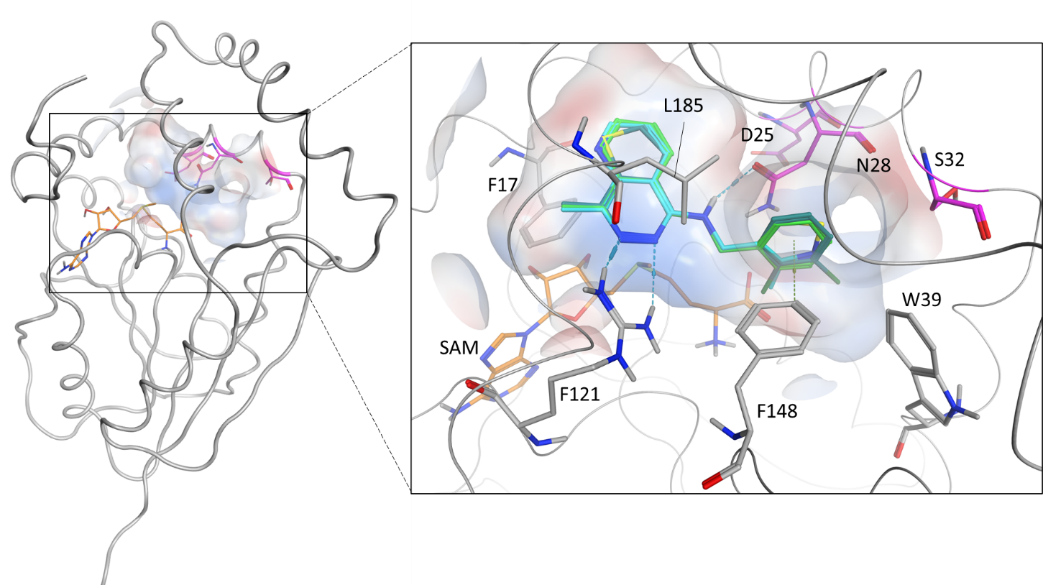

Fig. S7: Docking of JNJ-6887 into an AlphaFold model of *M. tuberculosis* MenG – AlphaFold homology model of MenG (234 amino acids; gray) with SAM (orange) and location of the resistance mutation D25, N28, and S32 (purple). The surface is showing the putative binding pocket and superposed docked ligands are shown in blue, green and teal.

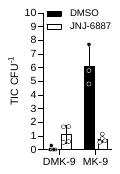

Fig. S8: Independent repeat of metabolomics – Inhibition of menaquinone biosynthesis in *M. tuberculosis* in the presence and absence of 660 nM JNJ-6887 after 6 d of treatment. n = 4 biological replicates. DMK-9: demethylmenaquinone; MK-9: menaquinone-9. Data shown are mean ± s.d. TIC: total ion count.

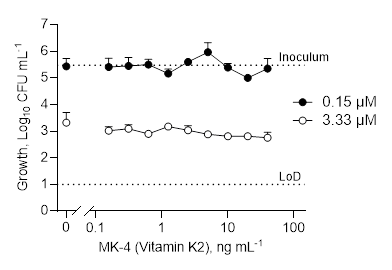

Fig. S9: MK-4 (vitamin K2) does not rescue JNJ-6887 activity - Concentrations of 0.15 µM and 3.33 µM JNJ-6887 in the presence of varying concentrations of MK-4 (vitamin K2). LoD: Limit of detection. Human serum levels of MK-9 have been measured with a range of 0.050-1.598 ng mL^-1^ (*35*). n = 3 technical replicates. Data shown are mean ± s.d.

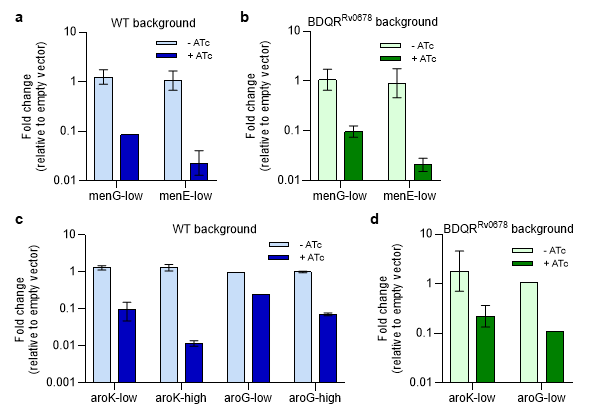

Fig. S10: Validation of CRISPRi-mediated transcript knockdown strains - qRT-PCR of CRISPRi *aroG, aroK, menE*, *menG* strains, transcript levels were normalised against a reference gene, *sigA*. Strains were induced with 100 ng mL^-1^ ATc and incubated for 2-4 days. **a**, *menG* and *menE* strains in WT background, 2-day induction. **b**, *menG* and *menE* strains in BDQR^Rv0678^ background, 2-day induction. **c**, *aroK*-low/high and *aroG*-low/high strain in WT background, 4-day induction. **d**, *aroK*-low and *aroG*-low strain in BDQR^Rv0678^ background, 4-day induction. n = 2 biological replicates (menG, menE, aroK) or n = 1 biological replicate (aroG). Data is the mean ± s.d.

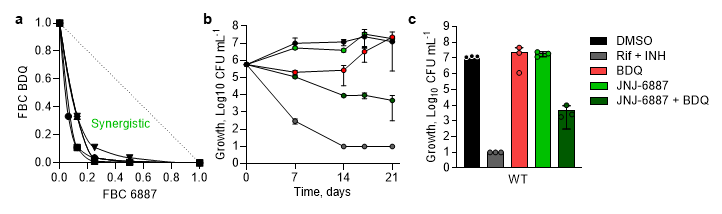

Fig. S11: Determination of optimal synergist concentrations of BDQ and JNJ-6887 – **a**, Isobologram showing synergy between JNJ-6887 and bedaquiline (BDQ). Five independent experiments are shown. FBC: Fractional bactericidal concentration. **b-c,** Time-kill kinetic assay (**b**) using 0.13 µM bedaquiline (1x MIC_90_ BDQ) and 0.11 µM JNJ-6887 (2x MIC_90_) in a WT background. These optimum synergistic concentrations were determined based on the isobologram. Dimethyl sulfoxide (DMSO; black) used as a negative kill control. Rifampicin (RIF; 14.58 µM) and isoniazid (INH; 5.8 µM) (grey) used as positive kill control. Day 21 CFU counts (**c**) from time-kill kinetics. n = 3 biological replicates. Two independent experiments each consisting of three technical replicates. Representative experiment shown. Data shown are mean ± s.d.

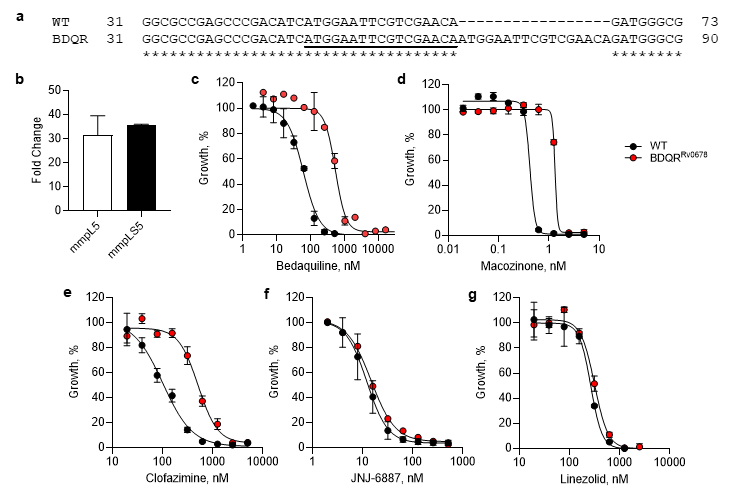

Fig. S12: Validation of BDQ-resistant (BDQR^Rv0678^) strain – **a**, Visualisation of indel in *Rv0678* and impact on translation via a 17 bp insertion (duplication of the region between A49 to A65 inserted between nucleotides A65 and G66). **b**, qRT-PCR showing upregulation of *mmpL5*-*mmpS5* efflux pump components. Transcript levels were normalised against a reference gene, sigA, then against WT relative transcript levels. n = 2 independent experiments consisting of two technical replicates. **c-g**, Dose-response curves of bedaquiline (**c**; MIC_50_: WT = 61.1 nM, BDQR^Rv0678^ = 549 nM), macozinone (**d**; MIC_50_: WT = 0.43 nM, BDQR^Rv0678^ = 1.33 nM), clofazimine (**e**; MIC_50_: WT = 105 nM, BDQR^Rv0678^ = 511 nM), JNJ-6887 (**f**; MIC_50_: WT = 13.0 nM, BDQR^Rv0678^ = 16.3 nM) and linezolid (**g**; MIC_50_: WT = 265 nM, BDQR^Rv0678^ = 320 nM; control) against WT (black) and BDQR^Rv0678^ (red) strains. n = 3 biological replicates. Data shown are mean ± s.d.

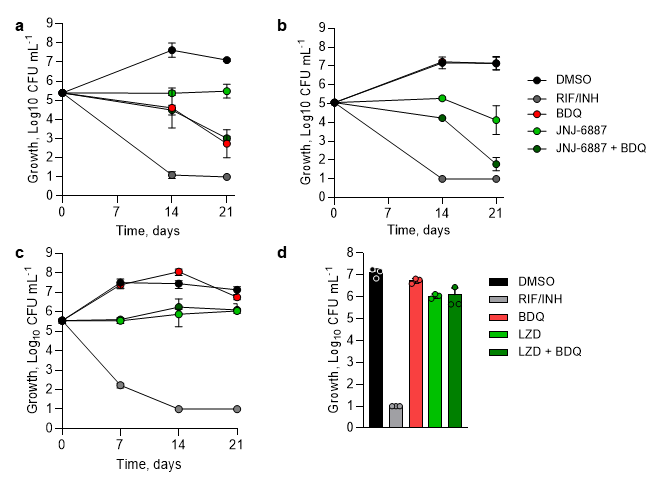

Fig. S13: Resensitisation of bedaquiline resistance – **a-b,** Time-kill kinetics assays using monotherapies and combination of 0.5 µM bedaquiline (BDQ) and 200 nM JNJ-6887 in (**a**) WT or (**b**) BDQR^Rv0678^ background strain related to data shown in **Fig. 3a**. Concentrations were determined as the highest inhibitor concentration that that had no bactericidal activity in the BDQR^Rv0678^ strain. Two independent experiments each consisting of three technical replicates. n = 3 biological replicates. Representative experiment shown. **c-d**, Time-kill kinetics assay (**c**) using monotherapies and combination of 0.5 µM bedaquiline (BDQ) and 2.5 µM linezolid (LZD) in the BDQR^Rv0678^ background, showing no resensitisation of bedaquiline resistance. Day-21 CFU counts (**d**). Dimethyl sulfoxide (DMSO; black) used as a negative kill control. Rifampicin (RIF; 14.58 µM) and isoniazid (INH; 5.8 µM) (grey) used as positive kill control. Data shown are mean ± s.d.

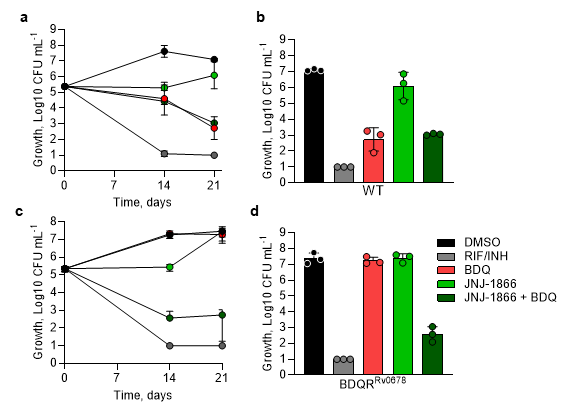

Fig. S14: Resensitisation with JNJ-1866 – **a-b**, Time-kill kinetics assay (**a**) using monotherapies and combination of 0.5 µM bedaquiline (BDQ) and 0.9 µM JNJ-1866 in a WT background. Concentrations were determined as the highest inhibitor concentration that that had no bactericidal activity in the BDQR^Rv0678^ strain. Day 21 CFU counts from time-kill kinetics assay (**b**). **c-d**, Time-kill kinetic assay (**c**) using monotherapy and combination of BDQ and JNJ-1866 in a BDQR^Rv0678^ background strain. Day 21 CFU counts from time-kill kinetics assay (**d**). DMSO (black) used as a negative kill control. Rifampicin (RIF; 14.58 µM) and isoniazid (INH;5.8 µM) used together as positive kill control (dark grey). n = 3 biological replicates. Two independent experiments each consisting of three technical replicates ± SD. Representative experiment shown. Data shown are mean ± s.d.

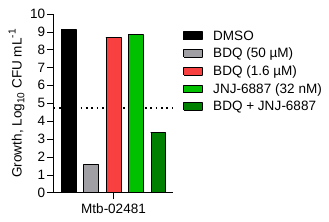

Fig. S15: Resensitisation in an alternative efflux-based resistant strain – Day 21 CFU counts from a time-kill kinetic study with an alternative *Rv0678* mutant (A344del; 2013-02481) purchased from BCCM/ITM. Combination of 1.6 µM bedaquiline (BDQ) and 32 nM JNJ-6887 resensitised the strain to bedaquiline. Concentrations were determined as the highest inhibitor concentration that that had no bactericidal activity in this strain.

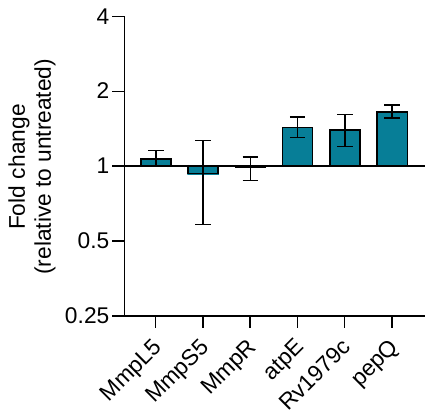

Fig. S16: Transcript levels of genes related to bedaquiline resistance in BDQR^Rv0678^ background following JNJ-6887 treatment - Transcript levels were normalised against a reference gene, sigA, then against WT relative transcript levels. n = 2 independent experiments consisting of two technical replicates. Data shown are mean ± s.d.

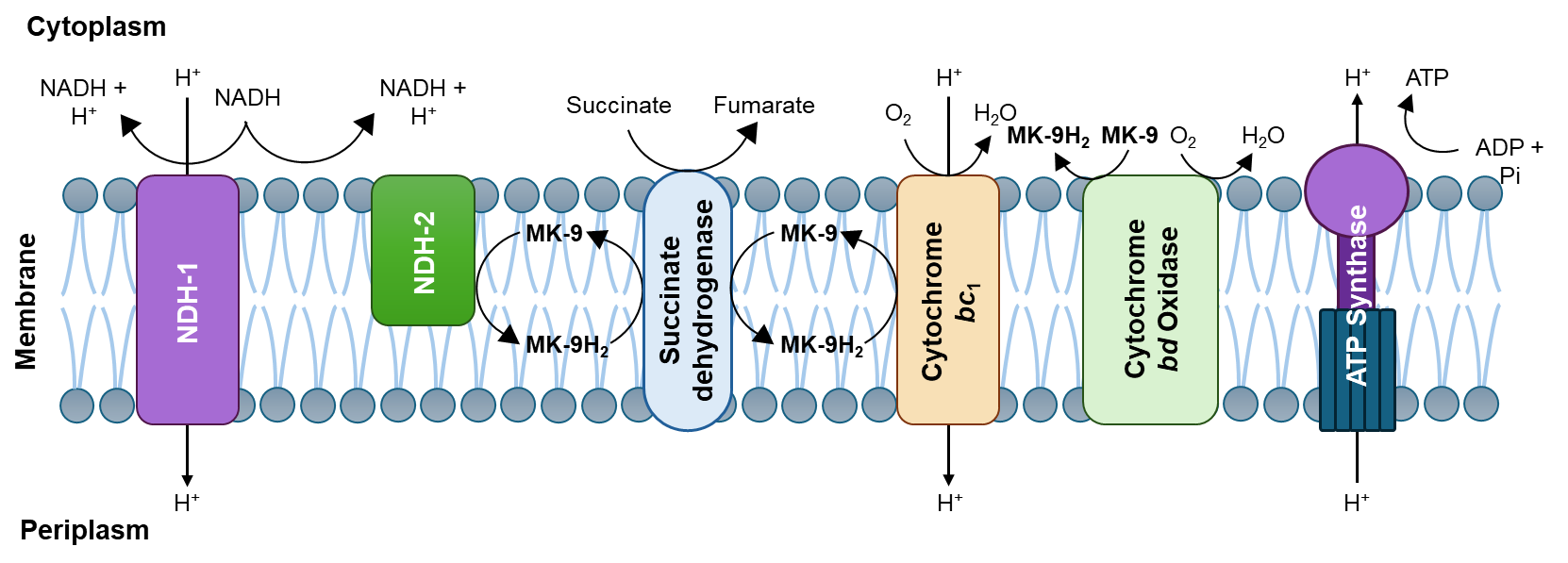

Fig. S17: Outline of the electron transport chain (ETC) **-** The *M. tuberculosis* ETC, also known as the Electron Transfer System (ETS), relies on menaquinone-9 (MK-9) as well as β-dihydromenaquinone-9 (MK-9(II-H2)) (*34*) as central lipid-soluble electron carriers. These shuttle electrons from primary dehydrogenases to terminal oxidases, sustaining respiration and ATP synthesis. Electrons enter the ETC through NDH-1 (Complex I), a multi-subunit, proton-pumping NADH dehydrogenase, or through NDH-2, a single-subunit, non-proton-pumping flavoprotein that transfers electrons directly to MK-9. Additional dehydrogenases feed into the menaquinone pool, which is reduced to MK-9H₂ and subsequently donates electrons to the cytochrome *bc*₁ complex (Complex III) or cytochrome *bd* oxidase (Complex IV), generating the proton motive force (PMF) that drives ATP synthase (Complex V; target of bedaquiline), making both components essential for oxidative phosphorylation.

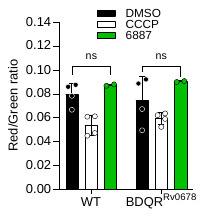

Fig. S18: JNJ-6887 does not impair membrane potential – Bacteria treated with 1 µM JNJ-6887 for 1 hour compared to positive control (25 µM CCCP: carbonyl cyanide m-chlorophenyl hydrazone), in WT and BDQR^Rv0678^ backgrounds. n = 3 biological replicates. Representative from two independent experiments. Data shown are mean ± s.d.

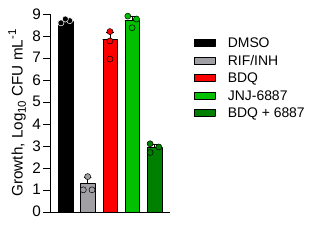

Fig. S19: Resensitisation of a JNJ-6887-resistant mutant with bedaquiline – Day 21 CFU counts from a time-kill kinetics assay using monotherapies and combination of 0.4 µM bedaquiline (BDQ) and 8.9 µM JNJ-6887 in a JNJ-6887-resistant (D25G) background. Concentrations were determined as the highest inhibitor concentration that that had no bactericidal activity in this strain. Rifampicin (RIF; 14.58 µM) and isoniazid (INH; 5.8 µM) used together as positive kill control (dark grey). n = 3 biological replicates. Data shown are mean ± s.d.

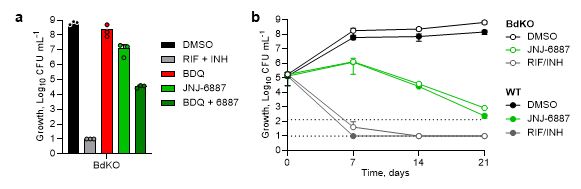

Fig. S20: Time-kill kinetics with bedaquiline and JNJ-6887 in a cytochrome *bd* knockout – **a**, Day 21 CFU counts from a time kill-kinetics assay using monotherapies and combination of 0.1 µM BDQ and 0.2 µM JNJ-6887 in a WT strain with clean deletion of cytochrome *bd* oxidase (complex IV). Concentrations were determined as the highest inhibitor concentration that that had no bactericidal activity in this strain. n = 3 biological replicates. Two independent experiments each consisting of three technical replicates. Representative experiment shown. **b**, Time-kill kinetics comparing the effect of 3.3 µM JNJ-6887 in the presence and absence of cytochrome *bd* oxidase deletion. WT (solid circles) and *bd*KO (open circles) plotted, DMSO was used as a negative kill control, 3.3 µM JNJ-6887, and combination of rifampicin (14.58 µM) and isoniazid (5.8 µM) was used a positive kill control. n = 3 biological replicates. Data shown are mean ± s.d.

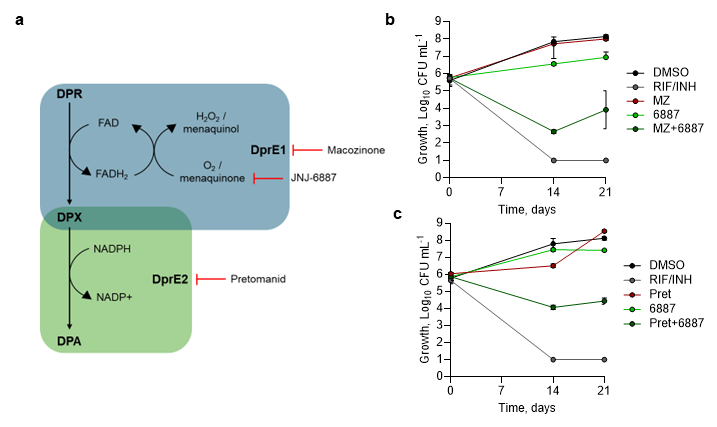

Fig. S21: Time-kill kinetics with macozinone, pretomanid and JNJ-6887 – **a**, The DPA pathway and its interactions with macozinone, pretomanid and JNJ-6887. Decaprenylphosphoryl-D-ribose (DPR) is oxidised by DprE1, using flavin adenine dinucleotide (FAD) as a cofactor, to form decaprenylphosphoryl-2-keto-erythro-pentofuranose (DPX). Menaquinone-9 (MK-9) then accept electrons from reduced FAD (FADH_2_) and transfers them to the electron transport chain, allowing FAD to be recycled for DprE1 to continue functioning. DPX is subsequently reduced to decaprenylphosphoryl-D-arabinose (DPA) by DprE2 converting nicotinamide adenine dinucleotide phosphate (NADPH) to NADP+. DPA is the sole arabinose donor for cell wall arabinan synthesis and is essential for *M. tuberculosis* growth. JNJ-6887 inhibits MK-9 production, which is required for DprE1 and indirectly, DprE2 function, providing a mechanistic explanation for the enhanced activity observed in combination with macozinone (DprE1 inhibitor) and pretomanid (DrpE2 inhibitor). **b-c**, Time-kill kinetics assays using monotherapy and combination of (**b**) 1.5 nM macozinone and (**c**) 500 nM pretomanid with 0.2 µM JNJ-6887. Concentrations were determined as the highest inhibitor concentration that that had no bactericidal activity in the BDQR^Rv0678^ strain. DMSO (black) used as a negative kill control. Related to **Fig. 4b**. Representative of two independent experiments shown. n = 3 biological replicates. Rifampicin (RIF; 14.58 µM) and isoniazid (INH; 5.8 µM) used together as positive kill control (dark grey). Data shown are mean ± s.d.

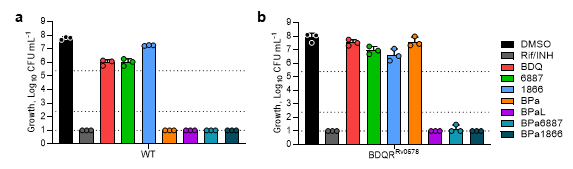

Fig. S22: Combination time-kill kinetics in WT and BDQR^Rv0678^ – **a-b**, Day 21 CFU counts from a time-kill kinetics of 0.5 µM bedaquiline (B), 7 µM pretomanid (Pa), 6 µM linezolid (L), 0.2 µM JNJ-6887 (6887) and 0.9 µM JNJ-1866 (1866) in (**a**) WT and (**b**) BDQR^Rv0678^ background. Rifampicin (RIF; 14.58 µM) and isoniazid (INH; 5.8 µM) (grey) used as positive kill control. n = 3 biological replicates. Representative example from two independent experiments. Data shown are mean ± s.d.

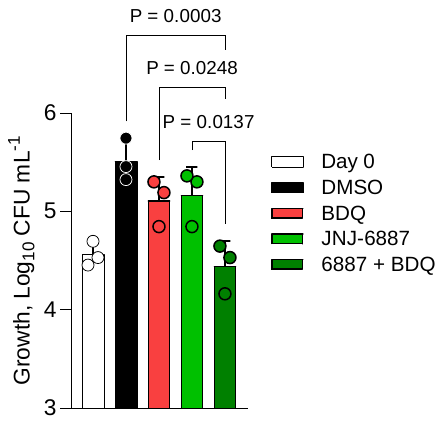

Fig. S23: Resensitisation in an **intracellular THP-1** macrophage model - Macrophage survival assay using activated THP-1 cells infected with BDQR^Rv0678^ background strain. Cells were treated with monotherapy of 0.12 µM bedaquiline (BDQ) or 10 µM JNJ-6887 and a combination. Cells were lysed and CFU counted at Day 0 and 3. One independent experiment consisting of three biological replicates. Data shown is the mean ± s.d.

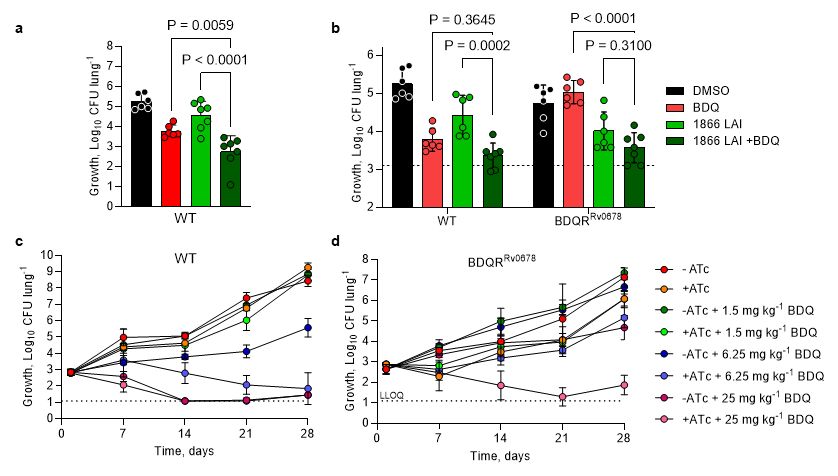

Fig. S24: *In vivo* proof-of-concept of resensitisation **with** JNJ-1866 **and CRISPRi *menG* knockdown - a,** *In vivo* demonstration of enhancement with 1,250 mg kg^-1^ JNJ-1866 LAI (subcutaneous; once weekly) and 6.25 mg kg^-1^ bedaquiline oral administration (PO) once daily (qd) in the WT background in a short acute model. n = 7 mice. Representative of two independent experiments. **b,** *In vivo* efficacy of 1,250 mg kg^-1^ JNJ-1866 (subcutaneous, once fortnightly) and 6.25 mg kg^-1^ bedaquiline (PO; qd) combination in WT (left) and BDQR^Rv0678^ (right) backgrounds. Line indicates CFU at treatment start (WT: 2.9 Log_10_; BDR^Rv0678^: 3.25 Log_10_). n = 6 mice **c**, *In vivo* demonstration of bedaquiline enhancement using a MenG “low” CRISPRi in a WT background using a short acute model. CRISPRi strain was pre-induced for 7 days. n = 5 mice. **d**, *In vivo* demonstration of bedaquiline resensitisation using a MenG “low” CRISPRi in a BDQR^Rv0678^ background. n = 5 mice. BDQ: bedaquiline. Full time course experiments related to **Fig. 4d**.

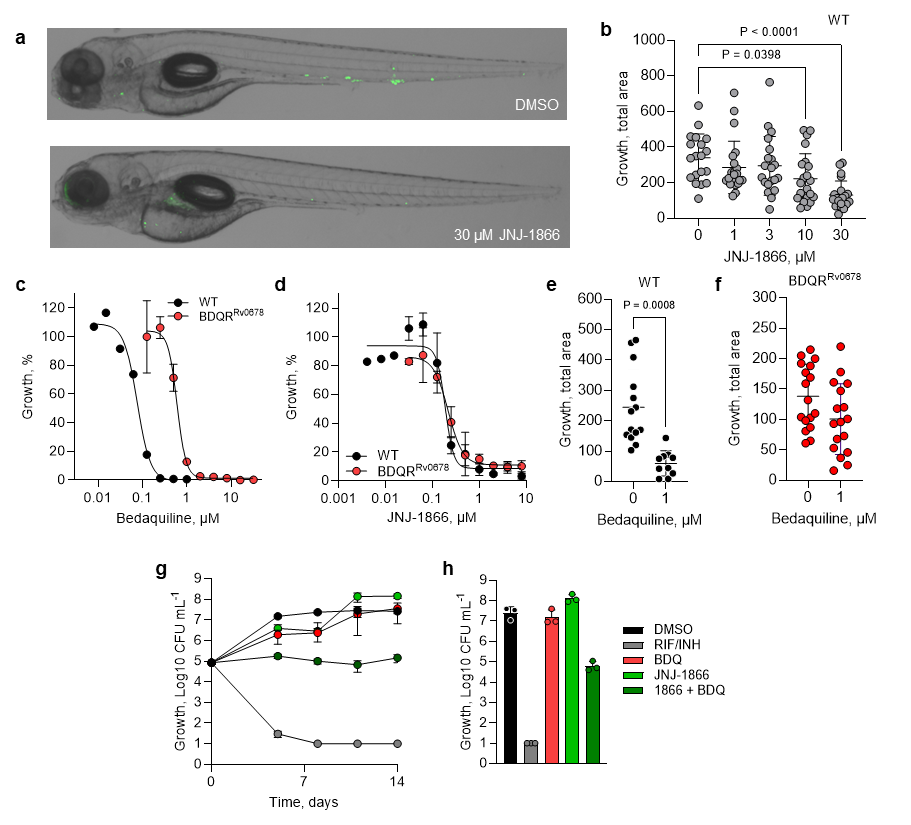

Fig. S25: Resensitisation of bedaquiline-resistant *Mycobacterium marinum-infected* zebrafish with JNJ-1866 **– a**, Representative images of *M. marinum*-infected (400 CFU) zebrafish treated for 2 days with DMSO or 30 µM JNJ-1866. **b**, *In vivo* proof-of-concept for JNJ-1866 in *M. marinum*-infected zebrafish after 2 days treatment. Zebrafish embryos were infected with 400 CFU *M. marinum* at 2 days post fertilisation (dpf) via the caudal vein and treatment was initiated at 3 dpf. n ≥ 19 zebrafish. **c-d**, *in vitro* dose-response of bedaquiline (**c**; MIC_50_: WT = 80 nM; BDQR^Rv0678^ = 620 nM) and JNJ-1866 (**d**; MIC_50_: WT = 188 nM; BDQR^Rv0678^ = 210 nM) in WT and BDQR^Rv0678^ backgrounds. n = 3 biological replicates. **e-f**, *in vivo* demonstration of bedaquiline resistance between WT (**e**) and BDQR^Rv0678^ (**f**) *M. marinum*-infected zebrafish (400 CFU) after 2 days treatment with 1 µM bedaquiline. Zebrafish embryos were infected at 2 dpf via the caudal vein and treatment was initiated at 3 dpf. n ≥ 10 zebrafish. **g**, Time-kill kinetics of 1.4 µM JNJ-1866 in combination with 570 nM bedaquiline in the BDQR^Rv0678^ background *in vitro*. Concentrations were determined as the highest inhibitor concentration that that had no bactericidal activity in the BDQR^Rv0678^ strain. **h**, Day-11 CFU counts showing resensitisation of bedaquiline resistance. Combination of rifampicin (7 µM) and isoniazid (15 µM) was used a positive kill control. n = 3 biological replicates. Data shown in **b**-**h** are mean ± s.d.

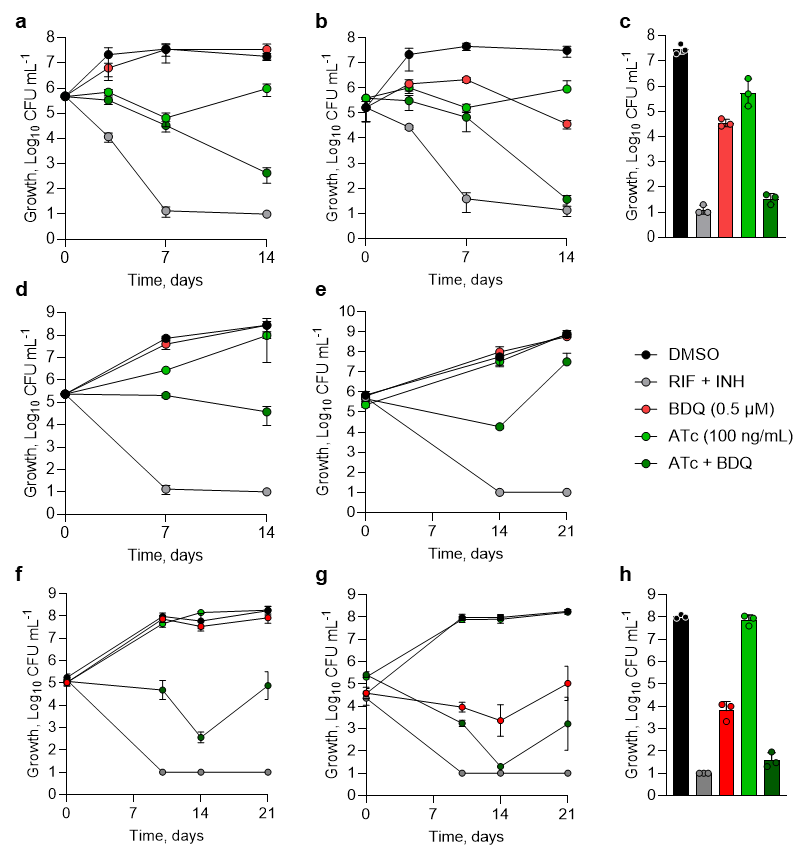

Fig. S26: Time-kill kinetic assays using *menG*, *menE*, *aroG* and *aroK* CRISPRi knockdown strains – **a-c**, strains containing a *menG* “low” inducible transcript knockdown vector, induced with 100 ng mL^-1^ ATc, in combination with 0.5 µM BDQ in either a BDQR^Rv0678^ (**a**) and WT (**b-c**) background strain. Alternative view of day-14 CFU counts shown (**c**). **d-e**, *menE* (**d**) and *aroG* (**e**) “low” knockdown strains in BDQR^Rv0678^ background. **f-h**, strains containing a *aroK* “low” inducible transcript knockdown vector, induced with 100 ng mL^-1^ ATc, in combination with 0.5 µM BDQ in either a BDQR^Rv0678^ (**f**) and WT (**g-h**) background strain. Alternative view of day 14 CFU counts shown (**h**). CRISPRi knockdown was induced with 100 ng mL^-1^ ATc either 7 days (*menG*, *menE*) or 5 days (*aroG*, *aroK*) prior to commencing the experiment. Bedaquiline concentration was determined as the highest concentration that that had no bactericidal activity in the BDQR^Rv0678^ strain. Dimethyl sulfoxide (DMSO; black) used as a negative kill control. Rifampicin (RIF; 14.58 µM) and isoniazid (INH; 5.8 µM) (grey) used as positive kill control. Related to **Fig. 4b-e**. n = 3 biological replicates. Representative of at least two independent experiments. Data shown are mean ± s.d.

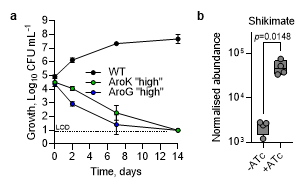

Fig. S27: Validation of essentiality and function of shikimate enzymes – **a**, *In vitro* growth curves of *aroG* and *arok* “high” knockdown strains with 5 µg mL^-1^ ATc. n = 3 technical replicates. **b**, LC-MS analysis showing shikimate accumulation in the *aroK* “high” CRISPRi strain after induction with ATc for 5 days. n = 4 biological replicates. Representative of two independent experiments. Significance was calculated with two-sided (Bonferroni–Dunn) Student’s t-test with Welch correction.

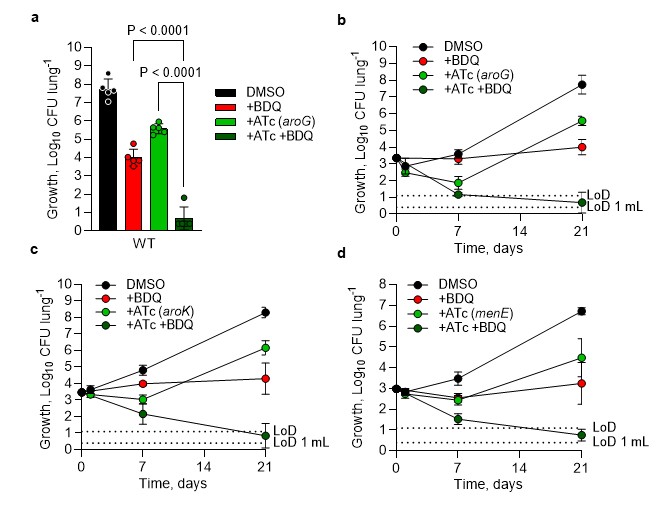

Fig. S28: *In vivo* proof-of-concept of bedaquiline enhancement using CRISPRi knockdown **– a-d**, *In vivo* demonstration of bedaquiline enhancement using *aroG* “high” (**a-b**), *aroK* “high” (**c**) or *menE* “low” (**d**) CRISPRi-mediated knockdown in a WT background using a short acute model. CRISPRi knockdown strains were induced with doxycycline in chow, and in combination with 6.25 mg kg^-1^ bedaquiline (BDQ; once daily oral administration, 5/7 days). Day-21 CFU counts showing enhancement of bedaquiline resistance with *aroG* CRISPRi knockdown (**b**). *menE* CRISPRi strain was pre-induced with 100 ng mL^-1^ ATc for 7 days. n = 5 mice. Data shown are mean ± s.d. Full time course experiments (**c-d**) related to **Fig. 5h-i**.

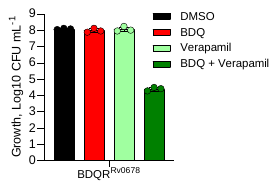

Fig. S29: Rescue of bedaquiline resistance by verapamil **–** Day 14 CFU counts from a time kill kinetics assay with 50 µM verapamil and 0.5 µM BDQ in the BDQR^Rv0678^ background. n = 3 biological replicates. Data shown are mean ± s.d.

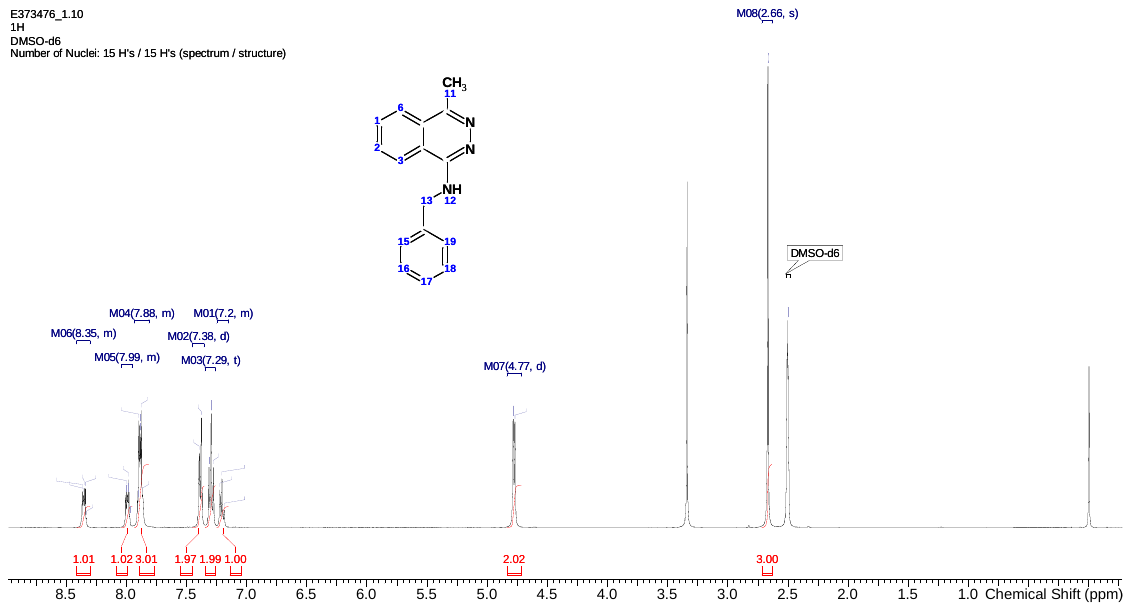

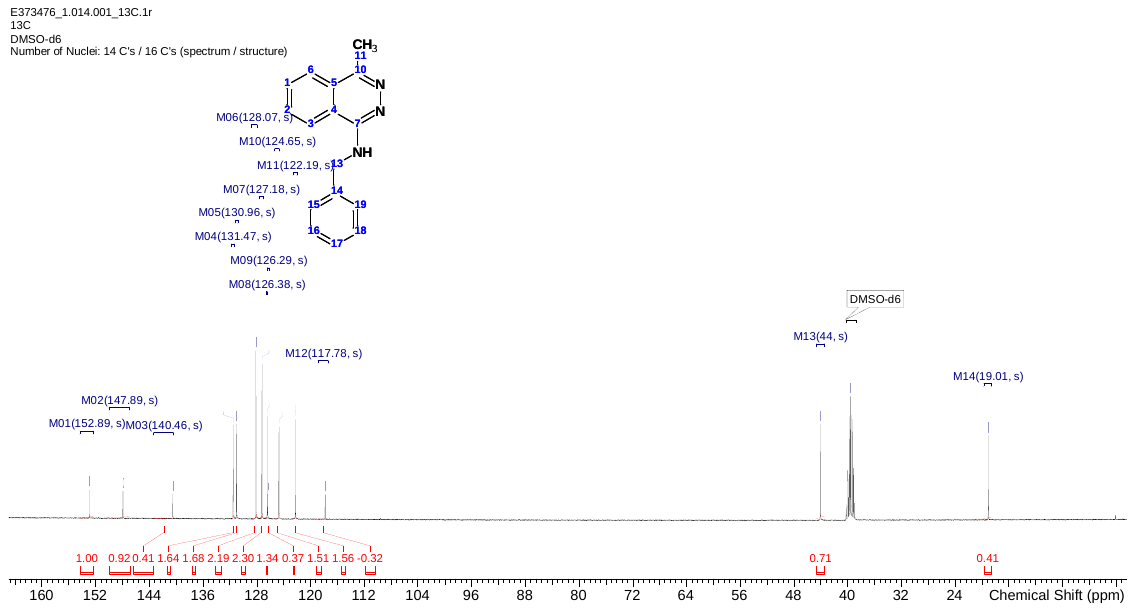

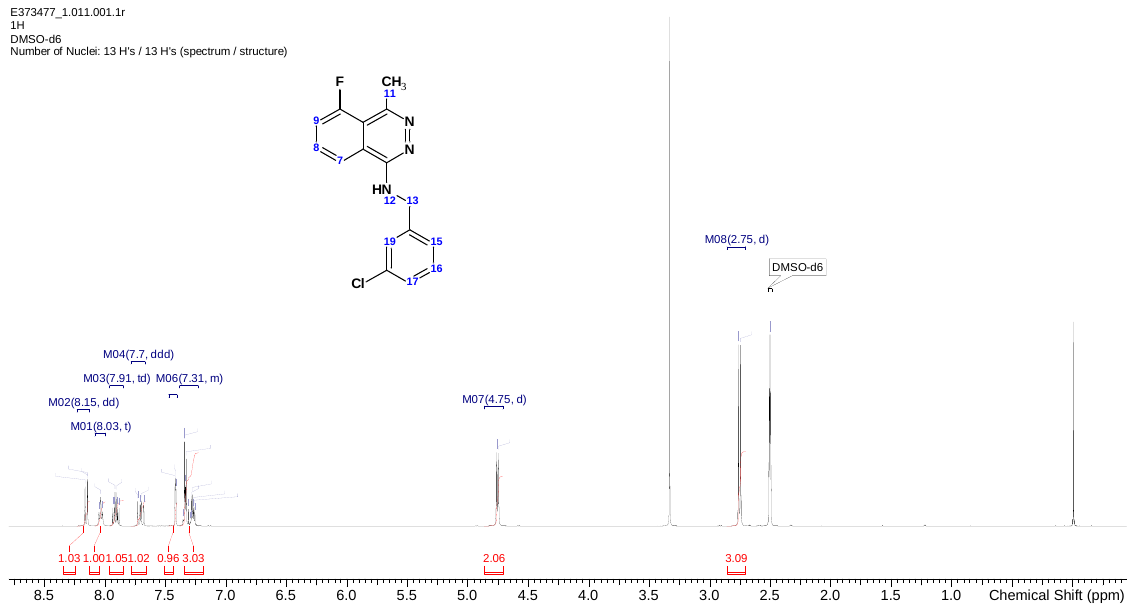

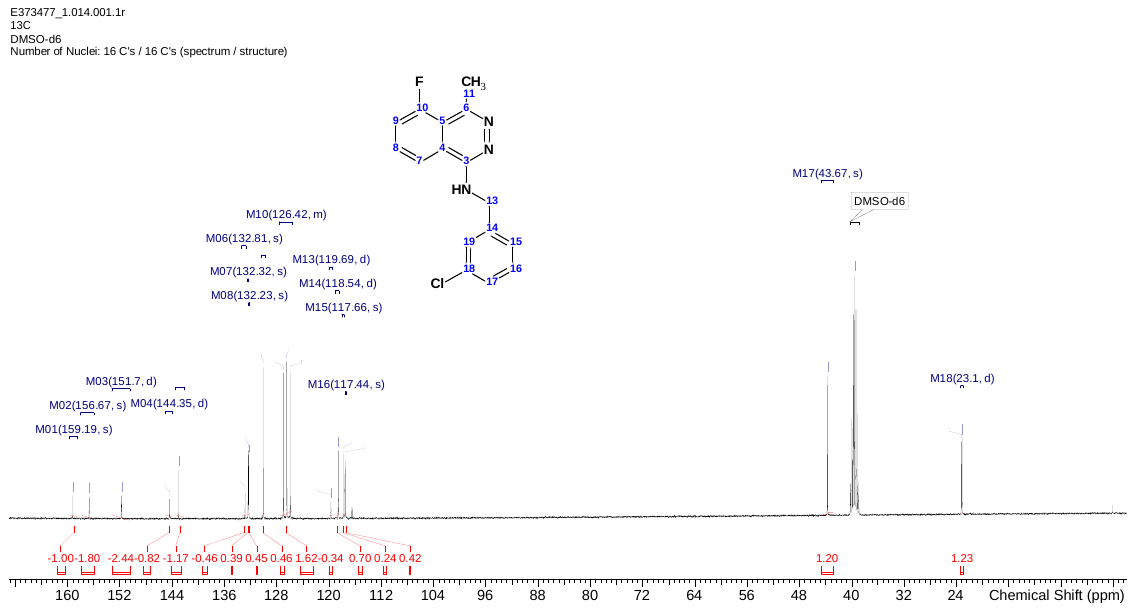

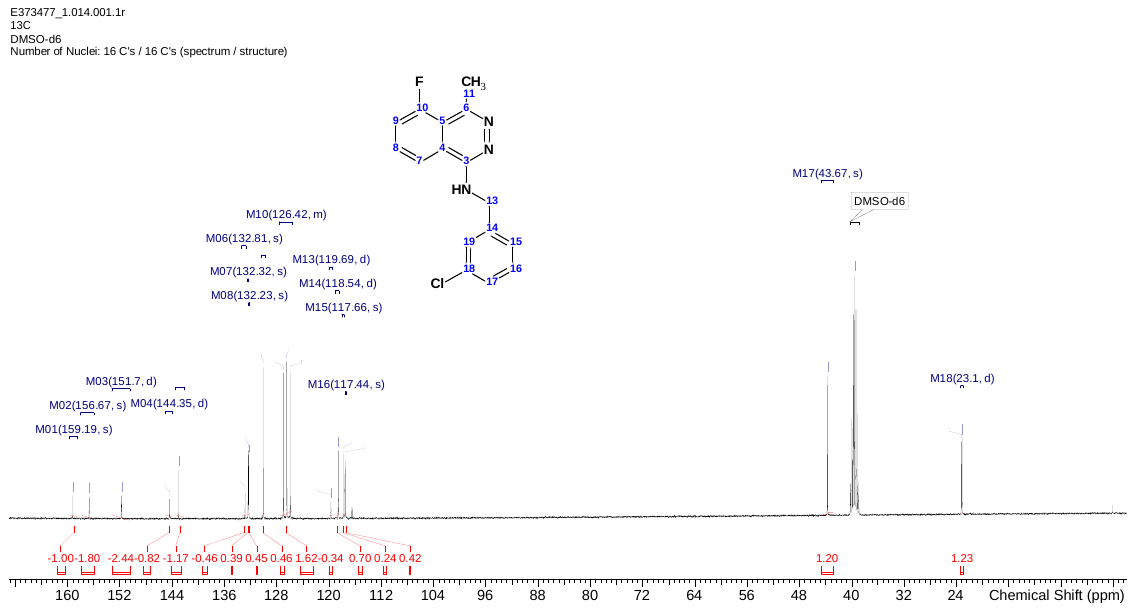

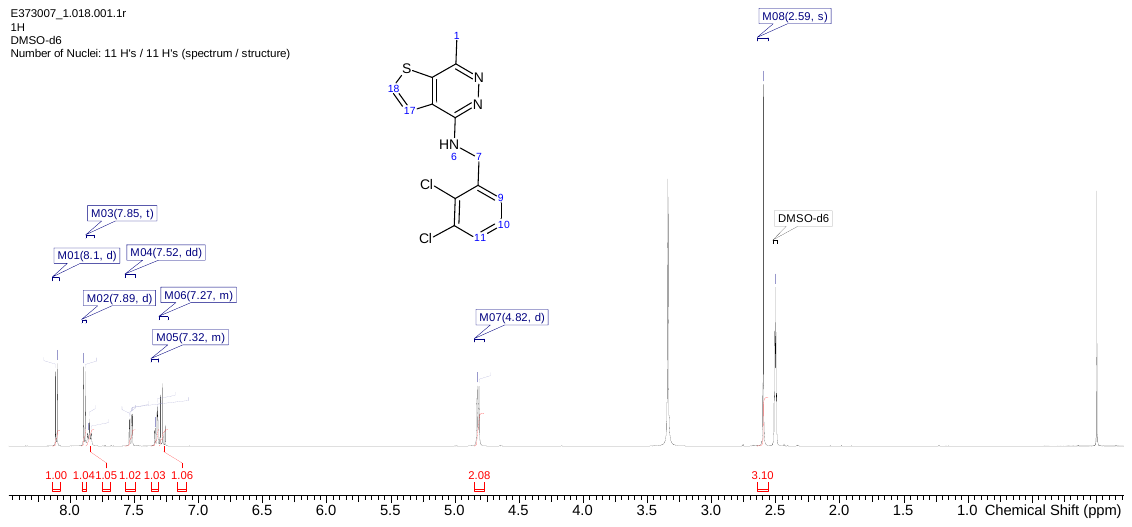

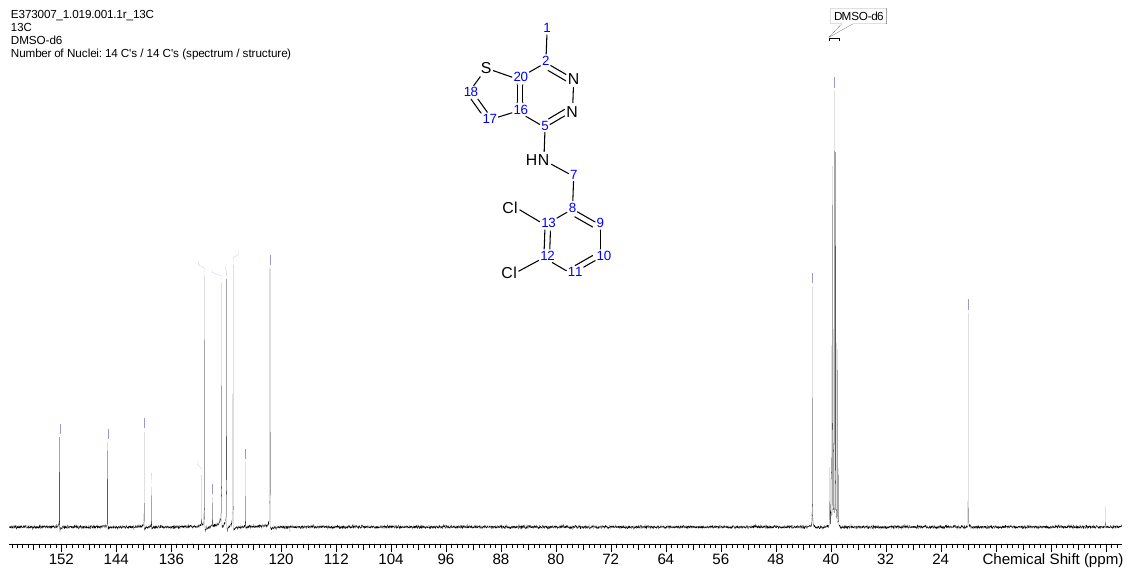

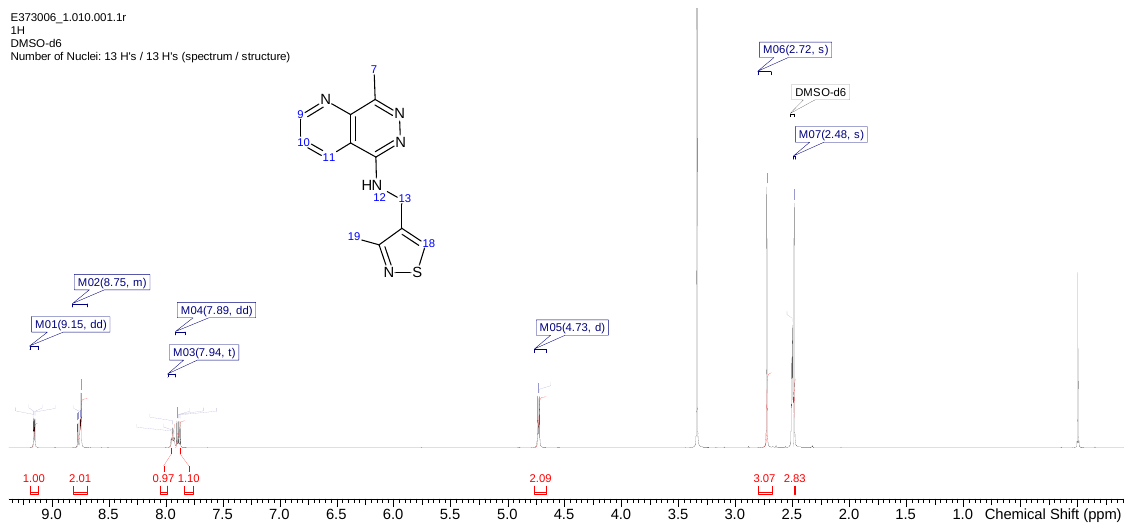

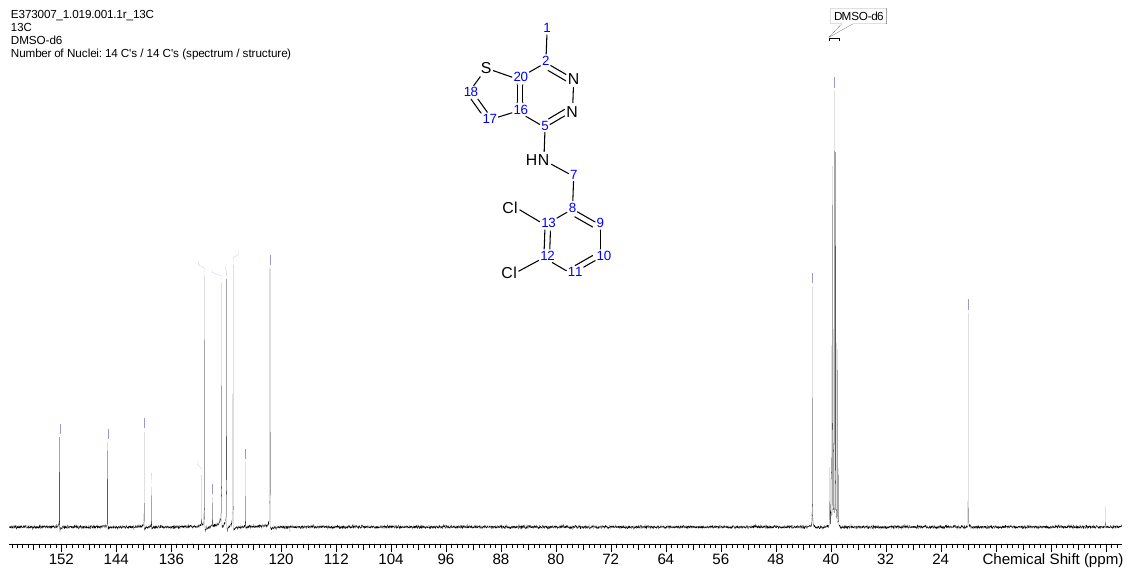

#### Fig. S30: NMR 1H, 13C and 19F spectra for JNJ-8833, JNJ-2842, JNJ-6887 and JNJ-1866.

Table S1: *In vitro* metabolic stability

| **Assay** | **JNJ-8833** | **JNJ-2842** | **JNJ-6887** | **JNJ-1866** |
| --- | --- | --- | --- | --- |
| **CL_int_ (Human)** (µL min mg) | 120 | 250 | 193 | <7.7 |
| **CL_int_ (Mouse)** (µL min mg) | >347 | >347 | >347 | 14.9 |

CL_int_: intrinsic clearance.

#### Table S2: Profiling of JNJ-6887 and JNJ-1866

| Condition | JNJ-6887 | JNJ-1866 |
| --- | --- | --- |
| *M. tuberculosis* H37Rv MIC_90_ (nM) | 50 | 360 |
| *M. tuberculosis* H37Rv MBC_99_ (µM) | 2 | 11 |
| *Mt*MenG enzyme activity IC_50_ (nM) | 1.3 | 12 |
| *M. tuberculosis* low pH media MIC_50_ (nM) | 620 | 1200 |
| *M. tuberculosis* 0.2% IC_50_ (µM) | >50 | >50 |
| *M. marinum MIC_90_* (nM) | 53 | 435 |

Table S3: ADME and in vitro toxicology profile for JNJ-6887 and JNJ-1866.

| **Parameter** | **JNJ-6887** | **JNJ-1866** |
| --- | --- | --- |
| hERG (hts) IC_50_ (µM) | >30.2 | >30.2 |
| CTCM | No hazard  identified up to 10 µM | No hazard  identified up to 10 µM |
| LM CL_int_, m/h (µL min mg^-1^) | >347/186.5 | 14.9/<7.7 |
| Hep CL_int_, m/h (µL min 10^6^ cells^-1^) | 119/19.5 | 70/4.4 |
| PPB, m/h (% bound) | 99.58/99.93 | 68.43/98.34 |
| MDCK AB+inhib, BA/AB | 19.1/2.08 | 23.4/10.2 |
| Eq. sol pH 2/7 (µM) | 579.5/0.23 | 473.75/401.35 |
| CHI log D pH 7.4 | 2.79 | 0.62 |
| Ames II | Negative | Negative |
| Cytotoxicity (HepG2 IC_20_ µM) | nd | >100 |
| Mitotoxicity (Glu/Gal IC_50_ µM) | >50/28.12 | >100/>100 |
| CYP450 inh IC50 (µM) (1A2/3A4/2C8/2C9/2C19/2D6) | >20/3.58/14.2/  14.7/16.2/2.9 | >20/>20/>20/  >20/>20/>20 |

nd: not determined.

Table S4: Comparison of dose-dependent pharmacokinetic profiles of JNJ-6887 and JNJ-1866

|  | **T_1/2_ (h)** | **C_max_ (ng mL^-1^)** | **AUC (h ng mL^-1^)** | **F(%)** |
| --- | --- | --- | --- | --- |
| **JNJ-6887** |  |  |  |  |
| 5 mg kg^-1^ | 3.19 | 125 | 333 | 17.5 |
| 30 mg kg^-1^ | 3.09 | 611 | 1850 | - |
| 100 mg kg^-1^ | 2.63 | 2338 | 7528 | - |
| **JNJ-1866** |  |  |  |  |
| 5 mg kg^-1^ | 1.3 | 943 | 1920 | 88.9 |
| 30 mg kg^-1^ | 3.51 | 3927 | 8854 | - |
| 150 mg kg^-1^ | 2.26 | 14667 | 92155 | - |

PO administration; F(%) = bioavailability, T_1/2_ = half-life, C_max_ = maximum concentration reached. AUC = area under the curve.

Table S5: Pharmacokinetic profiling of JNJ-1866

| **Parameter** | **JNJ-1866** | | | |
| --- | --- | --- | --- | --- |
| **Route** | **IV** | **PO** | **SC** | **SC** |
| Dose (mg kg^-1^) | 1 | 5 | 250 | 1250 |
| CL (mL min^-1^ kg^-1^) | 39.0 ± 5.3 | - | - | - |
| C_max_ (ng mL^-1^) | - | 943 ± 211 | 6470 | 11800 |
| t_1/2_ (h) | 0.247 | 1.30 ± 1.09 | 67 | 51.1 |
| T_max_ (h) | - | 0.5 | 1 | 4 |
| AUC_last_ (ng h mL^-1^) | 431 ± 55 | 1912 ± 221 | 141000 | 708000 |
| AUC_inf_ (ng h mL^-1^) | 432 ± 55 | 1920 ± 222 | 147000 | 710000 |
| F (%) | - | 88.9 ± 10.3 | >100 | >100 |
| Vss (L kg^-1^) | 0.585 ± 0.044 | - | - | - |
| Vz (L kg^-1^) | 1.27 ± 0.69 | - | - | - |
| MRT (h) | 0.243 ± 0.01 | 1.61 ± 0.16 | 133 | 206 |

Results are the mean ± s.d. n = 3 animals were dosed for the IV and PO arms and n = 3 animals by alternating sampling (from a total of 6 dosed) for SC arms (LAI), alternating sampling per time point was performed to limit the number of times blood was collected per animal, therefore we were unable to calculate error for these treatments. CL: clearance; C_max_ = maximum concentration reached; t_1/2_: half-life; AUC: area under the curve; MRT: Mean residence time; F: bioavailability; Vss: volume of distribution; Vz: terminal elimination phase.

Table S6: Drug susceptibility of JNJ-1866 and JNJ-6887 resistant strains against clinical compounds

| **Compound** | **WT** | **R6887-R2** | **R6887-R3** | **R1866-R1** | **R1866-R2** | **R1866-R3** |
| --- | --- | --- | --- | --- | --- | --- |
| Mutation |  | D25H | D25H | D25G | D25G | D25G |
| JNJ-8833 | 0.62 | >50 (80) | 44.0 (71) | 31.5 (50) | 31.3 (50) | 43.0 (69) |
| JNJ-2842 | 0.21 | >50 (236) | 32.4 (153) | 32.9 (155) | 37.0 (174) | 31.8 (150) |
| JNJ-6887 | 0.02 | 3.53 (211) | 1.52 (91) | 1.40 (83) | 1.19 (71) | 1.34 (80) |
| JNJ-1866 | 0.09 | 23.6 (265) | 20.8 (234) | 26.8 (302) | 22.9 (258) | 24.0 (271) |
| Bedaquiline | 0.10 | 0.03 (0.25) | 0.08 (1) | 0.09 (1) | 0.08 (1) | 0.09 (1) |
| Isoniazide | 0.25 | 0.13 (0.5) | 0.30 (1) | 0.29 (1) | 0.27 (1) | 0.28 (1) |
| Linezolid | 0.65 | 0.49 (1) | 0.59 (1) | 0.62 (1) | 0.62 (1) | 0.59 (1) |
| Rifampicin | 0.003 | 0.003 (1) | 0.003 (1) | 0.003 (1) | 0.003 (1) | 0.003 (1) |
| Ethambutol | 0.86 | 0.89 (1) | 0.98 (1) | 0.94 (1) | 0.91 (1) | 0.91 (1) |
| Ethionamide | 1.13 | 1.41 (1) | 1.49 (1) | 1.36 (1) | 1.46 (1) | 1.13 (1) |
| SQ109 | 1.55 | 1.44 (1) | 1.79 (1) | 1.98 (1) | 1.68 (1) | 2.11 (1) |
| Pretomanid | 0.10 | 0.09 (1) | 0.12 (1) | 0.11 (1) | 0.11 (1) | 0.12 (1) |
| Q203 | 0.003 | 0.0008 (0.33) | 0.003 (1) | 0.003 (1) | 0.003 (1) | 0.003 (1) |

MIC_50_ (µM) with fold difference from WT in brackets.

Table S7: Profiling of compounds against *M. tuberculosis* drug sensitive and resistant clinical isolates and other species

| **Strain** | **Lineage** | **Resistance** | **MIC_90_ (nM)** | | |
| --- | --- | --- | --- | --- | --- |
|  |  |  | **JNJ-6887** | **JNJ-1886** | **BDQ** |
| H37Rv | 4 | - | 33 | 322 | 239 |
| BDQR^Rv0678^ * | 4 | BDQ | 63 | 298 | 1231 |
| HN878 | 2 | - | 21 | 149 | 70 |
| Karonga-9271 | 4 | - | 97 | 957 | - |
| Karonga-7160 | 2 | - | 155 | 1073 | 190 |
| Karonga-6753 | 4 | - | 46 | 612 | 50 |
| Karonga-2703 | 3 | Rifampicin | 11 | 289 | 56 |
| Karonga-4779 | 2 | - | 8 | 167 | 31 |
| Karonga-7496 | 3 | - | 31 | 554 | 92 |
| Karonga-1781 | 2 | - | 21 | 189 | 26 |
| Karonga-1743 | 2 | - | 116 | 772 | 111 |
| Karonga-6787 | 4 | - | 53 | 450 | 153 |
| N0153 | 1 | - | 26 | 176 | 91 |
| N0155 | 2 | - | 25 | 231 | 46 |
| N1283 | 4 | - | 111 | 782 | 131 |
| N0145 | 2 | - | 37 | 317 | 70 |
| TB-TDR-0012 | 2 | MDR | 20 | 363 | 33 |
| TB-TDR-0154 | 3 | MDR | 22 | 510 | 19 |
| TB-TDR-0131 | 4 | MDR | 98 | 458 | 262 |
| 2013-02481^1,2^ | 3 | BDQ | 207 | 1345 | 1112 |
| 2014-03042^1,3^ |  | BDQ | 94 | 769 | 1492 |

MIC_90_ values of JNJ-6887, JNJ-1866 and bedaquiline (nM) against selected clinical strains taken from the Karonga and Gagneux collections (*25*). ^1^*In vitro* mutant selected for bedaquiline resistance (Rv0678: A65indel/frameshift) generated in this study. ^2^Alternative *in vitro* mutant with efflux-mediated (Rv0678: A344del/frameshift) resistance and fbiC-mediated (Rv1173: Arg536Leu) clofazimine resistance. ^3^*In vitro* mutant with MmpL5-mediated (Rv0676c: Ile948Val, Asn142Lys) and AtpE-mediated (Rv1305: Glu61Asp) resistance to bedaquiline. MIC_90_ of JNJ-6887 and JNJ-1866 within 5-fold of H37Rv WT strain (with exception of 2013-02481).

Table S8: Oligonucleotides used in this study

| **Name** | **Sequence** | **Notes** |
| --- | --- | --- |
| menH “low” CRISPRi_1 | GGGAGAACAACGCATTGGTGGGCGT |  |
| menH “low” CRISPRi_2 | AAACACGCCCACCAATGCGTTGTTC |  |
| menE “low” CRISPRi_1 | GGGAGGCGACCAGCGACGTGTATCGCCGG |  |
| menE “low” CRISPRi_2 | AAACCCGGCGATACACGTCGCTGGTCGCC |  |
| aroK-high CRISPRi fwd | GGGAACTTGCCGGAGCCCGGCAGGCCGA |  |
| aroK-high CRISPRi rvs | AAACTCGGCCTGCCGGGCTCCGGCAAGT |  |
| aroK-low CRISPRi fwd | GGGAATCGCGACGTCGGTGT |  |
| aroK-low CRISPRi rvs | AAACACACCGACGTCGCGAT |  |
| aroG-high CRISPRi fwd | GGGAGTAGATTGCGGTCGGCCACCCCA |  |
| aroG-high CRISPRi rvs | AAACTGGGGTGGCCGACCGCAATCTAC |  |
| aroG-low CRISPRi fwd | GGGAGTAGATTGCGGTCGGCCACCCCA |  |
| aroG-low CRISPRi rvs | AAACTGGGGTGGCCGACCGCAATCTAC |  |
| qPCR_menE_f | CCACCACCAACGCTAGAAG |  |
| qPCR_menE_r | GGTAGCACGTTCACGACAT |  |
| qPCR_menG_f | TGCGGTCACCATCAGTTTC |  |
| qPCR_menG_r | GAGAATTCGCACACTAGTAGCC |  |
| aroK qPCR fwd | CGTCGTCTACCTGGAGATCA |  |
| aroK qPCR rvs | ATCAGCGCGCGGTATTT |  |
| aroG qPCR fwd | CTACAGACCGCCGAAATCTATG |  |
| aroG qPCR rvs | GTGCGGACAGGTCAAACA |  |
| menG-C146R OE fwd | GCTACTAGTGCGCGAATTC | Site direct mutagenesis primer |
| menG-C146R OE rvs | CGCCCGCCCGGCCGGGTGA | Site direct mutagenesis primer |
| qPCR_sigA_f | GACGAAGACCACGAAGACC |  |
| qPCR_sigA_r | CATCCCAGACGAAATCACC |  |
| CV010 | ATGGCGACCACAACCAGG | Taken from Andries *et al.* (2014) (*2*) |
| CV017 | TTTTACGCGTGTTGCTCATCAGTCGTCCTC | Taken from Andries *et al.* (2014) (*2*) |
| AtpE_F | TGTACTTCAGCCAAGCGATGG | Taken from Huitric *et al.* (2010) (*69*) |
| AtpE-R | CCGTTGGGAATGAGGAAGTTG | Taken from Huitric *et al.* (2010) (*69*) |
| pepQ-F | ATCAATGCCCCCTGGAAC | Taken from Yang *et al.* (2020) (*70*) |
| pepQ-R | GCACGTTCTTCAACTTGGTG | Taken from Yang *et al.* (2020) (*70*) |
| MenG FP1.1 | TGATCTGATAACCCCGCACC | PCR amplification and Sanger Sequencing |
| MenG RP1.1 | TACGCAGCCCGAAACTGATG | PCR amplification and Sanger Sequencing |
| MenG FP1.2 | ACCAATACCGTGTTGTCCC | PCR amplification and Sanger Sequencing |
| MenG RP1.2 | ACCCGCTAAGTCTTCCTACC | PCR amplification and Sanger Sequencing |
| qPCR_Rv0678_F | [GAACAGATGGGCGGCTATT](https://protect-eu.mimecast.com/s/pNpZCoy14uprZ9yhOpkj6?domain=thermofisher.com) |  |
| qPCR_Rv0678_R | [CGCTCGGGATCACACAC](https://protect-eu.mimecast.com/s/pNpZCoy14uprZ9yhOpkj6?domain=thermofisher.com) |  |
| qPCR_MmpS5_F | [GATCCGCACTTTCTTTGGTTC](https://protect-eu.mimecast.com/s/pNpZCoy14uprZ9yhOpkj6?domain=thermofisher.com) |  |
| qPCR_MmpS5_R | [AGCCGGAAACTTCGTACTC](https://protect-eu.mimecast.com/s/pNpZCoy14uprZ9yhOpkj6?domain=thermofisher.com) |  |
| qPCR_MmpL5_F | ACTACAACGACCGCAACTAC |  |
| qPCR_MmpL5_R | [TTTCGACCATCAGCACCTC](https://protect-eu.mimecast.com/s/pNpZCoy14uprZ9yhOpkj6?domain=thermofisher.com) |  |
| qPCR_pepQ_F | CTTCGAGAGCCACGTGGTC |  |
| qPCR_pepQ_R | GCAGTGACTCCACAGTTCCG |  |
| qPCR_Rv1979c_F | GCGTGCTGTTGGCCATCAAC |  |
| qPCR_Rv1979c_R | GCTGGGTGGTGATGATCCAC |  |
| qPCR_atpE_F | GTAACGCGCTTATCTCCGGT |  |
| qPCR_atpE_R | ACTTGACGGGTGTAGCGAAG |  |
| MMAR-1007-F | GCGCATGAAGTTGTTGTAGAC | *M. marinum* sequencing primer |
| MMAR-1007-R | TCACCTTCGACAAGATCGATC | *M. marinum* sequencing primer |

Movie S1.

Timelapse image sequence of *M. tuberculosis* treated with JNJ-6887 - *M. tuberculosis* expressing TdTomato was cultured in a microfluidic device in standard 7H9 broth (0-75 h) and then exposed to 1 µM JNJ-6887 (76-267 h) and 7H9 broth (drug washout, 268-336 h). Images were acquired every 1 h on the phase and red channels (Ex 555/Em 590 nm). Media conditions are depicted on top left and hours elapsed on bottom left. The numbers 1, 2, and 3 represent three different xy positions from the experiment and were combined in FIJI. Scale bar is shown on the bottom right.

### References

2.	Andries, K. et al. Acquired Resistance of Mycobacterium tuberculosis to Bedaquiline. PLOS ONE 9, e102135, doi:10.1371/journal.pone.0102135 (2014).

26.	Bosch, B. et al. Genome-wide gene expression tuning reveals diverse vulnerabilities of M. tuberculosis. Cell 184, 4579-4592.e4524, doi:10.1016/j.cell.2021.06.033 (2021).

29.	Phelan, J. et al. An open-access dashboard to interrogate the genetic diversity of Mycobacterium tuberculosis clinical isolates. Scientific Reports 14, 24792, doi:10.1038/s41598-024-75818-y (2024).

30.	Dai, Y. N. et al. Crystal structures and catalytic mechanism of the C-methyltransferase Coq5 provide insights into a key step of the yeast coenzyme Q synthesis pathway. Acta Crystallogr D Biol Crystallogr 70, 2085-2092, doi:10.1107/s1399004714011559 (2014).

31.	Jumper, J. et al. Highly accurate protein structure prediction with AlphaFold. Nature 596, 583-589, doi:10.1038/s41586-021-03819-2 (2021).

34.	Upadhyay, A. et al. Mycobacterial MenJ: An Oxidoreductase Involved in Menaquinone Biosynthesis. ACS Chemical Biology 13, 2498-2507, doi:10.1021/acschembio.8b00402 (2018).

35.	Dunovska, K., Klapkova, E., Sopko, B., Cepova, J. & Prusa, R. LC-MS/MS quantitative analysis of phylloquinone, menaquinone-4 and menaquinone-7 in the human serum of a healthy population. PeerJ 7, e7695, doi:10.7717/peerj.7695 (2019).

38.	Sukheja, P. et al. A Novel Small-Molecule Inhibitor of the Mycobacterium tuberculosis Demethylmenaquinone Methyltransferase MenG Is Bactericidal to Both Growing and Nutritionally Deprived Persister Cells. mBio 8, 10.1128/mbio.02022-02016, doi:10.1128/mbio.02022-16 (2017).

45.	Lamprecht, D. et al. Targeting de novo purine biosynthesis for tuberculosis treatment. Nature 644 (8075), 214-220, doi:10.1038/s41586-025-09177-7 (2025).

58.	Cole, S. T. et al. Deciphering the biology of Mycobacterium tuberculosis from the complete genome sequence. Nature 393, 537-544, doi:10.1038/31159 (1998).

59.	Manina, G. & Dhar, N. Single-Cell Analysis of Mycobacteria Using Microfluidics and Time-Lapse Microscopy. Methods Mol Biol 2314, 205-229, doi:10.1007/978-1-0716-1460-0_8 (2021).

60.	Schindelin, J. et al. Fiji: an open-source platform for biological-image analysis. Nat Methods 9, 676-682, doi:10.1038/nmeth.2019 (2012).

61.	Schmittgen, T. D. & Livak, K. J. Analyzing real-time PCR data by the comparative C(T) method. Nat Protoc 3, 1101-1108, doi:10.1038/nprot.2008.73 (2008).

62.	Berg, K. et al. SAR study of piperidine derivatives as inhibitors of 1,4-dihydroxy-2-naphthoate isoprenyltransferase (MenA) from Mycobacterium tuberculosis. Eur J Med Chem 249, 115125, doi:10.1016/j.ejmech.2023.115125 (2023).

63.	Lamprecht, D. A. et al. Turning the respiratory flexibility of Mycobacterium tuberculosis against itself. Nature Communications 7, 12393, doi:10.1038/ncomms12393 (2016).

64.	Kingdon, A. D. H., Meosa-John, A.-R., Batt, S. M. & Besra, G. S. Vanoxerine kills mycobacteria through membrane depolarization and efflux inhibition. Frontiers in Microbiology Volume 14 - 2023, doi:10.3389/fmicb.2023.1112491 (2023).

65.	Matty, M. A., Oehlers, S. H. & Tobin, D. M. Live Imaging of Host-Pathogen Interactions in Zebrafish Larvae. Methods Mol Biol 1451, 207-223, doi:10.1007/978-1-4939-3771-4_14 (2016).

66.	Takaki, K., Davis, J. M., Winglee, K. & Ramakrishnan, L. Evaluation of the pathogenesis and treatment of Mycobacterium marinum infection in zebrafish. Nature Protocols 8, 1114-1124, doi:10.1038/nprot.2013.068 (2013).

67.	Lu, H. R. et al. Identifying Acute Cardiac Hazard in Early Drug Discovery Using a Calcium Transient High-Throughput Assay in Human-Induced Pluripotent Stem Cell-Derived Cardiomyocytes. Frontiers in Physiology 13, doi:10.3389/fphys.2022.838435 (2022).

68.	Sharma, P. et al. Evolution of Small Molecule Inhibitors of Mycobacterium tuberculosis Menaquinone Biosynthesis. Journal of Medicinal Chemistry 68, 5774-5803, doi:10.1021/acs.jmedchem.4c03156 (2025).

69.	Huitric, E. et al. Rates and mechanisms of resistance development in Mycobacterium tuberculosis to a novel diarylquinoline ATP synthase inhibitor. Antimicrob Agents Chemother 54, 1022-1028, doi:10.1128/aac.01611-09 (2010).

70.	Yang, J. et al. Molecular characteristics and in vitro susceptibility to bedaquiline of Mycobacterium tuberculosis isolates circulating in Shaanxi, China. International Journal of Infectious Diseases 99, 163-170, doi:10.1016/j.ijid.2020.07.044 (2020).
